# Altered chromatin accessibility and nucleosome positioning landscape upon HDAC and LSD1 inhibition in cancer cell

**DOI:** 10.64898/2026.04.08.717275

**Authors:** Sagnik Sen, Pierre O. Estève, Devesh Tarasia, Rachel Danneberg, Ashmita Dey, Ujjwal Maulik, Sanghamitra Bandyopadhyay, Sriharsa Pradhan

**Author notes:** Corresponding authors: Sagnik Sen: Sriharsa Pradhan. These authors contributed equally (Sagnik Sen, Pierre O. Estève, Devesh Tarasia). This author supervised this work Sriharsa Pradhan, Ph.D. New England Biolabs Ipswich, MA 01983, USA.

## Abstract

Epigenetic enzymes, writers, readers and erasers regulate chromatin landscapes and participate in tumor heterogeneity. While therapeutic targeting of these enzymes has shown clinical promise, the comparative efficacy of mono-versus dual-inhibitor strategies remain unclear. Here, we introduce a multi-modal platform that uses NicE-viewSeq and integrates automated deep learning based spatially resolved chromatin accessibility profiling with high-throughput sequencing following epigenetic inhibitor application. Accessible chromatin landscapes were altered along with nucleosome positioning following inhibition of either LSD1 or HDACs alone, or both together. Coordinated modulation of histone marks and the CoREST complex on chromatin was observed across inhibitory conditions. Transcription factor binding analysis identified three predominant families, ETS, RUNT, and bZIP with enhanced chromatin association upon treatments. Mechanistically, a CoREST-RUNX regulatory axis was uncovered wherein JunB, a member of bZIP family displaces CoREST-RUNX at differentially accessible regions, triggering apoptotic pathways. Therefore, JunB-mediated mechanism reveals a convergent therapeutic vulnerability, offering new avenues for optimizing different combinatorial epigenetic therapy in cancer.

## Introduction

Epigenetic modifications play a crucial role in regulating the genomic landscape and maintaining tumor heterogeneity in cancer. The roles of histone lysine acetylation and methylation, and DNA cytosine methylation in tumor progression are largely explored, suggesting DNA hypermethylation and H3K9me3 at the tumor suppressor gene promoters to be one of the mechanisms of gene silencing (Shvedunova et al. 2022, Sasidharan Nair et al. 2018, Easwaran et al. 2014, Ohm et al., 2007, Kazanets et al. 2016). On the contrary, DNA hypomethylation of oncogene promoters, along with extensive demethylation at the repetitive elements, can facilitate aberrant gene expression, chromatin instability, and tumor development (Van et al. 2017, Ehrlich 2009). Indeed, The Cancer Genome Atlas (TCGA) has provided comprehensive cancer-specific genomic, epigenomic, and transcriptomic profiles across ∼33 matched cancer types and subtypes, offering insights into molecular events in tumor biology (Hoadley et al. 2018). Notably, TCGA data sets have confirmed that aberrant DNA methylation is associated with cancer (Liang et al. 2023). Previous studies underscore the importance of DNA methylation in shaping silent heterochromatin, leading to tumor specificity (Grewal 2023). This has led to the specification of the chromatin state and its dynamic accessibility as a tumor variability marker.

Over the past decade, the advent of single-cell RNA sequencing (scRNA-seq) has marked a breakthrough in personalized medicine, enabling deeper resolution of tumor heterogeneity. Furthermore, advancements in high-throughput sequencing assays for chromatin accessibility and transcription factor occupancy have resulted in enhanced data integration at single-and bulk cells in multi-omics studies, facilitating a more comprehensive understanding of cancer epigenetics. Additionally, spatial profiling of tissue samples has revolutionized 3D visualization and contextual analysis of genes in cells. Indeed, spatial ATAC-seq has set a benchmark for tissue and cellular identification profiling (Deng et al. 2022). However, its reliance on transposase activity poses challenges and has limited success with chemically fixed archival tissue samples. NicE-viewSeq addressed these limitations since it relies primarily on formaldehyde fixed cell samples. This assay employs dual labeling with one fluorescent dNTP and biotinylated dATP, enabling multimodal outputs consisting of microscopic imaging with fluorescent dNTP incorporation and high-throughput sequencing of biotinylated accessible DNA. This multimodality enables the study of chromatin accessibility dynamics, providing insights into their cellular localization. This comprehensive approach can elucidate visual and structural chromatin accessibility modulation in disease conditions and allows epigenetic therapeutic intervention.

Numerous studies have demonstrated the importance of therapeutic strategies aiming at epigenetic modifications by targeting enzymes (writers and erasers) and the binding proteins (readers) for cancers. HDAC inhibitors (Vorinostat, Belinostat), and DNMT1 inhibitors (Vidaza, Decitabine) are clinically prescribed for blood-related cancers (Duvic et al. 2007., Cao et al., 2023, Oran et al., 2018, Fili et al., 2019). In addition, a list of inhibitors is currently on clinical trials (Yu et al. 2024). Apart from traditional mono-target specific inhibitor, dual-inhibiting therapeutic agents targeting HDAC complexes (15a for DNMT1-HDACi; corin for LSD1-HDACi), have shown significantly high antitumor effects by inducing apoptosis (Huang et al. 2024, Kalin et al. 2018). Corin combines the pharmacophores of clinically used HDAC inhibitor and a preclinical LSD1 inhibitor to generate dual action of HDAC-LSD1 inhibitor, thus targeting the HDAC-LSD1 complex (CoREST). It shows substantially higher efficacy over the parental HDAC1 inhibitor (Entinostat), in malignancies, particularly with squamous cell carcinoma and melanoma (Marques et al. 2020). Indeed, combination of epigenetic drugs with other treatment modalities, such as chemotherapy or immunotherapy, shows potential in enhancement of efficacy and reducing drug resistance, especially in cancer (Song et al. 2025).

While the therapeutic efficacy of dual inhibitors over mono-HDAC inhibitors has been studied on few distinct cancer types, comparative understanding of the molecular and cellular changes, along with mechanistic differences, is lacking. To further investigate the mechanistic differences underlying chromatin regulation, we developed a pipeline to perform a comparative chromatin accessibility analysis in a single inhibitor treatment of HDACi or LSD1i, or a combination of both for multi-target inhibition. This study focuses on evaluating their relative efficacy in modulating chromatin accessibility alteration by leveraging the NicE-viewSeq multi-modality incorporating NicEL (NicE-view Learning), an integrated framework that combines a modified U-Net architecture for nuclei detection with qualitative chromatin accessibility assessment and high-throughput sequencing to enable comprehensive analysis of chromatin modifications post epigenetic inhibitor treatment in model cancer cell lines.

## Results

### NicEL-a NicE-viewSeq based deep learning pipeline for chromatin accessibility quantification

NicE-viewSeq is an integrative visualization and genomics method to detect accessible chromatin encompassing accessible promoters, enhancers, nucleosome positioning, transcription factor occupancy, and other chromosomal protein binding in fixed mammalian cells (Esteve et al. 2020). We used this method to develop the NicEL model. NicEL is an ensemble model consisting of two stages: 1. nuclei detection model, 2. qualitative measurement of chromatin accessibility (Fig. 1A). In the first stage, we applied a U-Net model for nuclei detection, treating it as a cell nuclei segmentation task (Fig. 1A ii-iv). Cell nuclei segmentation is a well-known open problem and one of U-Net’s applications (Vijayan et al. 2024). For NicE-view images, the nuclei detection task is particularly crucial for successful execution of subsequent steps, since random noise in the color channels may introduce artifacts.

**Figure 1.**
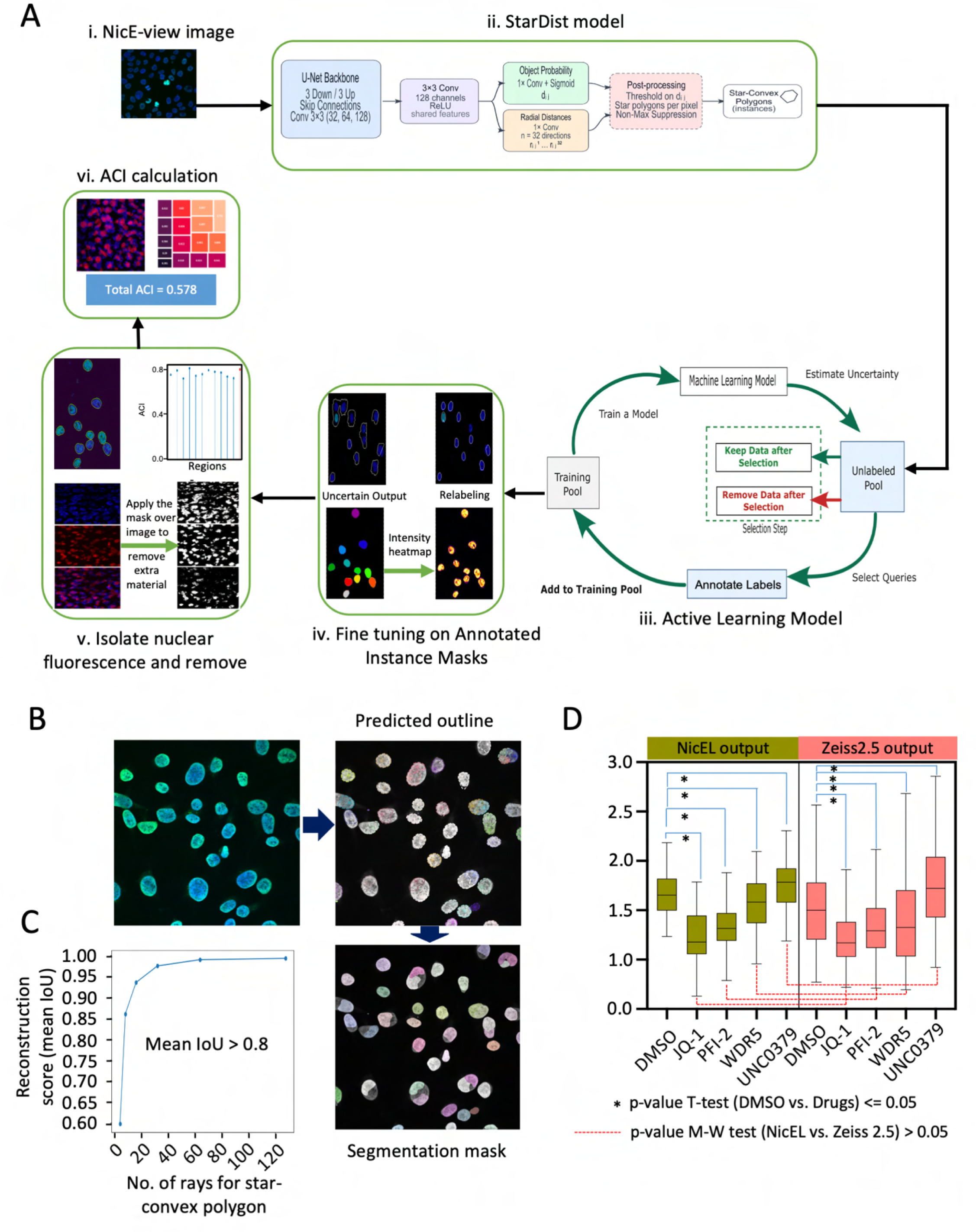
NicEL (NicE-view Learning model) precisely quantifies chromatin accessibility. **A.** A schematic diagram of NicEL stages (i-vi). **B.** Various stages of NicEL for detecting the segmentation mask of NicE-view images. **C.** Reconstruction score (mean IoU) showing a preferable range of star-convex polygon size. **D.** Comparative analysis to validate ACI scores generated by NicEL (Asparagus Green) using manually curated images (Salmon Red) from JQ-1, PFI-2, WDR5, UNC0379 therapeutic agents and DMSO control using box plots is shown. Asterisk (*) displayed statistical significance in ACI compared to DMSO calculated using ‘two-sample T-test’. Dotted lines (red line) showed acceptance of H_0_ hypothesis (calculated using Mann Whitney Rank Sum test), validating NicEL generated ACI scores.

Following nuclei detection, the next step involves segregating color channels to assess chromatin accessibility (Fig. 1A v). Typically, accessible chromatin regions are labeled with fluorescent dyes by fluorescent-conjugated dNTP incorporation, such as fluorescent green (Cy-3) or red (Texas Red), for readout. The intensity of these color channels serves as a proxy for chromatin accessibility, with higher intensity indicating greater accessibility and lower intensity representing reduced accessibility respectively. However, raw fluorescence images may exhibit variations in intensity due to inconsistencies in fluorescent labelling and imaging, leading to potential misinterpretation of accessibility levels. To mitigate this issue, we introduced a scoring metric termed the Accessible Chromatin Index (ACI, materials and methods), which normalizes accessibility measurements by computing the ratio of two channels; maximal intensity of chromatin accessibility, normalized to minimal intensity of the DNA label (DAPI, Fig. 1A vi). This approach ensures a standardized measure of chromatin accessibility across different conditions.

Application of ACI to NicE-view images produced high-quality nuclear segmentation masks across diverse imaging conditions, including densely packed nuclei (Fig. 1B, C). These masks enabled reliable extraction of nuclear fluorescence signals for chromatin accessibility analysis, where the nuclei boundary was determined based on convex polygons with IoU (area of intersection/area of union) > 0.8 (Fig. 1C). Raw ACR (accessible chromatin region) fluorescence intensities showed substantial inter-image variability, reflecting differences in staining and imaging conditions. Normalization using the ACI markedly reduced this variability, yielding tighter and more comparable accessibility distributions across samples as observed between NicEL output vs. Zeiss 2.5 output (Fig. 1D; Table S1). To assess NicEL performance in detail, we conducted a comparative analysis of the ACI derived from NicEL-based nuclei detection and manually selected nuclei using Zeiss 2.0 software. This analysis was performed across four inhibitor treatment conditions. JQ-1, PFI-2, WDR5, and UNC0379 along with DMSO as the control (Fig. 1D). Both JQ-1 and PFI-2 are H3K4 methyltransferase SETD7 inhibitors and WDR5 inhibitor targets the WD repeat domain 5 protein, a scaffolding component in histone methyltransferase complexes (like MLL/SET complexes) responsible for adding activating H3K4 methylation (Zhou et al. 2020, Barsyte-Lovejoy et al. 2014, Ruthenburg et al 2006). Therefore, these three inhibitors (JQ-1, PFI-2, and WDR5) are associated with reduced chromatin accessibility. However, UNC0379 selectively inhibits H3K9me2 methyltransferase G9a, thus it is expected to keep the chromatin in a more accessible state (Julien et al. 2020). For each condition, we analyzed 100 images. The ACI values obtained from both manual using Zen2.0 Zeiss software and NicEL-based detection showed strong concordance with expected drug effects. While the two-sample t-test, compared to DMSO, exhibited a significant difference in chromatin accessibility for all treatment conditions, the Mann-Whitney Rank-Sum test revealed no significant differences between both manually calculated ACI and NicEL-derived ACI. However, the distribution of ACI values from NicEL-based detection exhibited greater consistency with low standard deviation compared to manual detection (Fig. 1D). These findings confirm the accuracy and robustness of the NicEL model in detecting chromatin accessibility changes with reduced variability.

### NicE-seq multi-modality unveiled the differences between HDAC and LSD1 inhibitor

After the successful development of NicEL, we applied it to determine the mechanistic differences between therapeutic mono-and dual-inhibition. Two well-known therapeutic agents GSK-2879552 (LSD1 inhibitor (LSD1i)) and romidepsin (HDAC inhibitor (HDACi)), and their combinations for dual enzyme inhibition were chosen for the study. We treated HT1080 cells with LSD1i, HDACi or both LSD1i and HDACi, using DMSO as the control. For mono-inhibitory conditions, cells were treated with 5μM of GSK-2879552 and 1μM of romidepsin for 2 days and 6 hours, respectively. Due to the higher toxicity of romidepsin (Prince et al. 2020), dual-inhibitory cell populations were treated with 5μM GSK-2879552 for two days, followed by 6 hours of 1μM romidepsin. The treated and control cells were utilized for chromatin accessibility profiling using multi-modal NicE-seq. As the treatments were aimed at inhibiting mostly LSD1, a H3K4me1/me2 demethylase, and class I and II HDACs, we hypothesized an increase in chromatin accessibility post-treatment. Similarly, dual inhibitor treated cells would demonstrate an additive effect on chromatin accessibility. To analyze the changes, we acquired 100 images for each treatment to monitor accessibility after applying the segmentation mask (Fig. 2A, left panel). Identified nuclei were used for ACI calculations. Additionally, we performed high-throughput chromatin accessibility sequencing using NicE-seq of five experimental replicates for each treated condition.

**Figure 2.**
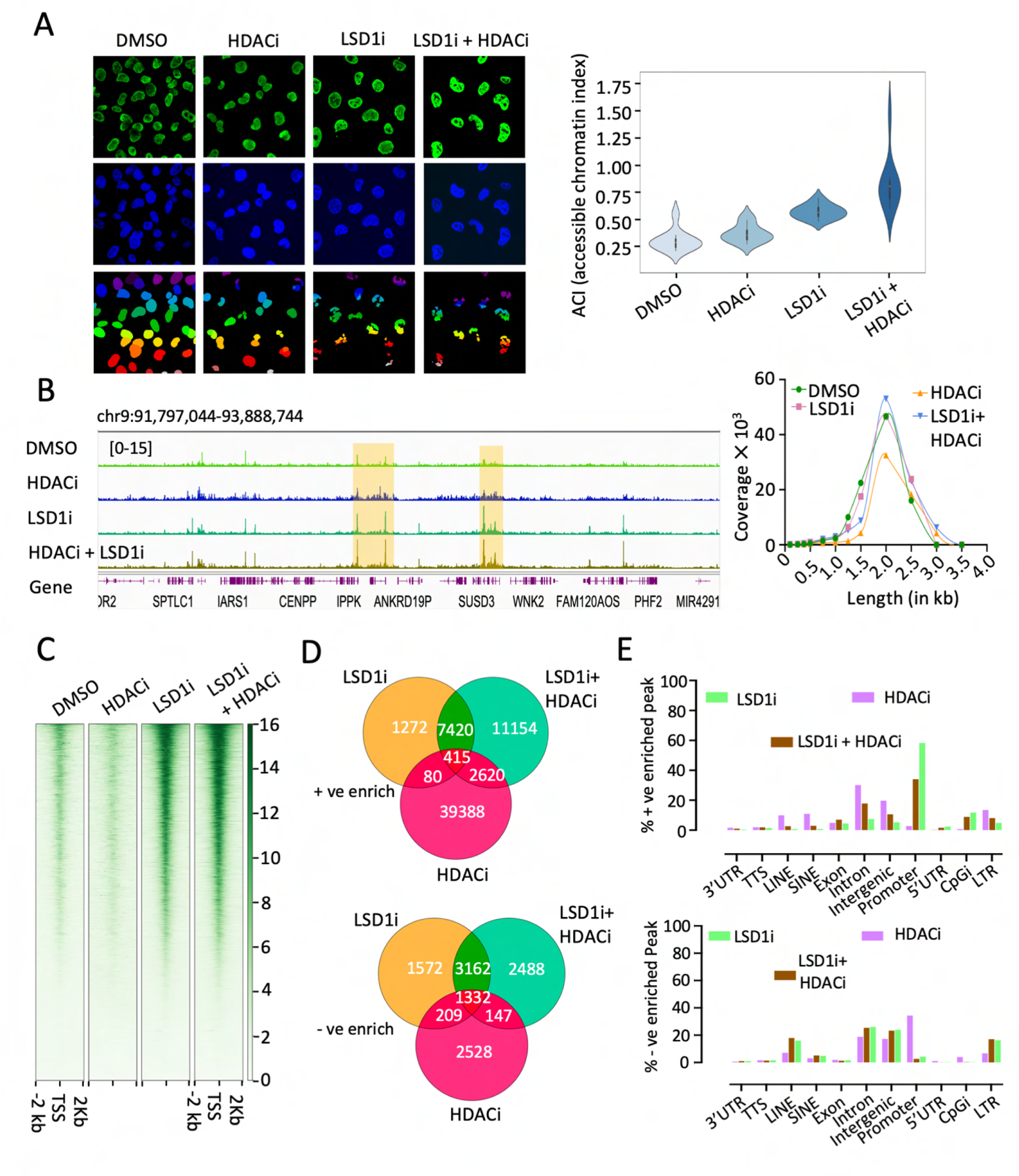
NicE-seq multi-modality reveals distinct chromatin accessibility profiles. **A.** NicE-view images from cells treated with LSD1i, HDACi, and DMSO as control with respective segmentation mask (left panel). Violin plots displaying the distribution of Accessible Chromatin Index (ACI) calculated from 100 images (right panel). **B.** The changes in accessible chromatin profile over the inhibitor treatments are highlighted (yellow) in Integrative Genome Viewer (IGV) (left panel). Coverage vs bin plots displayed genome-wide redistribution of accessibility over the treatments (right panel). **C.** Enrichment in chromatin accessibility upon LSD1i and LSD1i plus HDACi treatment at TSS ± 2kb is shown using heatmaps. **D.** Venn diagrams showed co-localization between positive enriched (+ve, upper panel) and negative enriched (-ve, lower panel) differential peaks for LSD1i, HDACi and LSD1i plus HDACi. **E.** Annotation of +ve enriched (upper panel) and-ve enriched (lower panel) differential peaks displaying genome wide distribution.

Indeed, the fluorescence intensities in treated cells were higher compared to DMSO, with combination of LSD1i and HDACi demonstrating the maximum increase in intensities (Fig. 2A). The ACI scores for mono-inhibition (i.e., LSD1i or HDACi cells) exhibited on average ∼0.3-0.5x higher ACI values compared to the DMSO control, while the LSD1i plus HDACi treated cells demonstrated ∼2x higher ACI over DMSO. (Fig. 2A, right panel). Indeed, the chromatin accessibility trend for all treated cells was higher than DMSO control. Corroborating with the images, NicE-seq of these treated cell samples also supported a similar trend of increased chromatin accessibility genome-wide compared to DMSO in IGV (Fig. 2B left panel). The HDACi and HDACi plus LSD1i treated accessible chromatin length increased genome-wide from 3.0-3.25 and 3.0-3.5 kb respectively, suggesting spreading of accessible chromatin regions in response to both inhibitors. Additionally, at 2 kb, the coverage was highest for LSD1i and HDACi combination (Fig. 2B, right panel). The metagenome heatmap exhibited substantially higher enrichment for LSD1i and LSD1i plus HDACi at ± 2 kb of TSS (Fig. 2C). This was in contrast with chromatin accessibility enrichment in HDACi treated cells, where enrichments were outside of TSS and displayed a diffused pattern (Fig. 2C, Fig. S1A). Next, we performed differential peak analysis to identify maximally impacted regions (peaks) across different inhibitor treated conditions. We observed a gradual increase in the number of differential peaks for LSD1i (∼15k), LSD1i plus HDACi (∼28k), and HDACi (∼46k) treated cells compared to DMSO, where ∼60%, ∼75%, and 91% of peaks were positively enriched, respectively (Fig. S1B). Higher percentage of positively enriched peaks in HDACi compared to HDAC1 plus LSD1i treated cells showed lack of synergistic effect. This may be due to an increase in accessible chromatin length following both inhibitor treatment leading to spreading of accessibility and decreasing the peak numbers (Fig. S1B). However, |log_2_FC| of differential peaks was ∼1.5-2x higher in LSD1i and LSD1i plus HDACi cells compared to HDACi alone as observed in MA-plots, (Fig. S1C-E) demonstrating a synergistic effect in accessible chromatin formation following combinatorial inhibition. Taken together, there could be a possibility of decrease in accessible peak numbers in dual inhibitor application due to an increase in accessible region width (Fig. 2B and Fig. S1C-E).

Since there was specificity in the genome-wide accessibility distribution among different inhibitor treatment conditions, we next examined the degree of overlap amongst the positively and negatively enriched chromatin accessibility peaks to study specificity. The Venn diagram within positively enriched accessible peaks confirmed the extended effect with differential peaks, where most peaks from HDACi cells were unique (∼39k) and a significant number of peaks were shared between LSD1i and LSD1i plus HDACi (7.4K; Fig. 2D, upper panel). However, the Venn diagram within negatively enriched accessible peaks demonstrated higher number of peaks colocalization among the three conditions 1.3 K (Fig. 2D, lower panel). Next, we annotated positively and negatively enriched accessible peaks using Homer to observe their genomic localization. Indeed, positively enriched accessible peaks from LSD1i and LSD1i plus HDACi treatment were strongly localized in the promoter regions of genes confirming TSS specific enrichment (Fig. 2E and 2C). Whereas in HDACi treated samples, the accessible peaks were mostly localized in gene-body regions, in the negatively enriched accessible peaks, a higher percentage of accessible chromatin was found in promoter regions following HDACi. In the same set of data, LSD1i, LSD1i and HDACi accessible chromatin was localized mostly in the gene-body, including introns and intrageneic regions (Fig. 2E).

Overall, both imaging and sequencing exhibited a similar trend, which was further supported by density plots on |log_2_FC| from ACI and differential peaks along with mean ± SD profile for LSD1i (Fig. S1F), LSD1i plus HDACi (Fig. S1G) and HDACi (Fig. S1H). However, HDACi cells displayed a lower accessible chromatin density overlap due to redistribution. Additionally, the violin plots and K-S test values confirmed no significant difference between both imaging and sequencing (Fig. S1I-K). Together, these results confirmed that the distinct chromatin accessibility profiles across different treatment conditions are specific to the drugs. Furthermore, we show that the chromatin accessibility changes from both modal outputs of NicE-view ACI and NicE-seq coverage is highly convergent.

### Altered chromatin accessibility and nucleosome positioning by HDAC1 and LSD1 inhibition

Since LSD1 and HDAC inhibition perturb H3K4me1, me2 and H3 lysine acetylation, we analyzed genome-wide histone H3 profiles by performing NEED-seq (Sen et al. 2025) for histone H3, H3K4me1, H3K4me2, H3K9ac and comparing with chromatin accessibility following LSD1i and HDACi exposure (Fig. 3A). These three histone H3 marks are associated with active genes, and IGV showed significant overlap between them and accessible chromatin, as expected. Interestingly, histone H3 coverage within genome-wide accessible peaks followed a pattern of spreading, where the sharp narrow peaks in DMSO control conditions started to distribute throughout the genome concurrent with significant loss of read coverage in the presence of inhibitors (Fig. 3B). The histone modification in coverage profile was highly systematic, where LSD1i demonstrated a sharp shift towards 2.5 kb, followed by spreading towards 3.5 and 4 kb bin size. However, in LSD1i and LSD1i plus HDACi treated samples, a significant loss in read coverage was observed. Drastic coverage loss along with profile spreading up to the 4 kb with HDACi exposure highly complied with its role in massive accessible chromatin spreading (Fig. 3B). Likewise, we also analyzed read coverage distributions of various H3 lysine modifications representing LSD1 and HDAC activities. Indeed, H3K4me1, me2 and H3K9ac mirrored histone H3 pattern observed in Fig. 3B, with significant shift in their distribution (∼1.5 kb) post-inhibitor treatment conditions (Fig. S2A-C). H3K4me1, the precursor for H3K4me2, coverage was significantly lowered in the same experimental conditions, suggesting the mark is possibly converted to H3K4me2 by enzymes (Fig. S2A-C). Surprisingly, we observed H3K9ac enrichment on accessible chromatin was highly affected by LSD1 inhibition (Fig. S2A). Similarly, HDAC1 inhibition resulted in lower H3K4me2 enrichment on accessible chromatin (Fig. S2C). These data suggest LSD1 or HDAC inhibition leads to spreading of H3K9ac and H3K4me2 occupancy on accessible chromatin.

**Figure 3.**
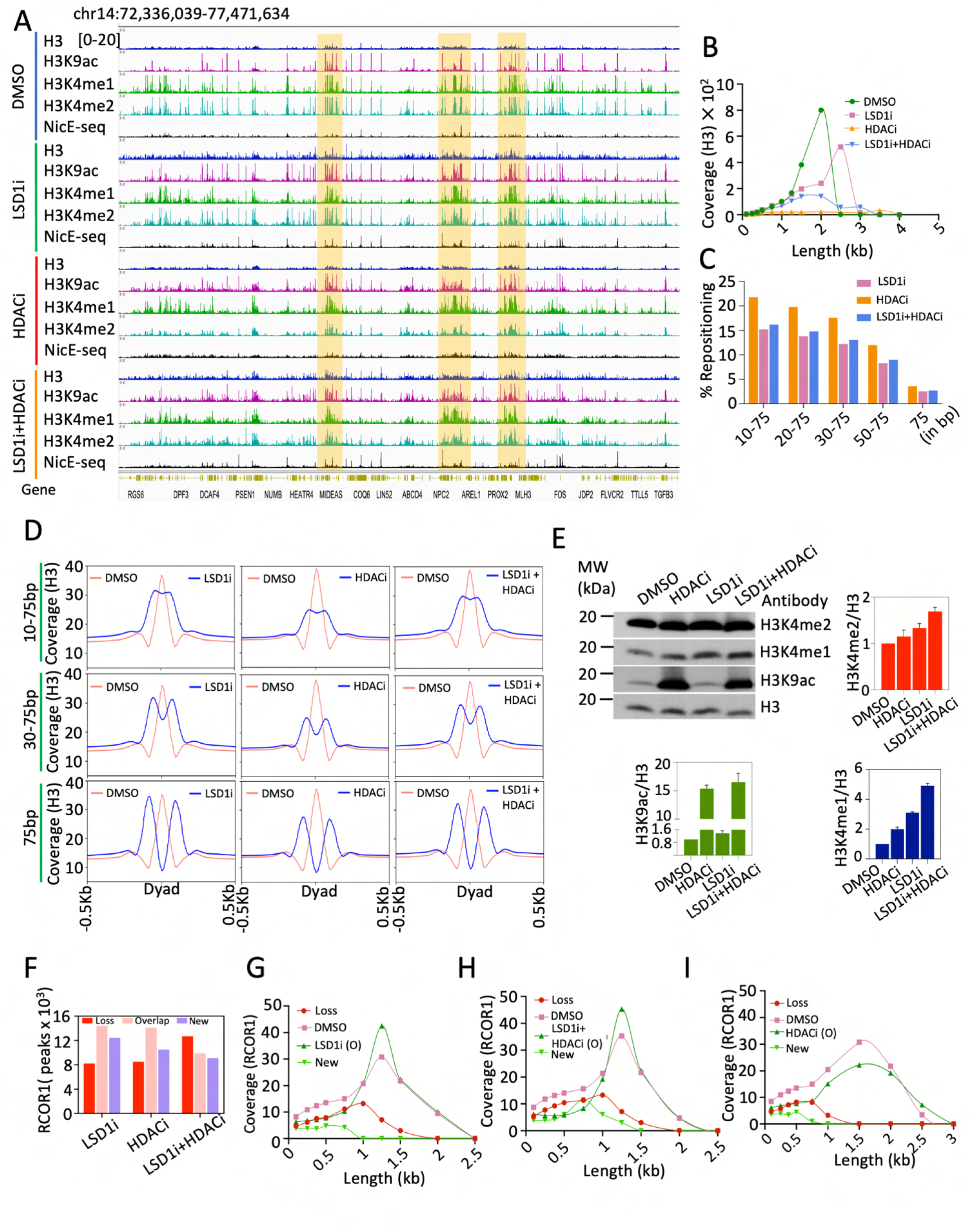
Chromatin accessibility changes associated with redistribution of histone H3 marks and RCOR1. **A.** IGV screenshots showing NEED-seq on histone H3, euchromatic marks (H3K4me1, H3K4me2, and H3K9ac) and NicE-seq generated accessible chromatin profiles for DMSO and all three treatment conditions, where changes are highlighted (yellow). **B.** Coverage vs bin plots displaying modifications in histone H3 coverage profile at genome-wide accessible chromatin regions for DMSO and all three treatment conditions, displaying histone H3 spreading on chromatin following the inhibitor treatments. **C.** Bar plots showing the percentage Histone H3 nucleosome repositioning between 10-75bp upon LSD1i, HDACi and LSD1i plus HDACi treatments respectively. **D.** Histone H3 NEED-seq enrichment showing the gradual increase in repositioning magnitude from ±10bp to 75bp for all three inhibitor application conditions. **E.** Western blots of HT1080 total cell extract and corresponding bar plots for H3K4me1 (blue), H3K4me2 (red) and H3K9ac (green) normalized to histone H3. **F.** Bar plots displaying the impact on RCOR1 binding bed regions over LSD1i, HDACi and LSD1i plus HDACi treatments respectively compared to DMSO control. **G-I.** Coverage versus bin plots displaying changes in RCOR1 coverage on peak regions extracted from Fig 3F. (O) indicates overlap.

We hypothesized that histone H3 spreading were primarily due to nucleosomal repositioning within chromatin. Using DANPOS3 (Chen et al. 2013), we analyzed nucleosome occupancy changes by comparing NEED-seq H3 profiles across different inhibitor exposure compared to the DMSO control. Nucleosome repositioning events with FDR ≤ 1% were considered for further analysis. Approximately 15–20% of total histone H3 nucleosomes exhibited positional shifts within ±10–75 bp from the dyad (Fig. 3C). A small, ∼2% of histone H3 nucleosomes (∼30,000–50,000) shifted beyond the linker region (±75 bp) with higher amplitude changes. Dyad-centered enrichment plots revealed a gradual increase in repositioning magnitude from 10 to 75 bp (Fig. 3D). Other histone marks i.e., H3K4me1, H3K4me2, and H3K9ac followed a similar pattern like histone H3 (Fig. S2D). As expected, post-HDACi treatment H3K9ac enrichment was pronounced. The combined LSD1i plus HDACi treatment produced comparable effects, reinforcing the functional crosstalk between LSD1 and HDAC.

To determine if these changes are in the nuclear genome, just not in the accessible regions, we western blotted the nuclear extract and quantified the relative amounts of H3K4me1, me2, and H3K9ac (Fig. 3E). As expected, there was significant increase in H3K9ac following HDAC or HDAC plus LSD1 inhibition, perhaps facilitating the spreading. Additionally nuclear H3K4me1 and me2 levels increased when HDAC was inhibited alone in combination with LSD1 inhibition suggesting crosstalk between both enzymes (Fig. 3E). This led us to conclude that both LSD1 and HDAC inhibitors have overlapping effect on H3K4 methylation and H3K9 acetylation.

### Chromatin accessibility alteration due to HDAC1 and LSD1 inhibition affects CoREST loading genome-wide

Since LSD1 and HDAC1/2 are part of CoREST complex, we hypothesized their inhibition can affect CoREST complex binding and transcriptional gene regulation. We compared epitope binding regions of REST corepressor, namely RCOR1, LSD1, and HDAC1, using NEED-seq, along with chromatin accessibility in both control DMSO and inhibitor treated conditions (Fig. S2E). We observed high Spearman correlation (r=0.88-0.89) between HDAC1, LSD1, and RCOR1 confirming strong correlation in control cells (Fig. S2F). In the presence of inhibitors, we observed decrease in correlation amongst them compared to control (LSD1i r=0.84-0.88; HDACi r=0.81-0.87; LSD1i plus HDACi r=0.83-0.86; Fig. S2G-I).

When either LSD1i or HDACi were treated, the peaks loss and overlap pattern for RCOR1 compared to DMSO essentially remained same (Fig. 3F). However, loss of RCOR1 peaks were more profound in LSD1i plus HDACi than mono-inhibitions. Since RCOR1 is a part of CoREST complex, next we analyzed the RCOR1 peak coverage to decipher CoREST complex binding on chromatin following inhibitor treatment. Each treatment conditions were compared to DMSO control condition and the coverage vs. bin size plots were generated based on fig. 3F. We observed bin size distribution shift of CoREST > 0.75 kb in all three inhibitor conditions (Fig. 3G-I). Secondly, we observed new peak distribution in LSD1i plus HDACi to be ∼1.5 kb demonstrating higher CoREST complex distribution due to HDAC1 plus LSD1 inhibition in the cells (Fig. 3H). These results confirm romidepsin and GSK-2879552 have a profound impact on CoREST complex.

### Runt related protein factors associate with CoREST

Chromatin accessibility controls transcription factor recruitment (Chen et al. 2024). Since we observed pronounced alteration of chromatin accessibility following HDAC plus LSD1 inhibition, we analyzed the possible transcription factor (TF) binding in the RCOR1, LSD1 and HDAC1 enriched regions in control cells. We identified 10 families of common TFs associated with RCOR1, LSD1, and HDAC1 epitope binding sites (Fig. 4A). Among the above 10 families of identified transcription factors, one was the Runt family. It was previously known that Runt family transcription factor motifs are highly enriched in LSD1 ChIP-seq (Rummukainen et al. 2022). We hypothesized that Runt family transcription factors are in complex with RCOR1, LSD1, HDAC1. To validate, we used gel filtration chromatography of nuclear cell extract and western blotted the eluted fractions with specific antibodies. Indeed, RCOR1, LSD1, and HDAC1 coeluted along with the Runt-related protein factor (RUNX), confirming their association in a complex, suggesting RUNX may be associated with CoREST (Fig. 4B).

**Figure 4.**
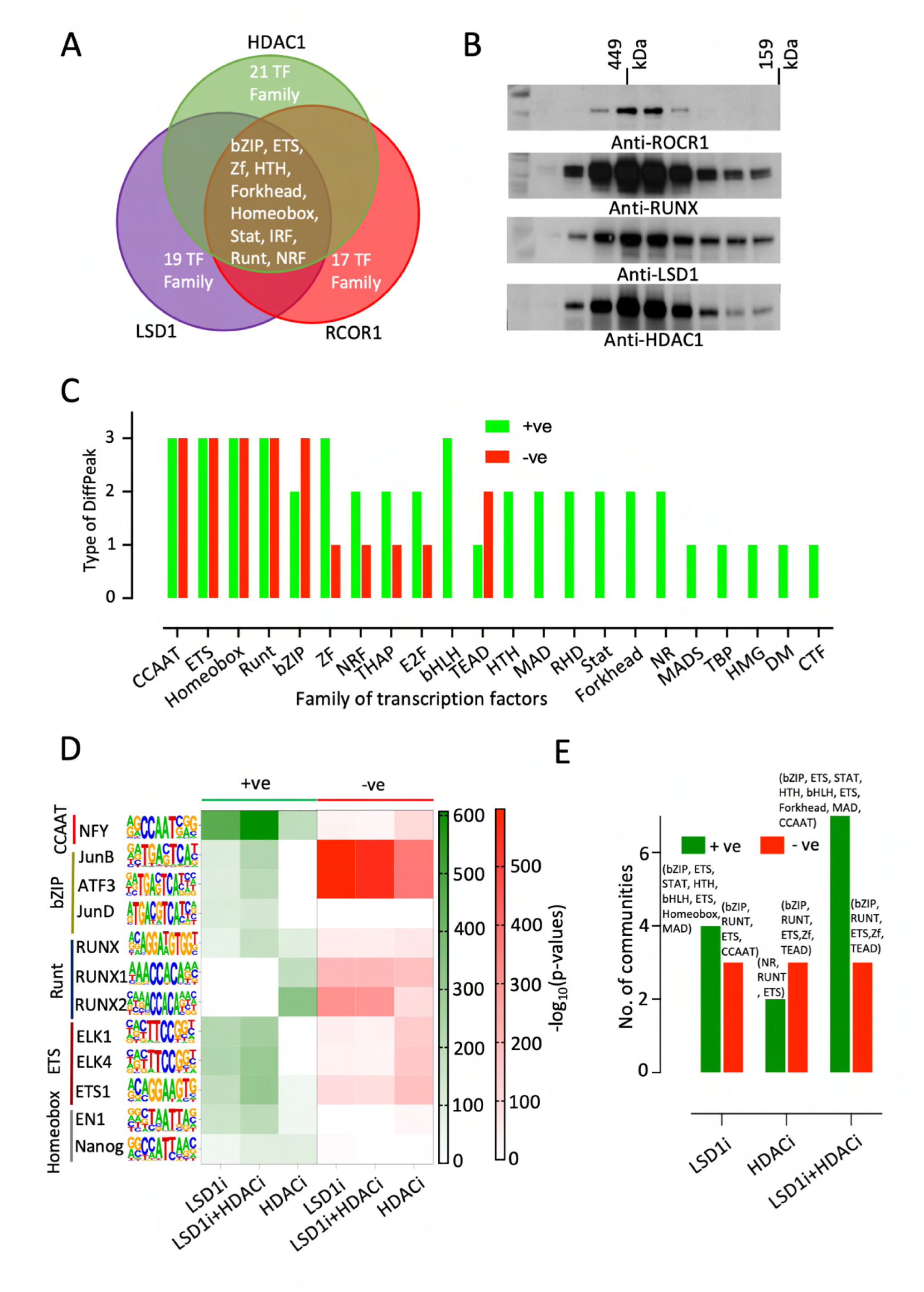
Altered chromatin accessibility modulates TFBS representation. **A.** Venn diagram showing the list of TF families commonly associated with LSD1, HDAC1, and RCOR1 NEED-seq peaks at the control condition. **B.** Western blot of fraction from gel filtration chromatography showing coelution of RUNX and the entire CoREST complex. **C.** List of TFBS families commonly identified from six different peak sets of positively (green) and negatively (red) differential NicE-seq peaks derived from LSD1i, HDACi and LSD1i plus HDACi treatments. **D.** Heatmap demonstrated-log_10_(pvalue) enrichment for top TFBS from five most frequent TFBS families i.e., CCAAT, bZIP, Runt, ETS, and Homeobox commonly identified in positive (green)/negatively (red) differential peaks. **E.** List of TFBS families commonly identified as seed nodes in TF-interaction networks given in fig S3B for positively (green) and negatively (red) differential NicE-seq peaks from LSD1i, HDACi and LSD1i plus HDACi treated cells.

### Transcription factor binding sequence identification exhibits transcription factor switch following HDAC and LSD1 inhibition

Since, transcription factor binding sequence (TFBS) could be associated with both positive and negative enriched peaks, loss or recruitment of a TF could increase or decrease the accessibility of the chromatin. Our TFBS analyses in NEED-seq peaks at the DMSO condition revealed TF list in the control condition, as shown in Fig. 4A, and these candidate TFs would be putatively perturbed over HDAC and LSD1 inhibition. To examine our hypothesis, we analyzed six differential peak sets comprising of both positive and negative enrichment accessible chromatin derived from three treatment conditions (i.e., LSD1i, HDACi, and LSD1i plus HDACi). Using Homer findMotifGenome.pl. TFBSs were predicted in six window sizes (i.e., ± 25, ± 50, ± 100, ± 150, ± 200, and ± 250 bp from the peak center). TFs that were present in at least five windows, covering almost the entire peak regions, were considered for further analysis.

When we compared the accessible chromatin from different inhibitor treatments, we predicted 22 TF families (Fig. 4C). Five TF families (CCAAT, ETS, Runt, Homeobox, bZIP) were identified in most conditions (bZIP absent in HDACi positively enriched), demonstrating correlation of NEED-seq epitope binding regions and TFBS (Fig. 4A, and C). Analysis of-log_10_(p-value) demonstrated a clear switch of enrichment in the same five families, CCAAT, bZIP, Runt, ETS and Homeobox. Specifically, JunB was enriched extensively in the negative quadrant while JunD was inclined only in the positive quadrant (Fig. 4D). Additionally, ETS and bZIP were predicted with higher frequencies (Fig. S3A) except positively enriched HDACi in the same differentially accessible regions (Fig. S3A II). This suggests there was a rapid TF dynamic following epigenetic drug treatment in cancer cells.

Next, we generated weighted evolutionarily conserved TF interaction networks for all six conditions (TFBS from six differential peaks, comprising of both positive and negative enrichment accessible chromatin derived from three treatment conditions (i.e., LSD1i, HDACi, and LSD1i plus HDACi) combining Rosetta2Fold (Zhang et al. 2025) and STRING V2.0 (Fig. S3B). Additionally, we performed Leiden clustering in the weighted networks to demonstrate numbers of interaction TF communities. Indeed, a higher number of communities in LSD1i plus HDACi positively enriched peaks was observed, suggesting maximum variability in TF interaction profiles (Fig. 4E, Fig. S3B). One reason could be genome-wide distribution of accessibility covers maximum motif binding sites upon dual inhibition, suggesting higher impact on transcriptional mechanisms. Indeed, the seed nodes from different communities were more distributed in 4 families in positively enriched peaks (i.e., bHLH, ETS, Runt, and bZIP) and 3 families in negatively enriched peaks (i.e., ETS, Runt, and bZIP; (Fig. 4E, Fig. S3C). Taken together, interactomes and TFBS motif enrichment from NEED-seq peaks as well as differential NicE-seq peaks, demonstrated ETS, Runt, and bZIP families were involved in all three inhibition conditions.

### Inversely proportionate chromatin localization of JunB with RUNX and RCOR1 following HDAC and LSD1 inhibition

Since CoREST recruitment to the JUN promoter is a critical epigenetic regulatory axis controlling cell identity and disease progression (Ismail et al. 2025), we explored the association among JunB (bZIP family), RUNX, and RCOR1 following LSD1i and HDACi treatment. We performed epitope binding chromatin analysis of RUNX and JunB using NEED-seq and compared them to RCOR1. Indeed, NEED-seq signal coverages demonstrated the colocalization of JunB, RUNX, and RCOR1 in the control condition (Fig. 5A), whereas JunB coverage was increased in both LSD1i, and HDACi conditions. This is in contrast with RCOR1 and RUNX enrichment corroborating with TFBS enrichment prediction (Fig. 5A, Fig. 4D). Next, we performed western blot of the cellular extract to validate if the chromatin enrichment of JunB was due to higher level of proteins. Indeed, both LSD1i and HDACi can boost the cellular levels of JunB, suggesting a direct correlation between JunB and its loading on the chromatin. We did observe moderate loss of RUNX protein in the cell (Fig. 5B).

**Figure 5.**
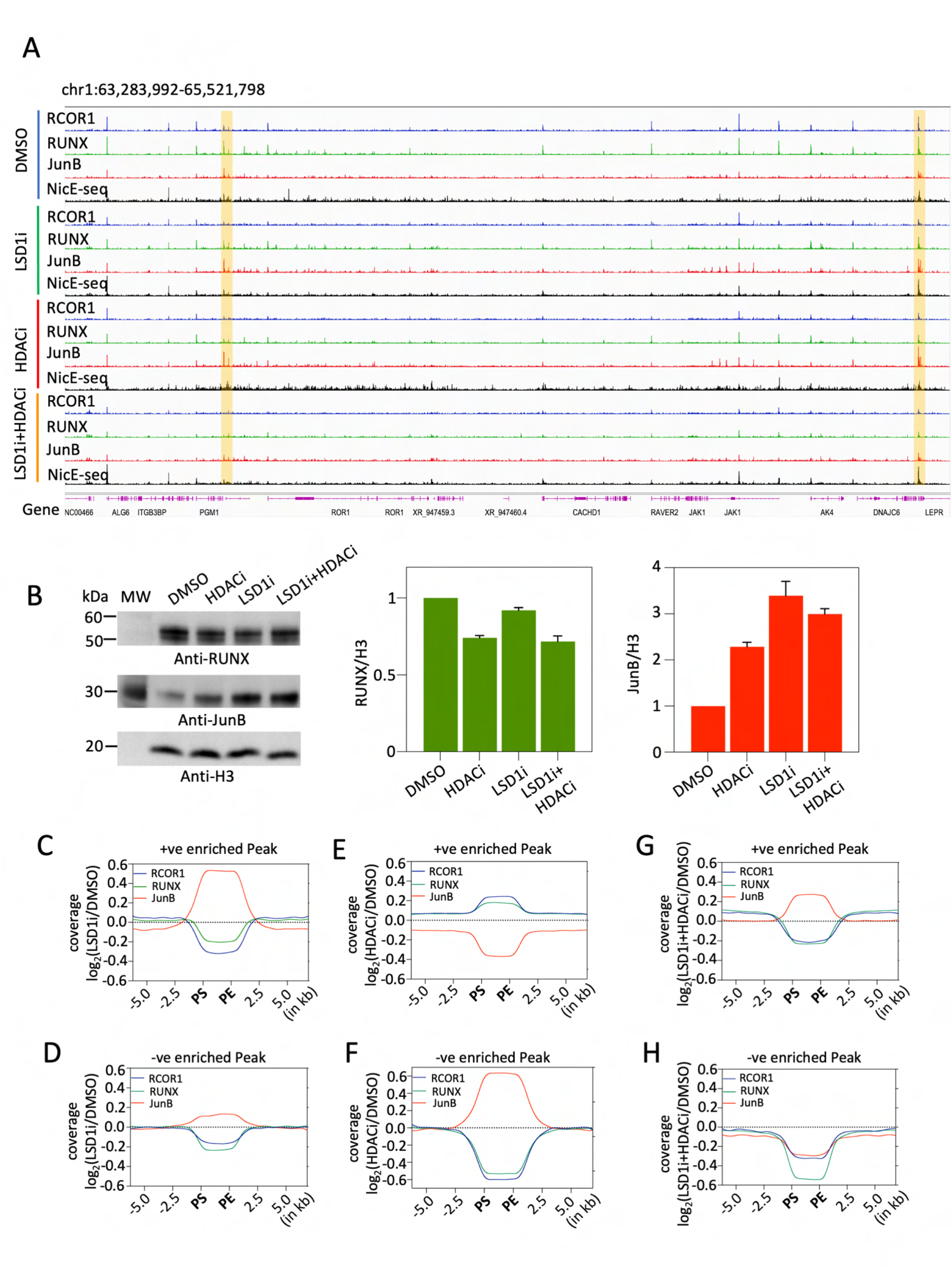
LSD1 and HDAC1 inhibition demonstrate CoREST-RUNX-JunB interplay at accessible chromatin regions. **A.** IGV screenshots showing NEED-seq on RCOR1, RUNX, and JunB in control DMSO and all three drug treatment conditions where the changes are highlighted (yellow) [Autoscaled]. **B.** Western blots of HT1080 total cell extract and corresponding bar plots for RUNX (green), and JunB (red) normalized to histone H3 (mean of two replicates). **C, E, G.** Comparative coverage enrichment is shown for CoREST (blue), RUNX (green), JunB (red) at positively enriched differential NicE-seq peaks. **D, F, H.** Comparative coverage enrichment is shown for CoREST (blue), RUNX (green), JunB (red) at negatively enriched differential NicE-seq peaks.

To determine the relative binding of RCOR1, RUNX, and JUNB on accessible chromatin regions, we perform comparative coverage enrichment analysis by mapping JunB, RCOR1, and RUNX NEED-seq in six differential NicE-seq peak regions (where positive/negative values demonstrate increase/decrease in enrichment respectively). In LSD1i differential peaks, JunB maintained inverse proportionality with RCOR1 and RUNX, where JunB exhibited higher enrichment in positively enriched peaks compared to negatively enriched peaks (Fig. 5C-D). Spearman correlation demonstrated anti-correlation between JunB and RCOR1/RUNX in negatively enriched peaks as expected (Fig. S4B-S4D). Positively enriched peaks in HDACi treatment demonstrated a reverse pattern (Fig. 5E), confirming the absence of the bZIP family in TFBS analysis (Fig S3A ii). Likewise, negatively enriched peaks in HDACi treatment exhibited a strong gain of JunB occupancy with loss of RUNX and RCOR1 on chromatin (Fig. 5F). Indeed, Spearman correlation coefficient showed anti-correlation in HDACi positive/negative differential peaks (Fig. S4C-S4D). Furthermore, dual inhibitors showed a similar trend of inverse pattern in positively enriched peaks like LSD1i (Fig. 5G). However, LSD1i plus HDACi negatively enriched peaks exhibited loss of RCOR1, RUNX, and JunB (Fig. 5H). Taken together, we demonstrated that both CoREST complex and associated RUNX protein are mutually exclusive with JunB on chromatin occupancy, suggesting differential roles in gene expression.

### Differential chromatin accessibility and nucleosome positioning regulate transcription upon LSD1 and HDAC inhibition

To further elucidate the role of chromatin accessibility, TF occupancy, and gene expression post-HDACi and-LSD1i treatment, we performed RNA-seq analysis. Differential gene expression (DEG) was performed using DMSO as a control, using p-value ≤ 0.05 and |log_2_FC| > 1. Volcano plots demonstrated the distribution of upregulated and downregulated genes in all three treatment conditions (Fig. 6A-C). As expected, the combination of LSD1i and HDACi resulted the maximum number of DEGs, mirroring higher chromatin accessibility i.e., 3532. Amongst them ∼70% were upregulated and ∼30% were downregulated (Fig. 6D). Mono-inhibitors demonstrated 540 (upregulated ∼84% and downregulated ∼16%) and 2155 (upregulated ∼58% and downregulated ∼42%) DEGs in LSD1i and HDACi treatments, respectively (Fig. 6D).

**Figure 6.**
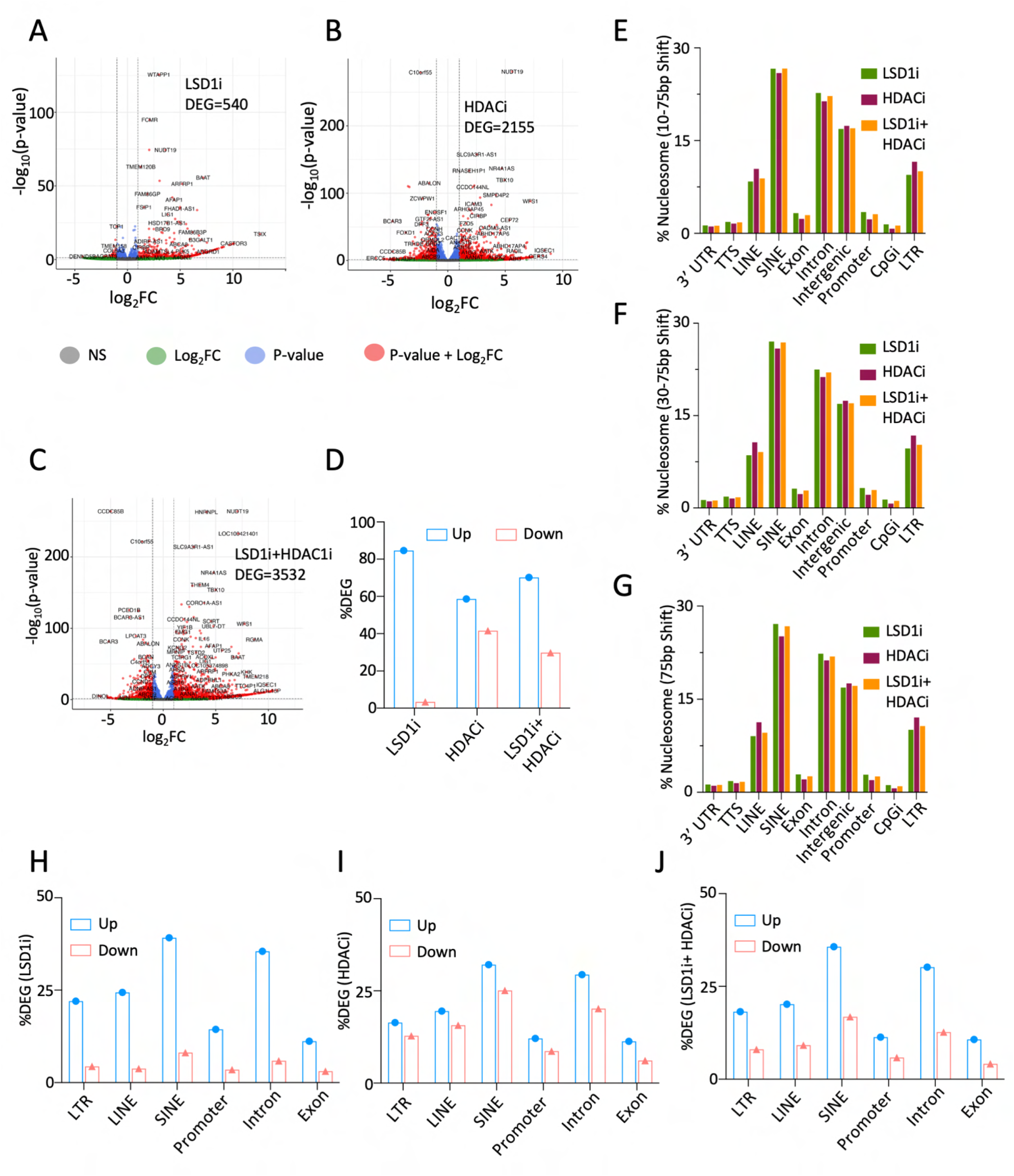
Histone H3 repositioning modulates transcription. Volcano plots showing the distribution of differentially expressed genes (DEG) following LSD1i (A), HDAC1i (B) and LSD1i plus HDACi (C). **D.** Percentage of up (blue) / down (red) regulated genes from LSD1i, HDACi and LSD1i plus HDAC1i using bar plots are shown. Genomic annotations displaying shifted nucleosomes genome-wide localization for 10-75bp (**E**), 30-75bp (**F**), and 75bp (**G**) repositioning. Percentage of up (blue) / down (red) regulated genes commonly identified as nearest RNA features associated with annotated histone H3 regions from LTRs, LINE, SINE along with intron, exon and promoters following inhibitor exposure, LSD1i (**H**), HDACi (**I**) and LSD1i plus HDACi (**J**).

Genomic annotation of repositioned nucleosomes from histone H3 and associated histone marks from NEED-seq unveiled prominent localization at the LTRs (Long Terminal Repeats), LINEs (Long Interspersed Nuclear Elements), SINEs (Short Interspersed Nuclear Elements), as well as within introns, exons, and briefly in promoter regions, supporting plausible modulation of expression in both transposable elements (TEs) and mRNAs (Fig. 6E-G).

To investigate chromatin dynamics with transcriptional changes, DEGs were mapped to the nearest RNA features associated with annotated genome-wide shifted histone H3 regions. LSD1i and LSD1i plus HDAC1i treated cells demonstrated a clear bias towards gene up-regulation, whereas HDAC1i treatment produced a more equivalent distribution of up-and down-regulated DEGs (Fig. 6H-J, like Fig. 6D). Intriguingly, nucleosome repositioning within intronic regions appeared to play a substantial role in transcriptional regulation. When we mapped the promoter-specific genes from differential accessible chromatin regions from fig 2E, we observed synergy with both up and downregulated genes. As shown, LSD1i and LSD1i plus HDACi positively enriched peaks associated with upregulated genes. This is complementary with HDACi negatively enriched peaks that was mostly associated with down regulated genes. (Fig. S5A-C). Therefore, differential accessibility at promoters, and the nucleosome repositioing correlated with transcriptional regulation.

To define the contributary role of TFs on accessible chromatin and gene expression, we extrapolated TFs associated with DEGs. We observed about half of TFs identified from DEGs overlapped with TFBSs for all three treatments. Specifically, JunD and JunB were associated with downregulated genes (Fig. S5D). Next, we analyzed the significant Gene Ontology Biological Process (GO:BP) for overlapping genes using data shown in fig S5A-C (Fig. S5E-J). While most of the pathways exhibited systematic cell death, few specific pathways emerged. Cell cycle arrest at HDACi downregulated genes, telomeric shortening pathways with LSD1i up regulated genes, inhibition of ribosome biogenesis in LSD1i plus HDACi down-regulation genes were prominent. The detailed list of GO:BP is listed in Table S2. Taken together, the transcriptional regulation due to perturbed accessibility profiles and transcription factor recruitment post epigenetic inhibitor treatment resulted in activation of genes associated with apoptotic pathway. Therefore, we suggest that dual epigenetic inhibitors have more efficacy on cancer epigenome.

## Discussion

Our newly developed NicEL can proficiently detect modifications in chromatin accessibility reflecting the impact of epigenetic drug treatments. NicEL also exhibits the accessibility modulation “per cell/nuclei” basis, showing the gradual progression of treatments within the cell population. Finally, higher compatibility between sequencing and ACI suggested that NicEL could be an efficient model for early detection of drug efficacy in terms of accessibility modulation. The limited efficacy of single pharmacological agent or mono-inhibitory therapies in malignancies has been a challenge, primarily due to drug resistance and compensatory mechanisms (Al-Lazikani et al. 2012, Brown et al. 2014). Therefore, synergistic therapeutic protocols or combination therapy employing two or more drugs are routinely used for enhancing treatment efficacy. Designing a multi-faceted therapeutic approach using machine learning, such as NicEL, could be one such solution. However, allosteric variations triggered by multi-target approaches may result in increased toxicity. Hence, systematic inhibition through targeting multiple enzymes within the same enzymatic complex could offer a more feasible strategy.

In this context, targeting the CoREST complex typically involves modulating two major members i.e., LSD1 and HDAC1/2. While no potent LSD1 inhibitors have yet been approved by the FDA, romidepsin is a well-established FDA-approved HDAC inhibitor used for cutaneous and peripheral T-cell lymphomas (Bates et al. 2015). Corin, a dual-inhibitory FDA-approved agent, has demonstrated histone demethylase activity comparable to that of GSK-2879552 and tranylcypromine (catalytic LSD1 inhibitors) (Bauer et al. 2019, Ulrich et al. 2017). Likewise, the synergistic efficacy of corin has already been reported (Kalin et al. 2018). In corin, the synthetic deacetylase inhibitor MS-275 is known for its target specificity compared to romidepsin which is more generically targeting Class I and II HDAC. Based on enzymatic function, therapeutic targeting of the CoREST complex, and its individual components is expected to enhance chromatin accessibility. Our results using NicE-viewSeq revealed comparable levels of accessibility enrichment, highlighting dynamic chromatin accessibility changes upon treatment. Notably, ∼86% of LSD1 associated with chromatin is known to be complexed with CoREST, which may explain the similarity between the accessibility patterns observed following LSD1i and LSD1i plus HDACi. Also, systematic pre-treatment with an LSD1i appeared to restrict the accessible chromatin randomization caused by HDACi. Cheng et al. further demonstrated an alternative role of LSD1 in regulating H3K27ac, showing that LSD1 knockout leads to H3K27ac enrichment at promoters, suggesting larger than expected impact on CoREST complex.

Beyond the equivalent impact observed on all three histone marks (i.e., H3K4me1, H3K4me2, and H3K9ac), our treatments modalities involving LSD1i, HDACi, and their combination, unveiled a convergent mechanism involving bZIP (JunD, JunB, etc.), RCOR1, and Runt (RUNX) family transcription factors. In breast cancer, the potential for complex formation between c-Jun and CoREST has been demonstrated, particularly in association with EMT-related genes (Garcia-Martinez et al. 2022). Our findings suggested a possible replacement of the CoREST complex by JunB and JunD, which may enhance heterodimeric AP-1 formation and antagonize c-Jun (Dong et al. 2023, Lee et al. 2012, Deng et al. 1993), specially in fibrosarcoma contexts.

The interplay between RUNX and CoREST can occur with two CoREST members, LSD1 and HDAC1 individually (Yu et al. 2021, Ali et al. 2012). Our data also demonstrated a proportional loss of both RUNX and CoREST complex components following HDAC and LSD1 inhibition. This is partially supported by switching between RUNX and Jun motifs in normal and malignant cells and their association with chromatin accessibility in romidepsin-mediated HDAC inhibition (Qu et al. 2017). Taken together, these observations suggest RUNX and Jun dynamics may be a common occurrence in accessible chromatin spreading upon histone deacetylase and histone demethylase inhibition that needs further investigation. Consistent with these findings, we observed marked differences in transcription factor interactomes upon treatment. Nonetheless, bZIP and Runt family proteins consistently emerged as seed nodes, supporting a shared mechanistic outcome between mono-and combined-inhibitory conditions. Additionally, transcriptomic and differential NicE-seq analyses further revealed regulation of apoptotic pathways, cell-cycle progression, chromatin organization, etc. following the inhibitor treatment indicating systematic induction of cell death across all three treatments. This was supported by NF-Y associated with histone deacetylation, where NF-Y mediated apoptosis is known to be triggered by HDAC inhibition (Roy et al. 2005). Indeed, regulome analyses on NF-Y (CCAAT) have revealed higher enrichment and motif overlap with families such as bZIP, Zf, bHLH, and CTF, which vary across cell types, suggesting their crosstalk (Dolfini et al. 2016, Ronzio et al. 2019). Therefore, integration of these multiomic datasets could provide a coherent and functionally precise interpretation of the chromatin response and gene expression to therapy.

The impact of epigenetic targeting on TE expression has previously been reported for DNMT1 inhibition (Ohtani et al., 2020), HDAC inhibition (Goyal et al., 2023), and LSD1 inhibition (Sheng et al., 2018), particularly at LTRs. Our annotation analysis suggests histone repositioning as a potential mechanistic driver of such modulation. Our observations indicate that modulation of chromatin accessibility through epigenetic targeting induces histone repositioning, which in turn is associated with transcriptional regulation of both genes and transposable elements. In summary, multi-modal NicE-viewSeq platform, which combines spatially resolved chromatin accessibility profiling at the single-nucleus level while preserving the native chromatin context (NicEL), with high-throughput sequencing to identify specific genomic loci, enables a detailed elucidation of chromatin landscapes including nucleosome repositioning. Together, this multi-modality constitutes a robust framework for assessing drug-induced chromatin accessibility alterations and deciphering transcriptional reprogramming associated with distinct therapeutic regimens.

## Materials and Methods

### Chemicals and epigenetic drug treatments

HT1080 cells were grown according to ATCC’s recommendations on slides (VWR micro cover glass # 48366067) in 6 well plates. For combination drug treatment, HT1080 grown with or without 5 μM LSD1 inhibitor (GSK # 2879552) for 3 days. DMSO was used as control. After 3 days, control and treated cells LSD1 inhibitor were harvested and were grown on slides with or without 1 μM of HDAC inhibitor (Romidepsin, Sigma-Aldrich # SML1175) for 6h at 37°C. For single epigenetic drug treatment, HT1080 were grown on slides with or without epigenetic inhibitors (10 μM) for 12h. DMSO used as control. Epigenetic drugs used were: WDR5-0103 (Tocris # 5323), PFI-2 (Xcessbio # M8499-2), UNC0379 (Selleckchem # S7570) and JQ-1 (Selleckchem # S7110).

### NicE-view

After crosslinking cells with 4% formaldehyde and 1.5 M Tris.Cl pH 8 quenching, 0.1% SDS (in 1x NEBuffer^TM^2) was added for 10 min at 58°C to remove cytoplasm. SDS was subsequently quenched with 1% Triton X-100. Fluorescent accessible chromatin DNA labeling was performed by incubating the nuclei in the presence of 2 U of Nt.CviPII (NEB R0626S), 10 U of DNA polymerase I (M0209S) and 30 μM of each dNTP, including 6 μM of Fluorescein-12-dATP (Perkin Elmer, NEL465001EA) in 800 μl of 1 × NEBuffer 2 per reaction. Reaction was carried out at 37 °C for 1h and subsequently stopped by adding 5 mM EDTA. 3 μl of RNAse A (NEB # T3018L) was also added to the labeling reaction and incubated at 37 °C for 30 min to digest cellular RNA. Slides were washed once with NEED-seq washing buffer (20 mM phosphate buffer, 0.5 M NaCl, and 1 % Triton X-100) and were air dried, mounted using Prolong Gold antifade reagent with DAPI (Invitrogen, P36935) and visualized using LSM 880 confocal microscope (Zeiss). Fluorescein-dATP, Texas Red–dATP and DAPI were detected using Argon 458, 488, 514 nm, DPSS 561 nm, and diode 405 nm laser respectively. Quantification of Fluorescein-dATP or Texas Red–dATP per nucleus was defined by mean pixel intensity per nucleus using the histogram and colocalization tools included in the Zen software (Zeiss).

### NicE-seq

After drug treatment (see above), cells grown in 6-well plates were crosslinked with 4 % formaldehyde. Formaldehyde was quenched with 1.5 M Tris. Subsequently, PBS + 0.1% SDS was added for 10 min at 58°C and quenched with PBS + 1% Triton X-100 for 5 min. After washing with PBS, the samples were processed as described in Esteve et al. 2020. For library amplification, 12 PCR cycles were implemented using the program for Ultra II NEBNext Ultra II. PCR products were purified using 0.9x volume of NEBNext Sample Purification beads. 1 nM of size selected DNA library was sequenced on NovaSeq 6000.

### NEED-seq

NEED-seq size selection protocol was used as described previously (Sen et al., 2025). Briefly, HT1080 cells treated with combination of drugs as described above were grown on 6-well plates to 80% confluency. Cells were then incubated for 5 min at 4°C using CSK buffer to remove cytoplasm, followed by crosslinking with 4% formaldehyde for 10 min at RT. 1.5 M Tris.Cl, pH 7.5 was used to quench formaldehyde for 5 min at RT. After 1x PBS wash, blocking buffer was added to cells for 1h at RT. Different antibodies (see Table S3) were then added at dilutions recommended by the manufacturer and incubated with cells for overnight at 4°C. After 3 washes with 1x T-PBS for 5 min at RT, blocking buffer was added with fluorophore conjugated antibody for 1h at RT. The samples were processed as described previously (Sen et al., 2025). The library was amplified using 12 PCR cycles. PCR products were purified using 0.9x volume of NEBNext Sample Purification beads. 1 nM of size-selected DNA library was sequenced on NovaSeq 6000.

### RNA-Seq

1 μg of RNA (purified from NEB # T2110S) was used (in triplicates for all drug treatments). RNA-Seq was performed using NEB # E7760S kit.

### Western blot

To extract the total amount of protein from HT1080 cell (treated with combination of drugs, see above), cell pellets were resuspended in 50 mM Tris.Cl pH 7.5, 200 mM NaCl and 1% SDS. SDS was quenched with 10% Triton X-100. After sonication (10 x 10 sec pulses), cell extracts were loaded on SDS-PAGE. Western blots were performed using H3K9ac, H3K4me1/2, H3, LSD1, HDAC1 and Jun B and RUNX antibodies (see Table S3).

### NicEL

NicEL is an automated learning model for assessing NicE-view microscopic images where nuclear instance segmentation was performed using a deep learning based StarDist framework with a U-Net backbone (Vijayan et al. 2024). A pre-trained StarDist model was fine-tuned on manually curated instance segmentation masks, in which each nucleus was assigned a unique identifier, using a low learning rate and extensive geometric and intensity-based data augmentation. The model jointly predicted per-pixel object probabilities and star-convex radial distance maps, enabling accurate delineation of densely packed nuclei. Final nuclear instances were obtained via non-maximum suppression applied to candidate star-convex polygons. The resulting masks were applied to multi-channel fluorescence images to extract nucleus-specific intensity values. For each nucleus i, the Accessible Chromatin Index (ACI) was defined as

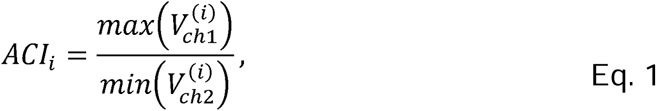

where *v_ch1_*^(i)^ and *v_ch2_*^(i)^ denote the non-zero voxel intensity sets of the chromatin accessibility marker and the nuclear reference channel, respectively. Nuclei with empty intensity sets or zero-valued maxima in the target channel were excluded from downstream analysis. More details are given in Supplementary Material 1.

### NEED-seq and NicE-seq data analysis

NEED-seq and NicE-seq analysis was started from Adaptor trimming. Applying Trim Galore, low-quality sequences and adaptor sequences were trimmed (http://www.bioinformatics.babraham.ac.uk/projects/trim_galore) from paired-end sequencing reads using the commend line interface (CLI): --clip_R1 4 –clip_R2 4 – three_prime_clip_R1 4 –three_prime_clip_R2 4. Subsequently, Bowtie2 (Langmead et al., 2012) was applied on trimmed read pairs to map reference genome (hg38) with the following setup: --dovetail –no-unal –no-mixed –no-discordant –very-sensitive-I 0-X 1000. After getting sam alignment files from Bowtie2, the files were converted to bam files using “samtools view-S-b” (Li et al., 2015) which were further purified by removing PCR duplicates using following settings: “picard MarkDuplicates REMOVE_DUPLICATES=true” and mitochondrial reads (http://broadinstitute.github.io/picard/).Resultant aligned read pairs were utilized for peak calling with MACS3 (Zhang et al., 2008). From NEED-seq, we generated two different set of peaks, (1) Macs3 call peak-f BAMPE-m 4 100 –bdg –SPMR (for narrow peak); (2) Macs3 call peak-f BAMPE-m 4 100 –bdg --broad –SPMR (for broad peak). For NicE-seq, we have called only narrowPeaks. We have generated five NicE-seq replicates each for DMSO and three treatment conditions.

### Differential peak analysis

To identify differential accessible chromatin peaks, DiffBind (Stark et al. 2011), a R package, was utilized. We have generated three replicates each for controlled and treated condition. Generated peak for each replicated along with the bam files were provided as input. DEseq2 option was selected in dba.analyze command where peaks with p-value ≤ 0.05 were considered as differential peak. Next, volcano plot was generated using dba.plotVolcano where log_2_FC < 0 were considered as-ve enriched peaks and vice versa.

### Peak annotation and Venn diagram

We have used Homer “annotatePeak.pl” to determine the genomic localization of the differential NicE-seq peak for hg38 assembly. Next, “intervene venn” (Khan et al. 2017) was utilized to check the sharing regions for positively and negatively enriched peaks between the treatments.

### Coverage distribution analysis

To study the NEED-seq distribution within NicE-seq peaks upon treatments, we have applied “bedtools coverage” where NEED-seq bam files used to calculate the coverage with NicE-seq bed regions. Further, the bed regions were divided into different bed size bins, and mean coverage was calculated for each bin and plotted using Prism 10.

### Enrichment analysis

Enrichment analyses were performed to study the enrichment of NEED-seq histone marks and TF NEED-seq. In this case, we have used two types of bigwigs i.e.,1. bigwig files generated using “bamCoverage” normalized to RPKM for histone marks and, 2. comparative bigwig files were generated using “bigwigCompare”–-operation “log2” where TF enrichment compared to control conditions. The first bigwig sets can only displayed read depth whereas second set demonstrated comparative gain/loss of enrichment upon the treatments. Now, histone mark enrichment in positively enriched NicE-seq peaks was performed using computeMatrix and plotProfile where we have generated two sets of peaks using hierarchical clustering. Next, these sets of peaks separately utilized to calculate modifications in histone mark enrichments. Further, histone mark enrichment in negatively enriched NicE-seq peaks was performed using computeMatrix and plotHeatmap. For TF NEED-seq, the enrichment analysis was performed for all the differential peaks and RCOR1 peak sets. Bigwig signals were shown using Integrative Genome Browser (IGV) (Thorvaldsdóttir et al. 2013).

### Nucleosome repositioning analysis

To calculate nucleosome repositioning, we used DANPOS3 (Chen et al. 2013). We have used Histone H3, H3K4me1, H3K4me2, and H3K9ac NEED-seq profile to perform comparative repositioning where DMSO was used as control for “–-paired 1 –-frsz 147” parameters where significant repositioning were filtered based on FDR ≤ 0.01.

### Correlation analysis

We have in general, performed Spearman correlation analysis where RCOR1, LSD1 and HDAC1 NEED-seq were compared using “multiBamSummary” and comparative bigwigs, generated from RCOR1, RUNX and JunB, were compared within differential peak regions.

### TFBS identification and interactions network analysis

For TFBS identification, we have used Homer “findMotifGenome.pl” for hg38 assembly where –-size parameters used six window sizes (i.e., ± 25, ± 50, ± 100, ± 150, ± 200 and ± 250 from the peak center). Next, 1e-10 was considered as p-value cutoff for TFBS selection.

Next selected TFBS from differential peaks were analyzed for protein-protein interaction using STRING V2.0 database (Szklarczyk et al. 2023). Next, Rosetta2Fold (Zhang et al. 2025) was used to calculate evolutionarily interaction probability where the interaction sets were filtered by considering a probability score ≥ 0.30. The remaining TF interactions were used to create a weighted network G where (V, E) ∈ G, V defines the TFs as nodes where E defines weighted edges where the interaction probabilities were used as weight (w). These weighted networks were further divided in communities based on Leiden clustering, where minimum members of the community should be more than 5. We also determined top three seed nodes jointly based node betweenness and weighted degree centrality.

### RNA-Seq data processing

For RNA-Seq, four different data sources i.e., DMSO, LSD1i, HDACi, and LSD1i plus HDACi were used two replicates each. We have started by aligning raw RNA-seq data. After quality check and adaptor trimming, STAR aligner was used to map the refined fastq with hg38 reference genome (Dobin et al. 2013). The mapped reads were used to identify the transcripts as count matrix applying htseq (Anders et al. 2015). Next, the count matrices were considered to identify differentially expressed genes (DEGs) applying DESEQ2 (an R package) (Love et al. 2014) comparing to DMSO control. DEGs were identified based on p-value ≤ 0.05. Following that, UP and down regulated transcript were selected based log_2_FC (Fold Change) ≥ 1 and log_2_FC ≤-1 respectively.

---

**Supplementary Figure 1. Parallel analysis displaying higher convergence between both modal outputs. A.** Enrichment in chromatin accessibility upon LSD1i, HDACi and LSD1i plus HDACi treatments at genome-wide accessible peak ± 2kb is shown using heatmaps. **B.** Percentage distribution of positive (Green) and negatively (Red) enriched peaks generated using DiffBind. **C-E.** MA plots of NicE-seq peaks showing overlapping ACI (accessible chromatin) and peak profile. **F-H**. Density plots showing the distribution of absolute log_2_FC values of differential peaks (Blue) and ACI scores (Red) for LSD1i, LSD1i plus HDACi and HDACi respectively demonstrating higher convergence. **I-K.** Violin plots showing distribution of absolute log_2_FC values of differential peaks (blue) and ACI scores (red) for LSD1i, LSD1i plus HDACi and HDACi respectively, where p-value generated by Kolmogorov–Smirnov test (K-S test) between differential peaks and ACI scores accepted H_0_ hypothesis.

**Supplementary Figure 2. LSD1 and HDAC1 inhibition impacting accessible chromatin (NEED-seq) profile of euchromatic marks and its impact on CoREST complex. A-C.** Coverage vs bin plots displayed modifications in NEED-seq coverage profiles of H3K9ac, H3K4me1 and H3K4me2 at genome-wide accessible chromatin regions for DMSO and all three treatment conditions. **D.** H3K4me1, H3K4me2, and H3K9ac NEED-seq enrichments demonstrating ±75bp repositioning upon all three inhibitor treatment conditions. **E.** IGV screenshots showing NEED-seq of HDAC1, LSD1, RCOR1, and NicE-seq profiles for DMSO and all three treatment conditions with change regions being highlighted (yellow) [Autoscaled]. **F-I.** Heatmaps based on Spearman correlation coefficient using NEED-seq reads to determine correlation between HDAC1, LSD1 and RCOR1 in DMSO, LSD1i, HDACi, and LSD1i plus HDACi treated cells.

**Supplementary Figure 3. The list of TFBS families identifies Runt, bZIP, and ETS as most represented families. A.** Pie chart of the number of TF members from each selected familiy identified using Homer [i-vi]. **B.** TF-interaction networks and predicted communities using Leiden clustering is shown. **C.** Word cloud showing presence of TF families identified as seed nodes in each community from networks of fig S3B where the size of the word exhibited the frequency of its presence (Venn diagram, middle panel).

**Supplementary Figure 4. LSD1 and HDAC1 inhibition impact on NEED-seq RCOR1 enrichment.** Heatmaps based on Spearman correlation coefficient using NEED-seq comparative coverage to determine correlation between RUNX, JunB, and RCOR1 in positive/negatively enriched differential NicE-seq peaks **A,B**. LSD1i, **C,D**. HDACi and **E, F**. LSD1i plus HDACi treated cells.

**Supplementary Figure 5. Altered chromatin accessibility regulates transcription. A-C.** Percentage of commonly identified up (blue) / down (red) regulated genes in promoter Specific differential peak locations (as shown in fig 2E upon LSD1i, HDAC1i, and LSD1i plus HDAC1i) is shown using bar plots. **D.** Overlap between TFs targeted DEGs and TFBS identified from promoter specific differential peaks where Not-I (representing the set not found in differential peaks, grey); Ov (common TFs targeting both Up/Down regulated genes (light magenta), and Up(U) (TFs uniquely targeting upregulated genes (blue), and Down(U) (TFs uniquely targeting down regulated genes, red) is shown. GO: BP terms identified are shown. **E-F** for LSD1i, **G-H** for HDACi and **I-J** for LSD1i plus HDAC1i are presented with down (red) and up (blue) genes observed in fig 6E-G are shown using bubble plots.

## Table Legends

**Supplementary Table 1:** List of therapeutic agents and their targets.

**Supplementary Table 2:** Total list of GO:BP terms identified from LSD1i; HDACi and LSD1i plus HDAC1i studies. **A.** up and **B.** down regulated genes represented in Fig 6E-G.

**Supplementary Table 3:** List of antibodies used for **A.** NEED-seq, and **B.** Western Blots

**Supplementary Material 1:** Complete description of NicEL Stages and Model Architecture.

## Data availability

NEED-seq, NicE-seq data performed in this study are available in NCBI Gene Expression Omnibus (GEO)

## Code availability

NicEL pipeline is available in https://github.com/Sagnik-Epigen/NicEL

## Declaration of generative AI and AI-assisted technologies in the writing process

During the preparation of this work the author(s) didn’t use any Generative AI assisted technology.

## Acknowledgments

We thank T. Evans, Sir R.J. Roberts, and S. Russello for encouragement. The project was funded by basic research grant to SP from New England Biolabs, Inc., and partly funded by R44HG011875 from NIH, Fulbright-Nehru Academic and Professional Excellence Fellowship Award No. 3087/F-NAPE/2024 from USIEF to UM, and JC Bose Fellowship grant No. JBR/2021/000036 from SERB, Govt of India to SB.

## Author contribution

POE, SS, and RD performed experiments. NicEL development and bioinformatic analysis were performed by DT, and SS. AD performed the beta testing of NicEL. SS, and SP conceptualized and planned experiments and wrote the manuscript with input from POE, RD, AD, and DT. The work was supervised by SP, UM, and SB.

