## Supplementary for "Altered chromatin accessibility and nucleosome positioning landscape upon HDAC and LSD1 inhibition in cancer cell"

Suppl Figure 1

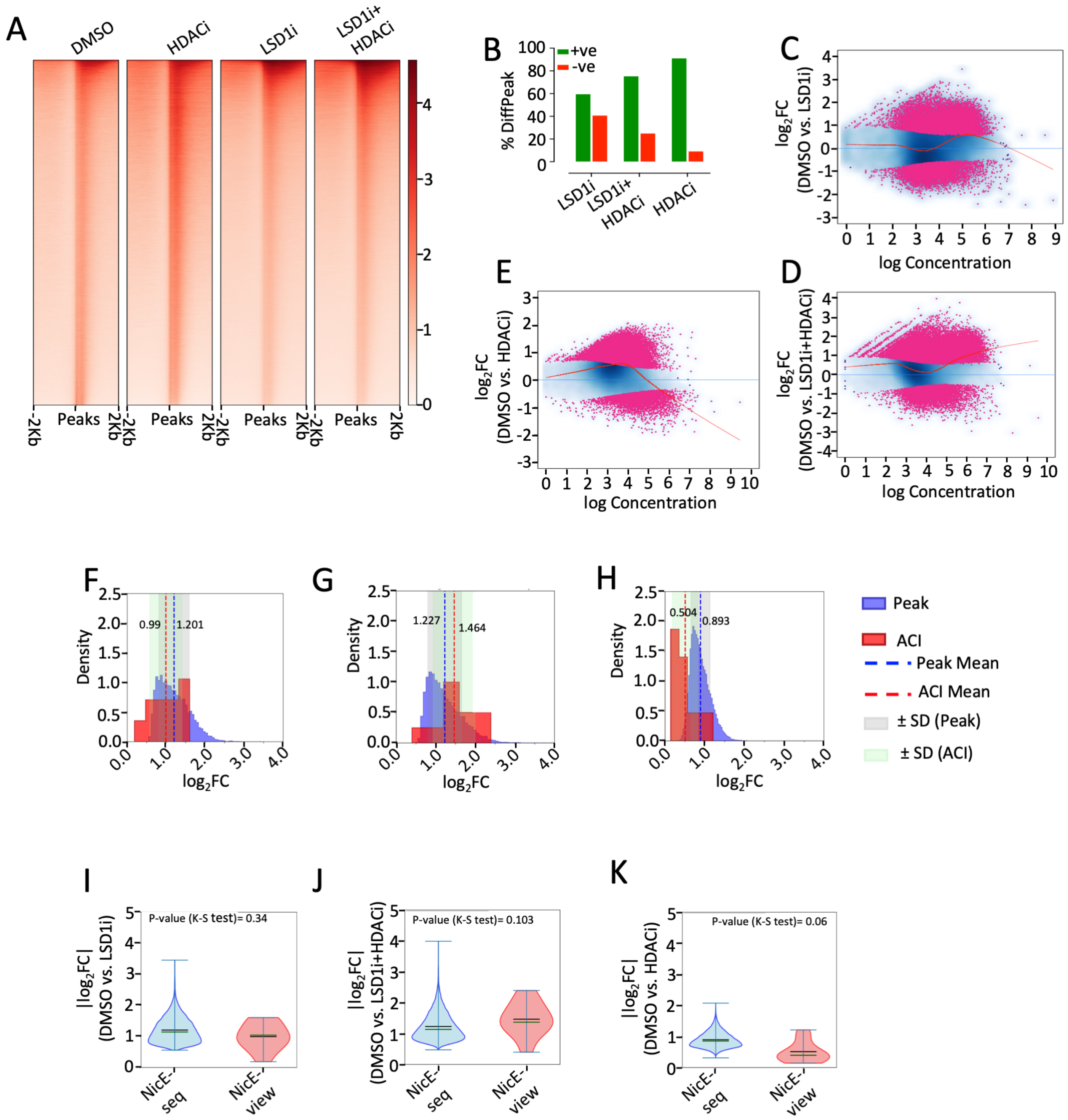

Suppl Fig. 2

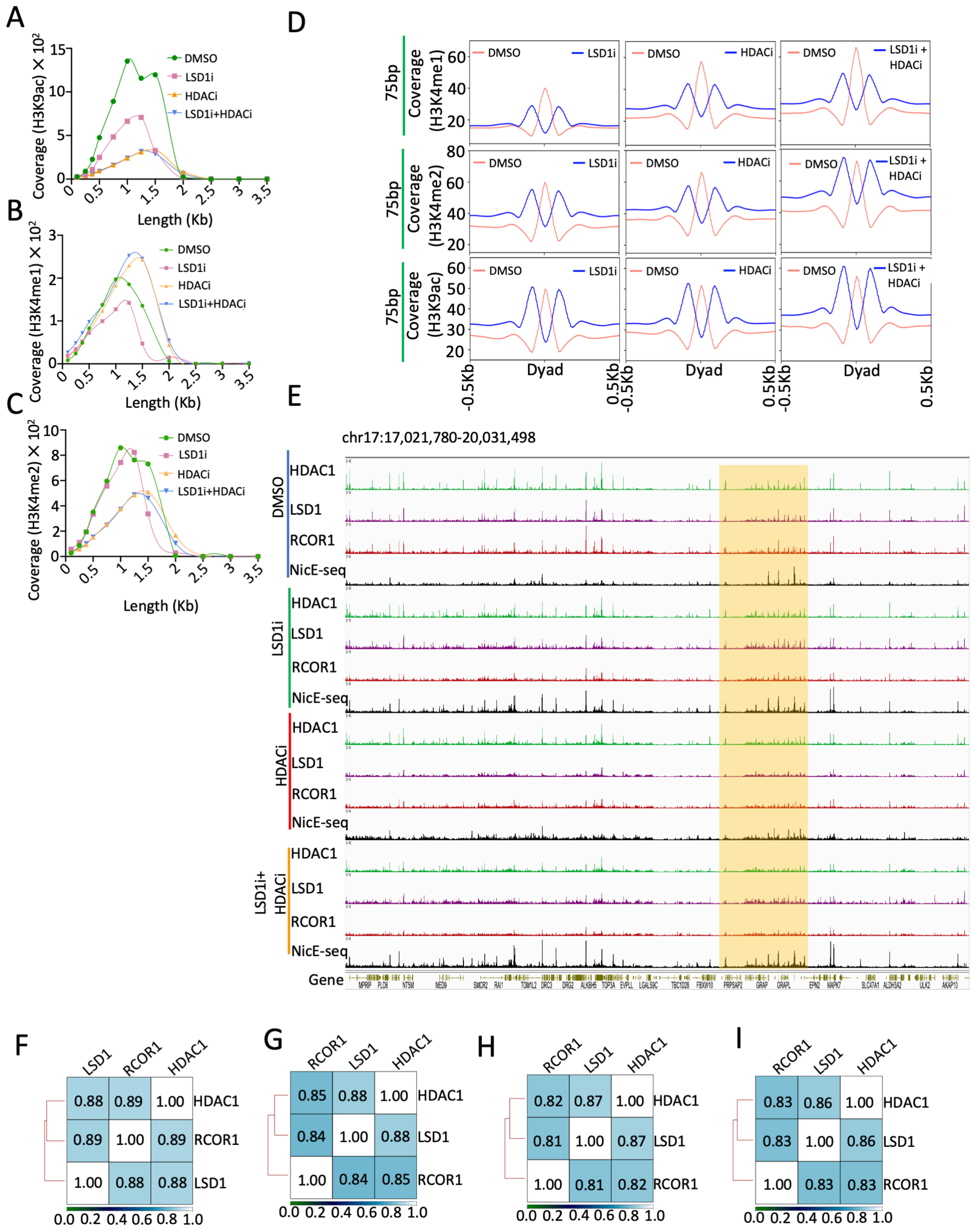
A
B
C
D
E
F
G
H
I
Coverage (H3K9ac) × 10²
Coverage (H3K4me1) × 10²
Coverage (H3K4me2) × 10²
Length (Kb)
DMSO
LSD1i
HDACi
LSD1i+HDACi
75bp
Coverage (H3K4me1)
Coverage (H3K4me2)
Coverage (H3K9ac)
LSD1i + HDACi
Dyad
-0.5Kb
0.5Kb
chr17:17,021,780-20,031,498
HDAC1
LSD1
RCOR1
NicE-seq
Gene
MPRIP
PLD6
NT5M
MED9
SMCR2
RAI1
TOM1L2
DRC3
DRG2
ALKBH5
TOP3A
EVPLL
LGALS9C
TBC1D28
FBXW10
PRPSAP2
GRAP
GRAPL
EPN2
MAPK7
SLC47A1
ALDH3A2
ULK2
AKAP10
0.88 0.89 1.00 HDAC1
0.89 1.00 0.89 RCOR1
1.00 0.88 0.88 LSD1
0.85 0.88 1.00 HDAC1
0.84 1.00 0.88 LSD1
1.00 0.84 0.85 RCOR1
0.82 0.87 1.00 HDAC1
0.81 1.00 0.87 LSD1
1.00 0.81 0.82 RCOR1
0.83 0.86 1.00 HDAC1
0.83 1.00 0.86 LSD1
1.00 0.83 0.83 RCOR1
0.0 0.2 0.4 0.6 0.8 1.0

Suppl Figure 3

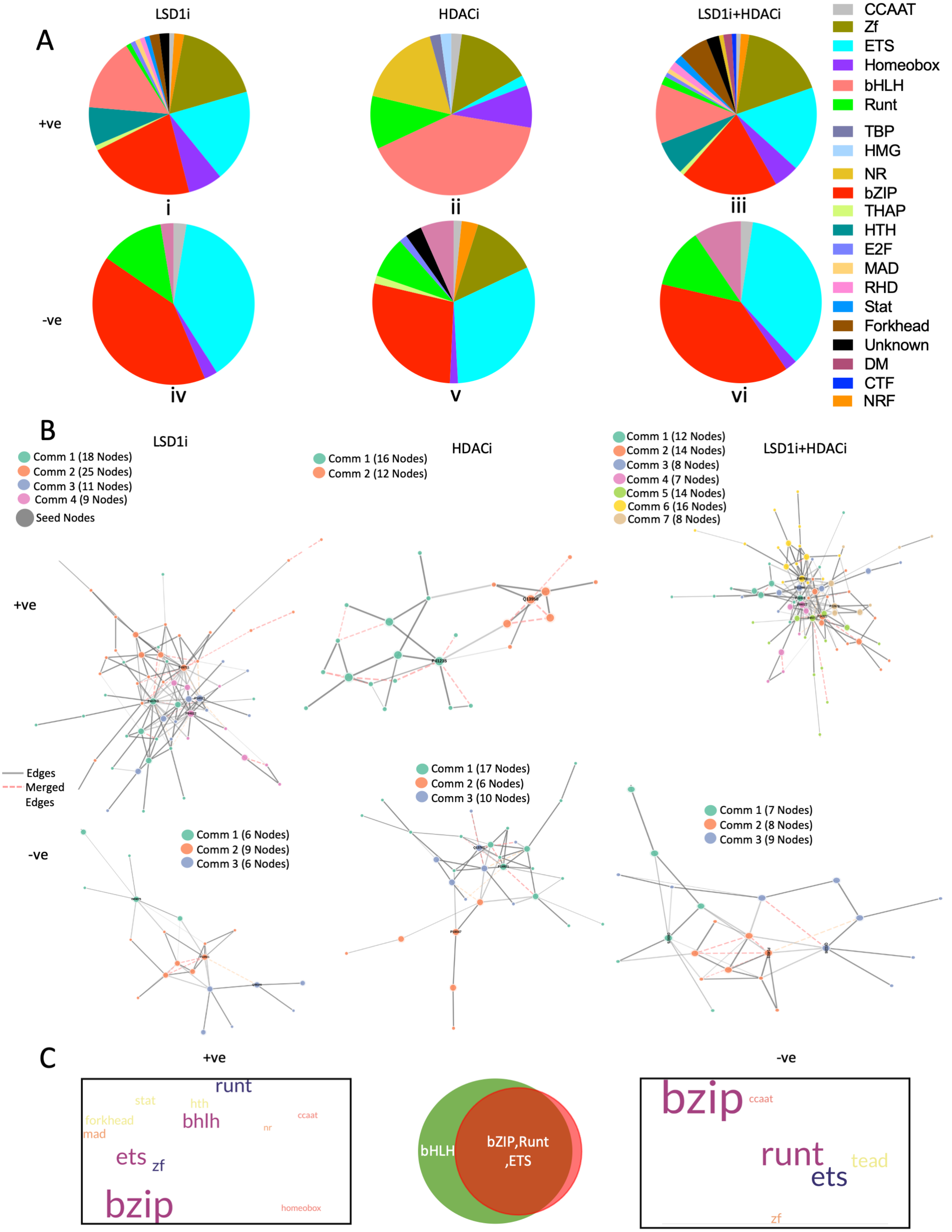

Suppl Figure 4

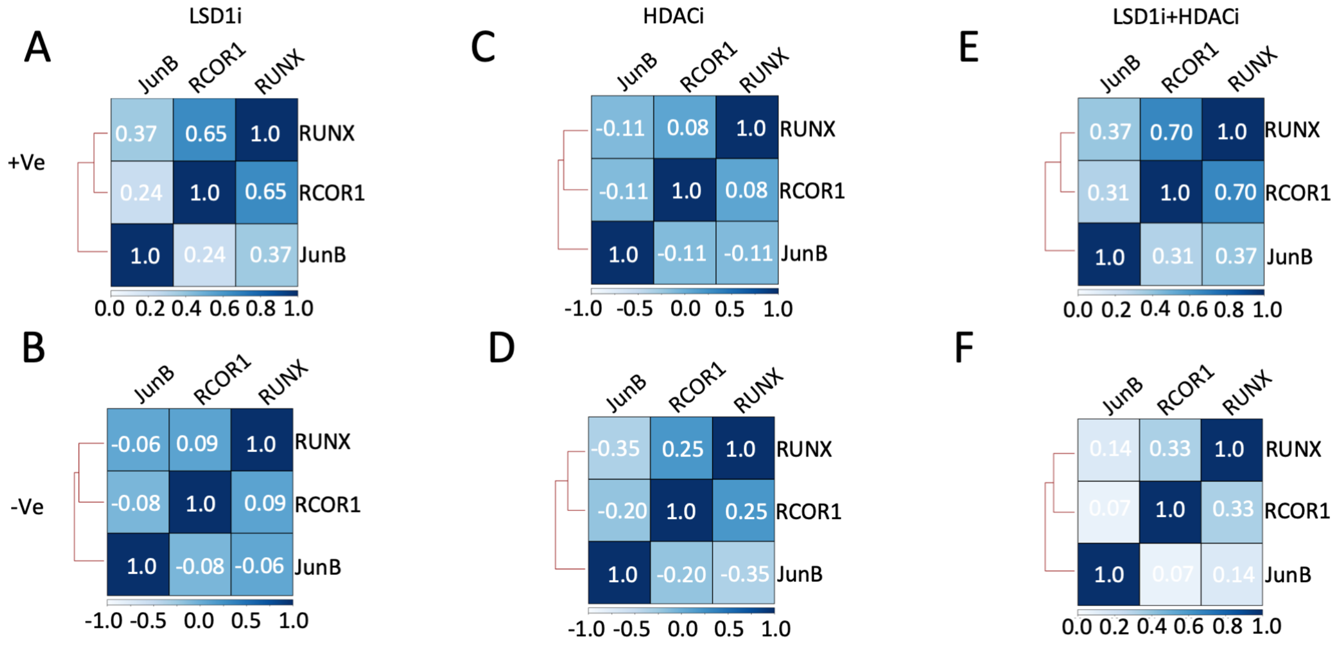

Suppl Figure 5

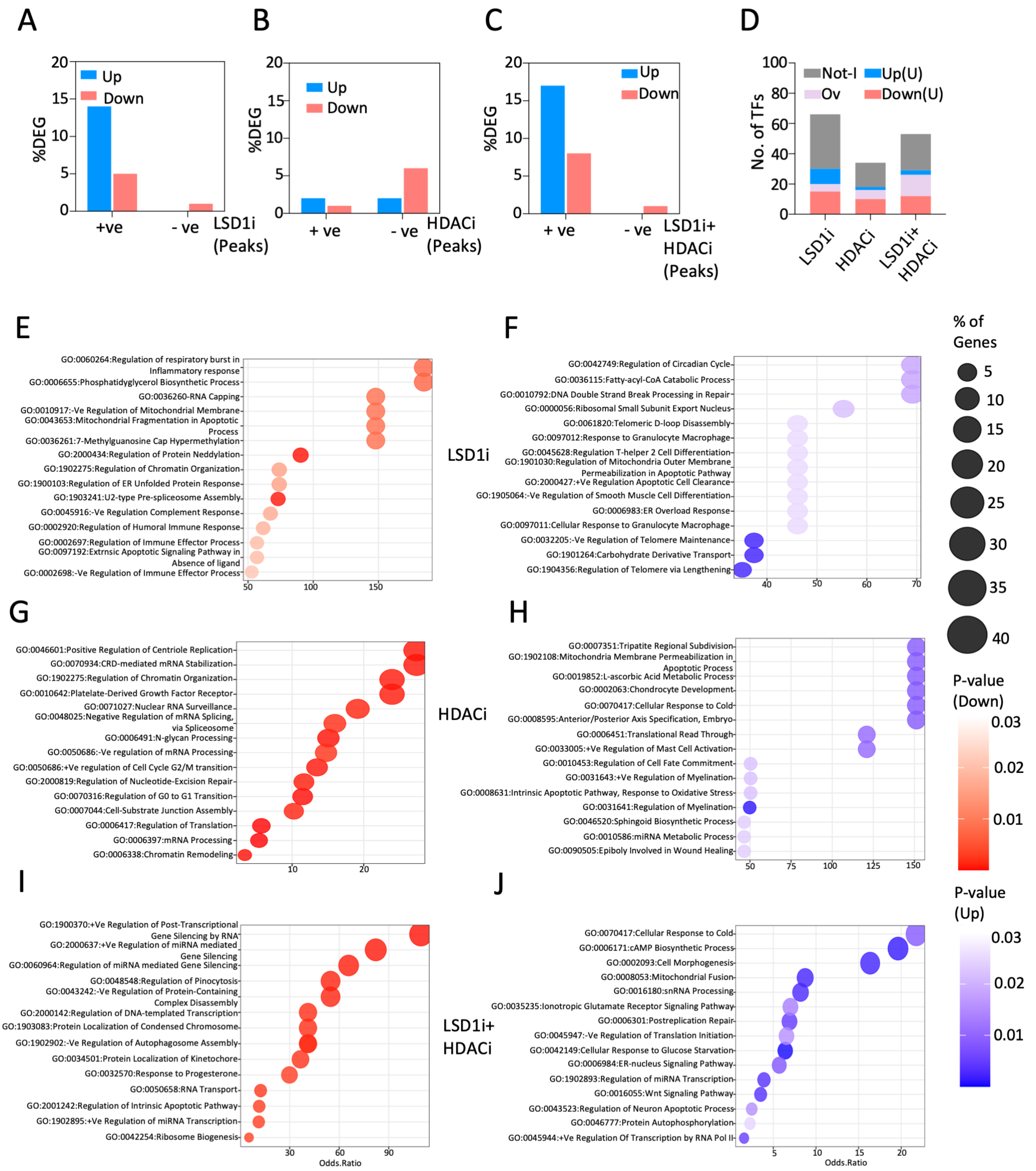

Table S1: List of therapeutic agents and primary targets.

| Compound | Primary Target | Chromatin Accessibility |
|---|---|---|
| JQ-1 | BET bromo-domains (e.g., BRD2, BRD4) | ↓ Zhou et al. 2020 |
| PFI-2 | SETD7 (SET7/9 lysine methyltransferase) | ↓ Barsyte-Lovejoy et al. 2014 |
| WDR5 inhibitors | WDR5 (a core component of MLL/SET1 H3K4 methyltransferase complexes) | ↓ Ruthenburg et al. 2006 |
| UNC0379 | SETD8 Inhibitor | ↑ Julien et al. 2020 |

**Table S2:** Total list of GO:BP terms identified from LSD1i; HDACi and LSD1i plus HDAC1i studies. **A.** up and **B.** down regulated genes represented in Fig 6E-G.

### A. Up Regulated Genes

| GO:BP(LSD1i) | Percentage of Genes | P-value |
|---|---|---|
| Positive Regulation of T-helper Cell Differentiation (GO:0045624) | 16.66666667 | 0.000846976 |
| Negative Regulation of Telomere Maintenance (GO:0032205) | 11.76470588 | 0.001724798 |
| Carbohydrate Derivative Transport (GO:1901264) | 11.76470588 | 0.001724798 |
| Regulation of Telomere Maintenance via Telomere Lengthening (GO:1904356) | 11.11111111 | 0.001935824 |
| Antigen Receptor-Mediated Signaling Pathway (GO:0050851) | 2.631578947 | 0.008398187 |
| Regulation of Telomere Maintenance (GO:0032204) | 3.846153846 | 0.015489228 |
| Nucleotide-Excision Repair (GO:0006289) | 3.773584906 | 0.016059054 |
| Negative Regulation of Transcription by RNA Polymerase II (GO:0000122) | 0.956284153 | 0.017250283 |
| Regulation of Circadian Sleep/Wake Cycle (GO:0042749) | 20 | 0.018118916 |
| Regulation of Circadian Sleep/Wake Cycle, Sleep (GO:0045187) | 20 | 0.018118916 |
| DNA Double-Strand Break Processing Involved in Repair via Single-Strand Annealing (GO:0010792) | 20 | 0.018118916 |
| fatty-acyl-CoA Catabolic Process (GO:0036115) | 20 | 0.018118916 |
| Glycolipid Transport (GO:0046836) | 20 | 0.018118916 |
| Blood Vessel Endothelial Cell Proliferation Involved in Sprouting Angiogenesis (GO:0002043) | 20 | 0.018118916 |
| Medium-Chain fatty-acyl-CoA Metabolic Process (GO:0036112) | 20 | 0.018118916 |
| Regulation of T-Circle Formation (GO:1904429) | 20 | 0.018118916 |
| Membrane Raft Distribution (GO:0031580) | 20 | 0.018118916 |
| Response to DNA Damage Checkpoint Signaling (GO:0072423) | 20 | 0.018118916 |
| Response to cGMP (GO:0070305) | 20 | 0.018118916 |
| Positive Regulation of T-helper 2 Cell Differentiation (GO:0045630) | 20 | 0.018118916 |
| Regulation of cGMP-mediated Signaling (GO:0010752) | 20 | 0.018118916 |
| Double-Strand Break Repair via Single-Strand Annealing (GO:0045002) | 20 | 0.018118916 |
| succinyl-CoA Catabolic Process (GO:1901289) | 20 | 0.018118916 |
| mRNA Stabilization (GO:0048255) | 3.50877193 | 0.018426564 |
| Response to Light Stimulus (GO:0009416) | 3.389830508 | 0.019662263 |
| Phospholipid Transport (GO:0015914) | 3.278688525 | 0.020931855 |
| Positive Regulation of Interleukin-5 Production (GO:0032754) | 16.66666667 | 0.021703663 |
| Negative Regulation of Granulocyte Differentiation (GO:0030853) | 16.66666667 | 0.021703663 |

| | | |
|---|---|---|
| Negative Regulation of Telomere Capping (GO:1904354) | 16.66666667 | 0.021703663 |
| Inner Ear Receptor Cell Differentiation (GO:0060113) | 16.66666667 | 0.021703663 |
| Regulation of Vascular Associated Smooth Muscle Cell Differentiation (GO:1905063) | 16.66666667 | 0.021703663 |
| Ribosomal Small Subunit Export From Nucleus (GO:0000056) | 16.66666667 | 0.021703663 |
| Phosphorylation (GO:0016310) | 1.342281879 | 0.023562311 |
| Regulation of miRNA Transcription (GO:1902893) | 3.03030303 | 0.02425055 |
| Telomeric D-loop Disassembly (GO:0061820) | 14.28571429 | 0.025275502 |
| Establishment of Planar Polarity (GO:0001736) | 14.28571429 | 0.025275502 |
| ER Overload Response (GO:0006983) | 14.28571429 | 0.025275502 |
| + Reg of Mt Outer Membrane Permeabilization Inv in Apoptotic Sgnlng Pway (GO:1901030) | 14.28571429 | 0.025275502 |
| Negative Regulation of Vascular Associated Smooth Muscle Cell Differentiation (GO:1905064) | 14.28571429 | 0.025275502 |
| Cellular Response to Granulocyte Macrophage Colony-Stimulating Factor Stimulus (GO:0097011) | 14.28571429 | 0.025275502 |
| Response to Granulocyte Macrophage Colony-Stimulating Factor (GO:0097012) | 14.28571429 | 0.025275502 |
| Regulation of T-helper 2 Cell Differentiation (GO:0045628) | 14.28571429 | 0.025275502 |
| Positive Regulation of Apoptotic Cell Clearance (GO:2000427) | 14.28571429 | 0.025275502 |
| Endonuc Clvg in ITS1 to Sep SSU-rRNA Fm 5.8S rRNA and LSU-rRNA Fm Tricist rRNA Trnscpt (GO:0000447) | 14.28571429 | 0.025275502 |
| Extrinsic Apoptotic Signaling Pathway (GO:0097191) | 2.941176471 | 0.02563451 |
| Synapse Pruning (GO:0098883) | 12.5 | 0.02883448 |
| Transcription-Coupled Nucleotide-Excision Repair (GO:0006283) | 12.5 | 0.02883448 |
| Positive Regulation of Membrane Invagination (GO:1905155) | 12.5 | 0.02883448 |
| T Cell Apoptotic Process (GO:0070231) | 12.5 | 0.02883448 |
| Inner Ear Receptor Cell Development (GO:0060119) | 12.5 | 0.02883448 |
| Intermembrane Lipid Transfer (GO:0120009) | 12.5 | 0.02883448 |
| Auditory Receptor Cell Stereocilium Organization (GO:0060088) | 12.5 | 0.02883448 |
| Positive Regulation of Vesicle Fusion (GO:0031340) | 12.5 | 0.02883448 |
| succinyl-CoA Metabolic Process (GO:0006104) | 12.5 | 0.02883448 |
| Positive Regulation of Protein Ubiquitination (GO:0031398) | 2.631578947 | 0.031480047 |
| Telomeric Loop Disassembly (GO:0090657) | 11.11111111 | 0.032380641 |
| Positive Regulation of Mitochondrial Membrane Permeability (GO:0035794) | 11.11111111 | 0.032380641 |
| Positive Regulation of Phagocytosis, Engulfment (GO:0060100) | 11.11111111 | 0.032380641 |
| Neuron Remodeling (GO:0016322) | 11.11111111 | 0.032380641 |
| Auditory Receptor Cell Morphogenesis (GO:0002093) | 11.11111111 | 0.032380641 |

| | | |
|---|---|---|
| Collagen Biosynthetic Process (GO:0032964) | 11.11111111 | 0.032380641 |
| Positive Regulation of Protein Modification by Small Protein Conjugation or Removal (GO:1903322) | 2.53164557 | 0.033795318 |
| T-Circle Formation (GO:0090656) | 10 | 0.035914031 |
| Positive Regulation of Interleukin-13 Production (GO:0032736) | 10 | 0.035914031 |
| Telomere Maintenance via Telomere Trimming (GO:0090737) | 10 | 0.035914031 |
| Positive Regulation of Lipid Localization (GO:1905954) | 10 | 0.035914031 |
| Negative Regulation of Interferon-Beta Production (GO:0032688) | 10 | 0.035914031 |
| Transposable Element Silencing by piRNA-mediated DNA Methylation (GO:0141196) | 10 | 0.035914031 |
| Regulation of Granulocyte Differentiation (GO:0030852) | 10 | 0.035914031 |
| Formation of Extrachromosomal Circular DNA (GO:0001325) | 10 | 0.035914031 |
| Gene Silencing by piRNA-directed DNA Methylation (GO:0141176) | 10 | 0.035914031 |
| Negative Regulation of Smooth Muscle Cell Differentiation (GO:0051151) | 10 | 0.035914031 |
| Regulation of Phagocytosis, Engulfment (GO:0060099) | 10 | 0.035914031 |
| Regulation of Telomere Capping (GO:1904353) | 10 | 0.035914031 |
| Positive Regulation of T-helper 17 Cell Differentiation (GO:2000321) | 10 | 0.035914031 |
| Regulation of T-helper 17 Cell Differentiation (GO:2000319) | 10 | 0.035914031 |
| Detection of Mechanical Stimulus Involved in Sensory Perception of Sound (GO:0050910) | 10 | 0.035914031 |
| Positive Regulation of Cellular Senescence (GO:2000774) | 10 | 0.035914031 |
| - Reg of Adenylate Cyclase-Activating GPCR Sgnlng Pway (GO:0106072) | 10 | 0.035914031 |
| Regulation of Interleukin-1 Beta Production (GO:0032651) | 2.43902439 | 0.036175169 |
| Carbohydrate Metabolic Process (GO:0005975) | 2.352941176 | 0.038617957 |
| Negative Regulation of Epithelial Cell Apoptotic Process (GO:1904036) | 9.090909091 | 0.039434695 |
| Tricarboxylic Acid Metabolic Process (GO:0072350) | 9.090909091 | 0.039434695 |
| Positive Regulation of Toll-Like Receptor 4 Signaling Pathway (GO:0034145) | 9.090909091 | 0.039434695 |
| Regulation of Programmed Necrotic Cell Death (GO:0062098) | 9.090909091 | 0.039434695 |
| cGMP Metabolic Process (GO:0046068) | 9.090909091 | 0.039434695 |
| Monocyte Differentiation (GO:0030224) | 9.090909091 | 0.039434695 |
| T Cell Receptor Signaling Pathway (GO:0050852) | 2.272727273 | 0.041122067 |
| Protein Stabilization (GO:0050821) | 1.41509434 | 0.042516677 |
| Regulation of Interleukin-5 Production (GO:0032674) | 8.333333333 | 0.042942678 |
| Negative Regulation of Vascular Permeability (GO:0043116) | 8.333333333 | 0.042942678 |
| Positive Regulation of Vascular Permeability (GO:0043117) | 8.333333333 | 0.042942678 |

| | | |
|---|---|---|
| Branching Involved in Blood Vessel Morphogenesis (GO:0001569) | 8.333333333 | 0.042942678 |
| Regulation of Retrograde Protein Transport, ER to Cytosol (GO:1904152) | 8.333333333 | 0.042942678 |
| Phospholipid Efflux (GO:0033700) | 8.333333333 | 0.042942678 |
| Mitochondrial Respiratory Chain Complex III Assembly (GO:0034551) | 8.333333333 | 0.042942678 |
| Respiratory Chain Complex III Assembly (GO:0017062) | 8.333333333 | 0.042942678 |
| Retina Homeostasis (GO:0001895) | 8.333333333 | 0.042942678 |
| Regulation of Adenylate Cyclase-Activating G Protein-Coupled Receptor Signaling Pathway (GO:0106070) | 8.333333333 | 0.042942678 |
| ERAD Pathway (GO:0036503) | 2.197802198 | 0.04368591 |
| Sensory Perception of Mechanical Stimulus (GO:0050954) | 2.173913043 | 0.044553527 |
| Regulation of Protein Metabolic Process (GO:0051246) | 2.150537634 | 0.045427551 |
| Inner Ear Receptor Cell Stereocilium Organization (GO:0060122) | 7.692307692 | 0.046438025 |
| Positive Regulation of Type 2 Immune Response (GO:0002830) | 7.692307692 | 0.046438025 |
| calcineurin-NFAT Signaling Cascade (GO:0033173) | 7.692307692 | 0.046438025 |
| Positive Regulation of T-helper 17 Type Immune Response (GO:2000318) | 7.692307692 | 0.046438025 |
| Endonucleolytic Clvg of Tricistronic rRNA Trnscpt (GO:0000479) | 7.692307692 | 0.046438025 |
| Protein Phosphorylation (GO:0006468) | 1.072386059 | 0.047488575 |
| B Cell Activation (GO:0042113) | 2.083333333 | 0.04808749 |
| Sensory Perception of Sound (GO:0007605) | 2.083333333 | 0.04808749 |
| Protein Modification Process (GO:0036211) | 0.917431193 | 0.04888689 |
| Regulation of Dephosphorylation (GO:0035303) | 7.142857143 | 0.049920782 |
| Negative Regulation of Toll-Like Receptor Signaling Pathway (GO:0034122) | 7.142857143 | 0.049920782 |
| Cellular Response to Estrogen Stimulus (GO:0071391) | 7.142857143 | 0.049920782 |
| Purine-Containing Compound Catabolic Process (GO:0072523) | 7.142857143 | 0.049920782 |
| Resolution of Meiotic Recombination Intermediates (GO:0000712) | 7.142857143 | 0.049920782 |
| Coenzyme A Metabolic Process (GO:0015936) | 7.142857143 | 0.049920782 |
| Retinoic Acid Receptor Signaling Pathway (GO:0048384) | 7.142857143 | 0.049920782 |

| **GO:BP(HDAC1i)** | **Percentage of Genes** | **P-value** |
|---|---|---|
| Regulation of Myelination (GO:0031641) | 7.407407407 | 0.000958686 |
| Negative Regulation of Growth (GO:0045926) | 2.608695652 | 0.000973134 |
| Negative Regulation of Cell Growth (GO:0030308) | 2.479338843 | 0.001127166 |
| Apoptotic Mitochondrial Changes (GO:0008637) | 6.25 | 0.001347535 |
| Positive Regulation of Establishment of Protein Localization to Mitochondrion (GO:1903749) | 6.25 | 0.001347535 |

| | | |
|---|---|---|
| NADH Dehydrogenase Complex Assembly (GO:0010257) | 5.714285714 | 0.001611348 |
| Mitochondrial Respiratory Chain Complex I Assembly (GO:0032981) | 5.714285714 | 0.001611348 |
| Positive Regulation of Cytokine Production Involved in Immune Response (GO:0002720) | 5.714285714 | 0.001611348 |
| Nuclear Receptor-Mediated Steroid Hormone Signaling Pathway (GO:0030518) | 4.166666667 | 0.003012867 |
| DNA-templated Transcription Initiation (GO:0006352) | 3.225806452 | 0.004976305 |
| Positive Regulation of Intracellular Protein Transport (GO:0090316) | 2.898550725 | 0.006128005 |
| Regulation of Cell Growth (GO:0001558) | 1.351351351 | 0.006265621 |
| Positive Regulation of Protein Localization to Nucleus (GO:1900182) | 2.666666667 | 0.007202645 |
| Tripartite Regional Subdivision (GO:0007351) | 20 | 0.0084719 |
| L-ascorbic Acid Metabolic Process (GO:0019852) | 20 | 0.0084719 |
| Cellular Response to Cold (GO:0070417) | 20 | 0.0084719 |
| Chondrocyte Development (GO:0002063) | 20 | 0.0084719 |
| Regulation of Mitochondrial Membrane Permeability Involved in Apoptotic Process (GO:1902108) | 20 | 0.0084719 |
| Anterior/Posterior Axis Specification, Embryo (GO:0008595) | 20 | 0.0084719 |
| Positive Regulation of Tumor Necrosis Factor Production (GO:0032760) | 2.43902439 | 0.008556135 |
| Positive Regulation of Tumor Necrosis Factor Superfamily Cytokine Production (GO:1903557) | 2.352941176 | 0.009168483 |
| Positive Regulation of Canonical Wnt Signaling Pathway (GO:0090263) | 2.247191011 | 0.010014594 |
| Translational Readthrough (GO:0006451) | 16.66666667 | 0.010157909 |
| Positive Regulation of Mast Cell Activation (GO:0033005) | 16.66666667 | 0.010157909 |
| Positive Regulation of Mast Cell Degranulation (GO:0043306) | 16.66666667 | 0.010157909 |
| PERK-mediated Unfolded Protein Response (GO:0036499) | 16.66666667 | 0.010157909 |
| Mitochondrial Fragmentation Involved in Apoptotic Process (GO:0043653) | 16.66666667 | 0.010157909 |
| Negative Regulation of Nervous System Process (GO:0031645) | 16.66666667 | 0.010157909 |
| Regulation of Rab Protein Signal Transduction (GO:0032483) | 16.66666667 | 0.010157909 |
| Selenocysteine Incorporation (GO:0001514) | 16.66666667 | 0.010157909 |
| - Reg of Blood Vessel Endothelial Cell Proliferation Inv in Sprouting Angiogenesis (GO:1903588) | 16.66666667 | 0.010157909 |
| Mitochondrial Respiratory Chain Complex Assembly (GO:0033108) | 2.222222222 | 0.010231366 |
| Regulation of Gene Expression (GO:0010468) | 0.532386868 | 0.010934438 |
| Establishment of Golgi Localization (GO:0051683) | 14.28571429 | 0.011841135 |
| ER Overload Response (GO:0006983) | 14.28571429 | 0.011841135 |
| Hair Cell Differentiation (GO:0035315) | 14.28571429 | 0.011841135 |
| Monosaccharide Metabolic Process (GO:0005996) | 14.28571429 | 0.011841135 |

| | | |
|---|---|---|
| siRNA Processing (GO:0030422) | 14.28571429 | 0.011841135 |
| Negative Regulation of Cellular Process (GO:0048523) | 0.753295669 | 0.012127379 |
| RNA Processing (GO:0006396) | 1.030927835 | 0.013076574 |
| Regulation of Opsin-Mediated Signaling Pathway (GO:0022400) | 12.5 | 0.013521585 |
| mRNA Pseudouridine Synthesis (GO:1990481) | 11.11111111 | 0.01519926 |
| Regulation of Translation in Response to Endoplasmic Reticulum Stress (GO:0036490) | 11.11111111 | 0.01519926 |
| Positive Regulation of Wnt Signaling Pathway (GO:0030177) | 1.754385965 | 0.016042396 |
| Positive Regulation of Interleukin-13 Production (GO:0032736) | 10 | 0.016874167 |
| Negative Regulation of Interferon-Beta Production (GO:0032688) | 10 | 0.016874167 |
| miRNA Catabolic Process (GO:0010587) | 10 | 0.016874167 |
| Heterochromatin Organization (GO:0070828) | 10 | 0.016874167 |
| Positive Regulation of Cellular Senescence (GO:2000774) | 10 | 0.016874167 |
| Regulation of Ceramide Biosynthetic Process (GO:2000303) | 10 | 0.016874167 |
| Positive Regulation of Cytokine Production (GO:0001819) | 0.923076923 | 0.017538066 |
| Transcription Initiation at RNA Polymerase II Promoter (GO:0006367) | 1.62601626 | 0.018510881 |
| Regulation of Establishment of Protein Localization to Mitochondrion (GO:1903747) | 9.090909091 | 0.01854631 |
| Golgi Inheritance (GO:0048313) | 9.090909091 | 0.01854631 |
| Regulation of Mast Cell Degranulation (GO:0043304) | 9.090909091 | 0.01854631 |
| Insulin-Like Growth Factor Receptor Signaling Pathway (GO:0048009) | 9.090909091 | 0.01854631 |
| Diol Metabolic Process (GO:0034311) | 9.090909091 | 0.01854631 |
| Sphingoid Metabolic Process (GO:0046519) | 9.090909091 | 0.01854631 |
| Progesterone Receptor Signaling Pathway (GO:0050847) | 9.090909091 | 0.01854631 |
| Protein K29-linked Ubiquitination (GO:0035519) | 9.090909091 | 0.01854631 |
| Regulation of Mitotic Cell Cycle (GO:0007346) | 1.612903226 | 0.018794509 |
| RNA Biosynthetic Process (GO:0032774) | 1.5625 | 0.019947409 |
| Endothelium Development (GO:0003158) | 8.333333333 | 0.020215692 |
| Positive Regulation of Leukocyte Degranulation (GO:0043302) | 8.333333333 | 0.020215692 |
| Regulation of ATP Biosynthetic Process (GO:2001169) | 8.333333333 | 0.020215692 |
| Regulation of Tumor Necrosis Factor Production (GO:0032680) | 1.526717557 | 0.020831206 |
| Cellular Response to Arsenic-Containing Substance (GO:0071243) | 7.692307692 | 0.021882319 |
| Pseudouridine Synthesis (GO:0001522) | 7.692307692 | 0.021882319 |
| Positive Regulation of Myelination (GO:0031643) | 7.692307692 | 0.021882319 |
| Intrinsic Apoptotic Signaling Pathway in Response to Oxidative Stress (GO:0008631) | 7.692307692 | 0.021882319 |
| Regulation of Cell Fate Commitment (GO:0010453) | 7.692307692 | 0.021882319 |

| | | |
|---|---|---|
| Positive Regulation of Macromolecule Biosynthetic Process (GO:0010557) | 0.845070423 | 0.022101454 |
| Epiboly Involved in Wound Healing (GO:0090505) | 7.142857143 | 0.023546194 |
| miRNA Metabolic Process (GO:0010586) | 7.142857143 | 0.023546194 |
| Sphingoid Biosynthetic Process (GO:0046520) | 7.142857143 | 0.023546194 |
| Sphingosine Biosynthetic Process (GO:0046512) | 7.142857143 | 0.023546194 |
| Negative Regulation of Cell Migration Involved in Sprouting Angiogenesis (GO:0090051) | 7.142857143 | 0.023546194 |
| Negative Regulation of Gene Expression (GO:0010629) | 0.824175824 | 0.023586015 |
| Regulation of Response to Endoplasmic Reticulum Stress (GO:1905897) | 6.666666667 | 0.025207322 |
| mRNA Modification (GO:0016556) | 6.666666667 | 0.025207322 |
| Positive Regulation of Membrane Protein Ectodomain Proteolysis (GO:0051044) | 6.666666667 | 0.025207322 |
| Positive Regulation of Mitochondrial Translation (GO:0070131) | 6.666666667 | 0.025207322 |
| Regulation of Interleukin-13 Production (GO:0032656) | 6.666666667 | 0.025207322 |
| Cellular Response to Misfolded Protein (GO:0071218) | 6.666666667 | 0.025207322 |
| Regulation of Intrinsic Apoptotic Signaling Pathway in Response to DNA Damage (GO:1902229) | 6.666666667 | 0.025207322 |
| Regulation of ATP Metabolic Process (GO:1903578) | 6.666666667 | 0.025207322 |
| Regulation of Peptidase Activity (GO:0052547) | 6.666666667 | 0.025207322 |
| Regulation of Cell Fate Specification (GO:0042659) | 6.666666667 | 0.025207322 |
| Regulation of Cell Cycle Process (GO:0010564) | 1.369863014 | 0.025489528 |
| Negative Regulation of DNA-templated Transcription (GO:0045892) | 0.497017893 | 0.026392476 |
| Regulation of DNA Damage Checkpoint (GO:2000001) | 6.25 | 0.026865707 |
| Positive Regulation of Stress-Activated MAPK Cascade (GO:0032874) | 6.25 | 0.026865707 |
| Positive Regulation of Stress-Activated Protein Kinase Signaling Cascade (GO:0070304) | 6.25 | 0.026865707 |
| Reg of Blood Vessel Endothelial Cell Proliferation Inv in Sprouting Angiogenesis (GO:1903587) | 6.25 | 0.026865707 |
| Negative Regulation of Fibroblast Proliferation (GO:0048147) | 5.882352941 | 0.028521354 |
| Regulation of Mitochondrial Membrane Permeability (GO:0046902) | 5.882352941 | 0.028521354 |
| Contractile Actin Filament Bundle Assembly (GO:0030038) | 5.882352941 | 0.028521354 |
| Embryonic Axis Specification (GO:0000578) | 5.882352941 | 0.028521354 |
| Stress Fiber Assembly (GO:0043149) | 5.882352941 | 0.028521354 |
| Regulation of Small GTPase Mediated Signal Transduction (GO:0051056) | 1.27388535 | 0.029150243 |
| Protein Branched Polyubiquitination (GO:0141198) | 5.555555556 | 0.030174267 |
| Estrogen Receptor Signaling Pathway (GO:0030520) | 5.555555556 | 0.030174267 |
| Negative Regulation of Translational Initiation (GO:0045947) | 5.555555556 | 0.030174267 |

| | | |
|---|---|---|
| Regulation of Autophagy of Mitochondrion (GO:1903146) | 5.555555556 | 0.030174267 |
| Regulation of Oxidoreductase Activity (GO:0051341) | 5.555555556 | 0.030174267 |
| Diol Biosynthetic Process (GO:0034312) | 5.555555556 | 0.030174267 |
| Maintenance of Protein Localization in Organelle (GO:0072595) | 5.263157895 | 0.031824451 |
| Regulation of Nuclear Division (GO:0051783) | 5.263157895 | 0.031824451 |
| Regulation of Endopeptidase Activity (GO:0052548) | 5 | 0.033471909 |
| Maintenance of Protein Location in Nucleus (GO:0051457) | 5 | 0.033471909 |
| Regulation of T Cell Cytokine Production (GO:0002724) | 5 | 0.033471909 |
| Sphingosine Metabolic Process (GO:0006670) | 5 | 0.033471909 |
| DNA Repair-Dependent Chromatin Remodeling (GO:0140861) | 4.761904762 | 0.035116647 |
| Cellular Response to Estradiol Stimulus (GO:0071392) | 4.761904762 | 0.035116647 |
| Positive Regulation of Nervous System Process (GO:0031646) | 4.545454545 | 0.036758668 |
| Anterior/Posterior Axis Specification (GO:0009948) | 4.545454545 | 0.036758668 |
| Regulation of Biosynthetic Process (GO:0009889) | 4.545454545 | 0.036758668 |
| Post-Transcriptional Gene Silencing (GO:0016441) | 4.545454545 | 0.036758668 |
| Positive Regulation of Gene Expression (GO:0010628) | 0.688073394 | 0.037382267 |
| Neuron Differentiation (GO:0030182) | 1.104972376 | 0.037812019 |
| Regulatory ncRNA-mediated Post-Transcriptional Gene Silencing (GO:0035194) | 4.347826087 | 0.038397977 |
| Positive Regulation of T Cell Cytokine Production (GO:0002726) | 4.347826087 | 0.038397977 |
| Regulation of Mitochondrial Translation (GO:0070129) | 4.347826087 | 0.038397977 |
| Positive Regulation of Amide Metabolic Process (GO:0034250) | 4.347826087 | 0.038397977 |
| Negative Regulation of Smoothened Signaling Pathway (GO:0045879) | 4.347826087 | 0.038397977 |
| Negative Regulation of Intrinsic Apoptotic Signaling Pathway in Response to DNA Damage (GO:1902230) | 4 | 0.041668475 |
| Chaperone Cofactor-Dependent Protein Refolding (GO:0051085) | 4 | 0.041668475 |
| Regulation of Membrane Protein Ectodomain Proteolysis (GO:0051043) | 4 | 0.041668475 |
| Negative Regulation of Catalytic Activity (GO:0043086) | 4 | 0.041668475 |
| Nucleic Acid Metabolic Process (GO:0090304) | 4 | 0.041668475 |
| Reg of Endoplasmic Reticulum Stress-Induced Intrinsic Apoptotic Sgnlng Pway (GO:1902235) | 3.846153846 | 0.043299673 |
| Response to Endoplasmic Reticulum Stress (GO:0034976) | 1.020408163 | 0.043666329 |
| Regulation of Reactive Oxygen Species Biosynthetic Process (GO:1903426) | 3.703703704 | 0.044928177 |
| Regulation of Cytokine Production Involved in Immune Response (GO:0002718) | 3.703703704 | 0.044928177 |
| ER-nucleus Signaling Pathway (GO:0006984) | 3.703703704 | 0.044928177 |

| | | |
|---|---|---|
| Protein Quality Control for Misfolded or Incompletely Synthesized Proteins (GO:0006515) | 3.703703704 | 0.044928177 |
| Wound Healing, Spreading of Cells (GO:0044319) | 3.703703704 | 0.044928177 |
| Positive Regulation of Vascular Endothelial Growth Factor Production (GO:0010575) | 3.703703704 | 0.044928177 |
| Negative Regulation of RNA Biosynthetic Process (GO:1902679) | 0.638297872 | 0.04507002 |
| Transcription by RNA Polymerase II (GO:0006366) | 1 | 0.045281573 |
| DNA-templated Transcription (GO:0006351) | 0.985221675 | 0.04650752 |
| Phospholipid Biosynthetic Process (GO:0008654) | 3.571428571 | 0.046553989 |
| Regulation of Mitophagy (GO:1901524) | 3.571428571 | 0.046553989 |
| Positive Regulation of Viral Genome Replication (GO:0045070) | 3.571428571 | 0.046553989 |
| Regulation of Vascular Endothelial Growth Factor Production (GO:0010574) | 3.448275862 | 0.048177116 |
| Positive Regulation of Protein Import Into Nucleus (GO:0042307) | 3.448275862 | 0.048177116 |
| Positive Regulation of Protein Targeting to Mitochondrion (GO:1903955) | 3.448275862 | 0.048177116 |
| Positive Regulation of Peptidyl-Serine Phosphorylation (GO:0033138) | 3.333333333 | 0.04979756 |
| Protein Stabilization (GO:0050821) | 0.943396226 | 0.05 |

| **GO:BP(LSD1i plus HDAC1i)** | **Percentage of Genes** | **P-value** |
|---|---|---|
| Cellular Response to Glucose Starvation (GO:0042149) | 16.27907 | 0.00026 |
| Auditory Receptor Cell Stereocilium Organization (GO:0060088) | 37.5 | 0.00132 |
| cAMP Biosynthetic Process (GO:0006171) | 37.5 | 0.00132 |
| Peptidyl-Tyrosine Phosphorylation (GO:0018108) | 12.06897 | 0.00162 |
| Ephrin Receptor Signaling Pathway (GO:0048013) | 13.95349 | 0.00163 |
| Auditory Receptor Cell Morphogenesis (GO:0002093) | 33.33333 | 0.00193 |
| Protein Phosphorylation (GO:0006468) | 5.898123 | 0.00196 |
| Mitochondrial Fusion (GO:0008053) | 21.05263 | 0.00212 |
| snRNA Processing (GO:0016180) | 20 | 0.00259 |
| Positive Regulation of miRNA Transcription (GO:1902895) | 12.76596 | 0.0026 |
| Endoplasmic Reticulum Unfolded Protein Response (GO:0030968) | 12.5 | 0.00289 |
| Wnt Signaling Pathway (GO:0016055) | 9.638554 | 0.00329 |
| Regulation of miRNA Transcription (GO:1902893) | 10.60606 | 0.00343 |
| snRNA Metabolic Process (GO:0016073) | 18.18182 | 0.00372 |
| Cellular Response to Vascular Endothelial Growth Factor Stimulus (GO:0035924) | 13.88889 | 0.00406 |
| Autophagosome Organization (GO:1905037) | 10.14493 | 0.0044 |
| Postreplication Repair (GO:0006301) | 17.3913 | 0.0044 |

| | | |
|---|---|---|
| cAMP Metabolic Process (GO:0046058) | 25 | 0.00474 |
| Positive Regulation of miRNA Metabolic Process (GO:2000630) | 11.32075 | 0.00478 |
| Positive Regulation of Transcription by RNA Polymerase II (GO:0045944) | 4.476094 | 0.00482 |
| Inner Ear Receptor Cell Stereocilium Organization (GO:0060122) | 23.07692 | 0.00602 |
| Protein Modification Process (GO:0036211) | 4.954128 | 0.00721 |
| Cellular Response to Unfolded Protein (GO:0034620) | 10.34483 | 0.00744 |
| ER-nucleus Signaling Pathway (GO:0006984) | 14.81481 | 0.00795 |
| Cellular Response to Cold (GO:0070417) | 40 | 0.00835 |
| Ventricular Cardiac Muscle Cell Differentiation (GO:0055012) | 40 | 0.00835 |
| Neuron Migration (GO:0001764) | 10 | 0.00875 |
| Positive Regulation of DNA-templated Transcription (GO:0045893) | 4.160126 | 0.00881 |
| Antiviral Innate Immune Response (GO:0140374) | 9.677419 | 0.01023 |
| Organelle Fusion (GO:0048284) | 13.7931 | 0.01026 |
| Response to UV-B (GO:0010224) | 18.75 | 0.01104 |
| Positive Reg of Myeloid Leukocyte Cytokine Production Inv in Imm Resp (GO:0061081) | 18.75 | 0.01104 |
| Peptidyl-Tyrosine Modification (GO:0018212) | 10.86957 | 0.01157 |
| IRE1-mediated Unfolded Protein Response (GO:0036498) | 33.33333 | 0.01228 |
| Regulation of Type B Pancreatic Cell Proliferation (GO:0061469) | 33.33333 | 0.01228 |
| Desmosome Assembly (GO:0002159) | 33.33333 | 0.01228 |
| Positive Regulation of Mast Cell Activation (GO:0033005) | 33.33333 | 0.01228 |
| Cellular Response to Nitrogen Compound (GO:1901699) | 6.508876 | 0.01262 |
| Fat Cell Differentiation (GO:0045444) | 9.230769 | 0.01276 |
| Protein Autoubiquitination (GO:0051865) | 9.230769 | 0.01276 |
| Regulation of Dendritic Spine Morphogenesis (GO:0061001) | 12.90323 | 0.01298 |
| Ionotropic Glutamate Receptor Signaling Pathway (GO:0035235) | 17.64706 | 0.01311 |
| Hexose Metabolic Process (GO:0019318) | 10.41667 | 0.01377 |
| Regulation of Postsynapse Organization (GO:0099175) | 12.5 | 0.01449 |
| Golgi Organization (GO:0007030) | 7.03125 | 0.01451 |
| Endomembrane System Organization (GO:0010256) | 6.122449 | 0.01477 |
| Regulation of Neuron Apoptotic Process (GO:0043523) | 6.976744 | 0.0152 |
| Cyclic Nucleotide Biosynthetic Process (GO:0009190) | 16.66667 | 0.01539 |
| Negative Regulation of Translational Initiation (GO:0045947) | 16.66667 | 0.01539 |
| Positive Regulation of Protein Kinase Activity (GO:0045860) | 7.954545 | 0.01602 |
| Cell Surface Receptor Protein Tyrosine Kinase Signaling Pathway (GO:0007169) | 5.405405 | 0.01645 |

| | | |
|---|---|---|
| ER Overload Response (GO:0006983) | 28.57143 | 0.01685 |
| Dense Core Granule Cytoskeletal Transport (GO:0099519) | 28.57143 | 0.01685 |
| Negative Regulation of Stress-Activated Protein Kinase Signaling Cascade (GO:0070303) | 28.57143 | 0.01685 |
| Nonribosomal Peptide Biosynthetic Process (GO:0019184) | 28.57143 | 0.01685 |
| Limb Morphogenesis (GO:0035108) | 15.78947 | 0.01788 |
| piRNA Processing (GO:0034587) | 15.78947 | 0.01788 |
| Regulation of p38MAPK Cascade (GO:1900744) | 11.42857 | 0.01967 |
| Positive Regulation of Cytokine Production Involved in Immune Response (GO:0002720) | 11.42857 | 0.01967 |
| Autophagosome Assembly (GO:0000045) | 8.333333 | 0.02031 |
| Endochondral Ossification (GO:0001958) | 25 | 0.02203 |
| Nucleic Acid Transport (GO:0050657) | 25 | 0.02203 |
| Reelin-Mediated Signaling Pathway (GO:0038026) | 25 | 0.02203 |
| COPII Vesicle Coating (GO:0048208) | 14.28571 | 0.02349 |
| Vesicle Targeting, Rough ER to cis-Golgi (GO:0048207) | 14.28571 | 0.02349 |
| Macroautophagy (GO:0016236) | 6.17284 | 0.02354 |
| Protein Autophosphorylation (GO:0046777) | 6.428571 | 0.02448 |
| Cellular Response to Alcohol (GO:0097306) | 8.928571 | 0.02535 |
| Positive Regulation of Interleukin-1 Beta Production (GO:0032731) | 8.928571 | 0.02535 |
| Cellular Response to Starvation (GO:0009267) | 6.382979 | 0.02549 |
| Regulation of Synapse Organization (GO:0050807) | 10.52632 | 0.02588 |
| Regulation of DNA-templated Transcription Elongation (GO:0032784) | 10.52632 | 0.02588 |
| Positive Regulation of Protein Modification Process (GO:0031401) | 7.142857 | 0.02718 |
| Regulation of Translation in Response to Endoplasmic Reticulum Stress (GO:0036490) | 22.22222 | 0.02778 |
| Glycophagy (GO:0061723) | 22.22222 | 0.02778 |
| Intestinal Cholesterol Absorption (GO:0030299) | 22.22222 | 0.02778 |
| Positive Regulation of Macrophage Cytokine Production (GO:0060907) | 22.22222 | 0.02778 |
| Regulation of Calcium Ion Import Across Plasma Membrane (GO:1905664) | 22.22222 | 0.02778 |
| Vacuolar Transport (GO:0007034) | 10.25641 | 0.02817 |
| Cellular Response to Oxidative Stress (GO:0034599) | 6.61157 | 0.02845 |
| Positive Regulation of Autophagy (GO:0010508) | 7.070707 | 0.02853 |
| Generation of Neurons (GO:0048699) | 5.729167 | 0.02932 |
| Translesion Synthesis (GO:0019985) | 13.04348 | 0.02994 |
| Negative Regulation of Smoothened Signaling Pathway (GO:0045879) | 13.04348 | 0.02994 |
| Positive Regulation of Amide Metabolic Process (GO:0034250) | 13.04348 | 0.02994 |

| | | |
|---|---|---|
| Positive Regulation of Amyloid Precursor Protein Catabolic Process (GO:1902993) | 13.04348 | 0.02994 |
| Regulation of Transcription Elongation by RNA Polymerase II (GO:0034243) | 7.594937 | 0.03042 |
| Negative Regulation of Transcription by RNA Polymerase II (GO:0000122) | 4.234973 | 0.03206 |
| Negative Regulation of Translation (GO:0017148) | 6.862745 | 0.03283 |
| Endothelial Cell Development (GO:0001885) | 12.5 | 0.03348 |
| Positive Regulation of Striated Muscle Cell Differentiation (GO:0051155) | 12.5 | 0.03348 |
| Negative Regulation of Cell Cycle (GO:0045786) | 7.407407 | 0.03383 |
| Negative Regulation of Neuron Apoptotic Process (GO:0043524) | 7.407407 | 0.03383 |
| Desmosome Organization (GO:0002934) | 20 | 0.03405 |
| Embryonic Appendage Morphogenesis (GO:0035113) | 20 | 0.03405 |
| Growth Hormone Receptor Signaling Pathway via JAK-STAT (GO:0060397) | 20 | 0.03405 |
| Microtubule Anchoring at Centrosome (GO:0034454) | 20 | 0.03405 |
| Negative Regulation of Interferon-Beta Production (GO:0032688) | 20 | 0.03405 |
| Neuron Projection Fasciculation (GO:0106030) | 20 | 0.03405 |
| Positive Regulation of Gluconeogenesis (GO:0045722) | 20 | 0.03405 |
| Positive Regulation of Interleukin-13 Production (GO:0032736) | 20 | 0.03405 |
| Regulation of Ceramide Biosynthetic Process (GO:2000303) | 20 | 0.03405 |
| Phosphorylation (GO:0016310) | 5.033557 | 0.03429 |
| Regulation of Interleukin-1 Beta Production (GO:0032651) | 7.317073 | 0.03562 |
| Regulation of Anatomical Structure Morphogenesis (GO:0022603) | 6.299213 | 0.03642 |
| Non-Canonical NF-kappaB Signal Transduction (GO:0038061) | 12 | 0.03723 |
| Regulation of Protein Kinase Activity (GO:0045859) | 7.228916 | 0.03746 |
| Negative Regulation of Organelle Assembly (GO:1902116) | 9.302326 | 0.03855 |
| Organelle Organization (GO:0006996) | 4.576659 | 0.03909 |
| Regulation of Macroautophagy (GO:0016241) | 6.603774 | 0.03923 |
| Negative Regulation of DNA-templated Transcription (GO:0045892) | 3.976143 | 0.03928 |
| Canonical Wnt Signaling Pathway (GO:0060070) | 7.936508 | 0.03948 |
| Positive Regulation of Interleukin-1 Production (GO:0032732) | 7.936508 | 0.03948 |
| Positive Regulation of Peptidyl-Tyrosine Phosphorylation (GO:0050731) | 7.936508 | 0.03948 |
| Cellular Response to Oxygen-Containing Compound (GO:1901701) | 4.6875 | 0.04 |
| RNA Splicing, via Endonucleolytic Cleavage and Ligation (GO:0000394) | 18.18182 | 0.0408 |
| Regulation of Mast Cell Degranulation (GO:0043304) | 18.18182 | 0.0408 |
| Defense Response to Tumor Cell (GO:0002357) | 18.18182 | 0.0408 |

| | | |
|---|---|---|
| Tricarboxylic Acid Metabolic Process (GO:0072350) | 18.18182 | 0.0408 |
| Intestinal Lipid Absorption (GO:0098856) | 18.18182 | 0.0408 |
| Negative Regulation of Epithelial Cell Apoptotic Process (GO:1904036) | 18.18182 | 0.0408 |
| Negative Regulation of Intracellular Transport (GO:0032387) | 18.18182 | 0.0408 |
| Positive Regulation of MHC Class II Biosynthetic Process (GO:0045348) | 18.18182 | 0.0408 |
| Positive Regulation of Behavior (GO:0048520) | 18.18182 | 0.0408 |
| Positive Regulation of Endoplasmic Reticulum Unfolded Protein Response (GO:1900103) | 18.18182 | 0.0408 |
| Regulation of Cardiac Muscle Cell Apoptotic Process (GO:0010665) | 18.18182 | 0.0408 |
| Cellular Response to Xenobiotic Stimulus (GO:0071466) | 11.53846 | 0.04118 |
| Limb Development (GO:0060173) | 11.53846 | 0.04118 |
| DNA-templated Transcription (GO:0006351) | 5.418719 | 0.04133 |
| Regulation of Autophagy (GO:0010506) | 5.217391 | 0.04348 |
| DNA Synthesis Involved in DNA Repair (GO:0000731) | 11.11111 | 0.04533 |
| Regulation of Reactive Oxygen Species Biosynthetic Process (GO:1903426) | 11.11111 | 0.04533 |
| Positive Regulation of Biosynthetic Process (GO:0009891) | 6.896552 | 0.04547 |
| Regulation of Leukocyte Adhesion to Vascular Endothelial Cell (GO:1904994) | 16.66667 | 0.04802 |
| Axonal Fasciculation (GO:0007413) | 16.66667 | 0.04802 |
| Branching Involved in Blood Vessel Morphogenesis (GO:0001569) | 16.66667 | 0.04802 |
| Cellular Component Maintenance (GO:0043954) | 16.66667 | 0.04802 |
| Cellular Response to Fluid Shear Stress (GO:0071498) | 16.66667 | 0.04802 |
| Endochondral Bone Morphogenesis (GO:0060350) | 16.66667 | 0.04802 |
| Microtubule Anchoring at Microtubule Organizing Center (GO:0072393) | 16.66667 | 0.04802 |
| Muscle Tissue Development (GO:0060537) | 16.66667 | 0.04802 |
| p38MAPK Cascade (GO:0038066) | 16.66667 | 0.04802 |
| Peptidyl-Tyrosine Autophosphorylation (GO:0038083) | 16.66667 | 0.04802 |
| Positive Regulation of T-helper Cell Differentiation (GO:0045624) | 16.66667 | 0.04802 |
| Positive Regulation of Cellular Extravasation (GO:0002693) | 16.66667 | 0.04802 |
| Positive Regulation of Reactive Oxygen Species Biosynthetic Process (GO:1903428) | 16.66667 | 0.04802 |
| Regulation of SNARE Complex Assembly (GO:0035542) | 16.66667 | 0.04802 |
| Regulation of Actin Filament Depolymerization (GO:0030834) | 16.66667 | 0.04802 |
| Positive Regulation of Leukocyte Migration (GO:0002687) | 10.71429 | 0.04968 |
| Endoplasmic Reticulum to Golgi Vesicle-Mediated Transport (GO:0006888) | 6.25 | 0.05 |

### B. Down Regulated Genes

| GO:BP(LSD1i) | Percentage of Genes | P-value |
|---|---|---|
| Regulation of Protein Neddylation (GO:2000434) | 10.526316 | 0.00032 |
| U2-type Prespliceosome Assembly (GO:1903241) | 8.6956522 | 0.00047 |
| RNA Processing (GO:0006396) | 1.3745704 | 0.00068 |
| protein-RNA Complex Assembly (GO:0022618) | 2.173913 | 0.00093 |
| Ribosome Assembly (GO:0042255) | 4.7619048 | 0.00157 |
| Negative Regulation of Protein Binding (GO:0032091) | 4 | 0.00222 |
| Macromolecule Biosynthetic Process (GO:0009059) | 1.5873016 | 0.00229 |
| Spliceosomal Complex Assembly (GO:0000245) | 3.1746032 | 0.0035 |
| Negative Regulation of Binding (GO:0051100) | 3.030303 | 0.00384 |
| Translation (GO:0006412) | 1.2931034 | 0.00407 |
| Regulation of Protein Binding (GO:0043393) | 2.5641026 | 0.00531 |
| Regulation of Respiratory Burst Involved in Inflammatory Response (GO:0060264) | 20 | 0.00698 |
| Phosphatidylglycerol Biosynthetic Process (GO:0006655) | 20 | 0.00698 |
| RNA Splicing (GO:0008380) | 2.173913 | 0.00732 |
| rRNA Metabolic Process (GO:0016072) | 2.0408163 | 0.00827 |
| 7-Methylguanosine Cap Hypermethylation (GO:0036261) | 16.666667 | 0.00837 |
| Negative Regulation of Mitochondrial Membrane Potential (GO:0010917) | 16.666667 | 0.00837 |
| RNA Capping (GO:0036260) | 16.666667 | 0.00837 |
| Mitochondrial Fragmentation Involved in Apoptotic Process (GO:0043653) | 16.666667 | 0.00837 |
| Clathrin-Coated Vesicle Cargo Loading (GO:0035652) | 16.666667 | 0.00837 |
| Clathrin-Coated Vesicle Cargo Loading, AP-3-mediated (GO:0035654) | 16.666667 | 0.00837 |
| Positive Regulation of Respiratory Burst (GO:0060267) | 16.666667 | 0.00837 |
| Cytoplasmic Translation (GO:0002181) | 1.980198 | 0.00876 |
| Positive Regulation of IRE1-mediated Unfolded Protein Response (GO:1903896) | 14.285714 | 0.00976 |
| rRNA Processing (GO:0006364) | 1.8181818 | 0.01032 |
| Negative Regulation of Membrane Potential (GO:0045837) | 12.5 | 0.01115 |
| UDP-N-acetylglucosamine Biosynthetic Process (GO:0006048) | 12.5 | 0.01115 |
| Negative Regulation of Ubiquitin Protein Ligase Activity (GO:1904667) | 12.5 | 0.01115 |
| Regulation of Cell Killing (GO:0031341) | 12.5 | 0.01115 |
| Endosome to Melanosome Transport (GO:0035646) | 12.5 | 0.01115 |
| Endosome to Pigment Granule Transport (GO:0043485) | 12.5 | 0.01115 |

| | | |
|---|---|---|
| Intrinsic Apoptotic Signaling Pathway (GO:0097193) | 1.7391304 | 0.01124 |
| Ribonucleoprotein Complex Biogenesis (GO:0022613) | 1.6666667 | 0.01219 |
| Synaptic Vesicle Budding From Presynaptic Endocytic Zone Membrane (GO:0016185) | 11.111111 | 0.01253 |
| Regulation of Complement-Dependent Cytotoxicity (GO:1903659) | 11.111111 | 0.01253 |
| Cardiolipin Metabolic Process (GO:0032048) | 11.111111 | 0.01253 |
| Regulation of Complement Activation (GO:0030449) | 10 | 0.01392 |
| UDP-N-acetylglucosamine Metabolic Process (GO:0006047) | 10 | 0.01392 |
| Actin Filament Depolymerization (GO:0030042) | 10 | 0.01392 |
| Negative Regulation of Sprouting Angiogenesis (GO:1903671) | 9.0909091 | 0.0153 |
| Amino Sugar Biosynthetic Process (GO:0046349) | 9.0909091 | 0.0153 |
| Negative Regulation of Ubiquitin-Protein Transferase Activity (GO:0051444) | 9.0909091 | 0.0153 |
| Positive Regulation of Endoplasmic Reticulum Unfolded Protein Response (GO:1900103) | 9.0909091 | 0.0153 |
| Regulation of Chromatin Organization (GO:1902275) | 9.0909091 | 0.0153 |
| Negative Regulation of Complement Activation (GO:0045916) | 8.3333333 | 0.01668 |
| Regulation of Cell Cycle Process (GO:0010564) | 1.369863 | 0.01767 |
| Regulation of Humoral Immune Response (GO:0002920) | 7.6923077 | 0.01805 |
| Extrinsic Apoptotic Signaling Pathway in Absence of Ligand (GO:0097192) | 7.1428571 | 0.01943 |
| Regulation of Immune Effector Process (GO:0002697) | 7.1428571 | 0.01943 |
| Negative Regulation of Immune Effector Process (GO:0002698) | 6.6666667 | 0.0208 |
| Regulation of IRE1-mediated Unfolded Protein Response (GO:1903894) | 6.6666667 | 0.0208 |
| Ribosome Biogenesis (GO:0042254) | 1.242236 | 0.02123 |
| Synaptic Vesicle Transport Along Microtubule (GO:0099517) | 6.25 | 0.02217 |
| Golgi to Vacuole Transport (GO:0006896) | 6.25 | 0.02217 |
| Negative Regulation of Myoblast Differentiation (GO:0045662) | 6.25 | 0.02217 |
| Ribosomal Small Subunit Assembly (GO:0000028) | 6.25 | 0.02217 |
| Anterograde Synaptic Vesicle Transport (GO:0048490) | 6.25 | 0.02217 |
| Positive Regulation of Transcription by RNA Polymerase III (GO:0045945) | 6.25 | 0.02217 |
| Regulation of Ubiquitin Protein Ligase Activity (GO:1904666) | 5.8823529 | 0.02354 |
| Release of Cytochrome C From Mitochondria (GO:0001836) | 5.8823529 | 0.02354 |
| Ribosomal Large Subunit Assembly (GO:0000027) | 5.8823529 | 0.02354 |
| RNA Splicing, via Transesterification Reactions With Bulged Adenosine as Nucleophile (GO:0000377) | 1.1695906 | 0.02375 |
| Melanosome Assembly (GO:1903232) | 5.5555556 | 0.02491 |

| | | |
|---|---|---|
| Regulation of Nuclear Division (GO:0051783) | 5.2631579 | 0.02628 |
| Protein Neddylation (GO:0045116) | 5 | 0.02764 |
| Negative Regulation of Phosphate Metabolic Process (GO:0045936) | 5 | 0.02764 |
| Positive Regulation of Release of Cytochrome C From Mitochondria (GO:0090200) | 5 | 0.02764 |
| Endoplasmic Reticulum Calcium Ion Homeostasis (GO:0032469) | 4.7619048 | 0.02901 |
| Regulation of Anoikis (GO:2000209) | 4.5454545 | 0.03037 |
| Diacylglycerol Metabolic Process (GO:0046339) | 4.5454545 | 0.03037 |
| Nucleotide-Sugar Biosynthetic Process (GO:0009226) | 4.3478261 | 0.03173 |
| Platelet Dense Granule Organization (GO:0060155) | 4.3478261 | 0.03173 |
| Monocyte Chemotaxis (GO:0002548) | 4.3478261 | 0.03173 |
| Phosphatidylglycerol Metabolic Process (GO:0046471) | 4.1666667 | 0.03308 |
| mRNA Splicing, via Spliceosome (GO:0000398) | 0.9478673 | 0.03498 |
| Protein Depolymerization (GO:0051261) | 3.8461538 | 0.03579 |
| Cellular Response to Topologically Incorrect Protein (GO:0035967) | 3.8461538 | 0.03579 |
| Secretory Granule Organization (GO:0033363) | 3.8461538 | 0.03579 |
| Maturation of SSU-rRNA Frm Tricistronic rRNA Trnscpt (GO:0000462) | 3.7037037 | 0.03714 |
| Regulation of Transcription by RNA Polymerase III (GO:0006359) | 3.7037037 | 0.03714 |
| Negative Regulation of Proteolysis Involved in Protein Catabolic Process (GO:1903051) | 3.7037037 | 0.03714 |
| Regulation of Protein Modification by Small Protein Conjugation or Removal (GO:1903320) | 3.5714286 | 0.03849 |
| mRNA Processing (GO:0006397) | 0.8849558 | 0.03963 |
| DNA-templated Transcription Elongation (GO:0006354) | 3.4482759 | 0.03984 |
| Transcription Elongation by RNA Polymerase II (GO:0006368) | 3.4482759 | 0.03984 |
| Vesicle Coating (GO:0006901) | 3.3333333 | 0.04119 |
| Clathrin-Dependent Endocytosis (GO:0072583) | 3.3333333 | 0.04119 |
| Intrinsic Apoptotic Signaling Pathway in Response to Endoplasmic Reticulum Stress (GO:0070059) | 3.3333333 | 0.04119 |
| Regulation of Transcription by RNA Polymerase I (GO:0006356) | 3.2258065 | 0.04253 |
| Extrinsic Apoptotic Signaling Pathway via Death Domain Receptors (GO:0008625) | 100 | 0.04387 |
| Apoptotic Mitochondrial Changes (GO:0008637) | 100 | 0.04387 |
| B Cell Receptor Signaling Pathway (GO:0050853) | 100 | 0.04522 |
| Protein Localization to Chromatin (GO:0071168) | 100 | 0.04522 |
| Synaptic Vesicle Recycling (GO:0036465) | 100 | 0.04655 |
| Mitochondrial Electron Transport, NADH to Ubiquinone (GO:0006120) | 100 | 0.04655 |

| | | |
|---|---|---|
| Positive Regulation of Transcription by RNA Polymerase I (GO:0045943) | 100 | 0.04655 |
| NADH Dehydrogenase Complex Assembly (GO:0010257) | 100 | 0.04789 |
| Mitochondrial Respiratory Chain Complex I Assembly (GO:0032981) | 100 | 0.04789 |
| Regulation of Mitochondrial Membrane Potential (GO:0051881) | 100 | 0.04789 |
| Positive Regulation of Immune Effector Process (GO:0002699) | 100 | 0.04923 |

| **GO:BP(HDAC1i)** | **Percentage of Genes** | **P-value** |
|---|---|---|
| Negative Regulation of Organelle Assembly (GO:1902116) | 9.30232558 | 0.0001396 |
| Positive Regulation of Intracellular Signal Transduction (GO:1902533) | 1.99386503 | 0.0001912 |
| protein-RNA Complex Assembly (GO:0022618) | 4.34782609 | 0.0002123 |
| Positive Regulation of Post-Transcriptional Gene Silencing by RNA (GO:1900370) | 40 | 0.0003706 |
| Negative Regulation of Cell Differentiation (GO:0045596) | 3.18181818 | 0.0004276 |
| Negative Regulation of DNA-templated Transcription (GO:0045892) | 1.59045726 | 0.0004478 |
| mRNA Processing (GO:0006397) | 3.09734513 | 0.0005022 |
| Positive Regulation of miRNA-mediated Gene Silencing (GO:2000637) | 33.3333333 | 0.0005537 |
| mRNA Splice Site Recognition (GO:0006376) | 10.3448276 | 0.0007379 |
| Regulation of miRNA-mediated Gene Silencing (GO:0060964) | 28.5714286 | 0.0007721 |
| Regulation of Cell Population Proliferation (GO:0042127) | 1.69050715 | 0.0009095 |
| Negative Regulation of Fat Cell Differentiation (GO:0045599) | 9.375 | 0.0009883 |
| Response to Steroid Hormone (GO:0048545) | 9.375 | 0.0009883 |
| Negative Regulation of Protein-Containing Complex Disassembly (GO:0043242) | 25 | 0.0010253 |
| Regulation of Pinocytosis (GO:0048548) | 25 | 0.0010253 |
| Regulation of DNA-templated Transcription (GO:0006355) | 1.16877045 | 0.0011698 |
| Import Into Nucleus (GO:0051170) | 5.33333333 | 0.0011821 |
| Transforming Growth Factor Beta Receptor Signaling Pathway (GO:0007179) | 5.2631579 | 0.0012419 |
| Protein Import Into Nucleus (GO:0006606) | 5 | 0.0015024 |
| Protein Localization to Condensed Chromosome (GO:1903083) | 20 | 0.0016346 |
| Regulation of DNA-templated Transcription Initiation (GO:2000142) | 20 | 0.0016346 |
| Negative Regulation of Autophagosome Assembly (GO:1902902) | 20 | 0.0016346 |
| Negative Regulation of Transcription by RNA Polymerase II (GO:0000122) | 1.63934426 | 0.0018599 |
| Positive Regulation of Viral Process (GO:0048524) | 7.5 | 0.0018995 |
| mRNA Splicing, via Spliceosome (GO:0000398) | 2.8436019 | 0.0019612 |
| Protein Localization to Kinetochore (GO:0034501) | 18.1818182 | 0.0019898 |

| | | |
|---|---|---|
| RNA Processing (GO:0006396) | 2.40549828 | 0.0021707 |
| Establishment of Protein Localization to Organelle (GO:0072594) | 4.44444444 | 0.0023156 |
| RNA Transport (GO:0050658) | 6.97674419 | 0.0023411 |
| Regulation of Transcription by RNA Polymerase II (GO:0006357) | 1.11111111 | 0.0023625 |
| Negative Regulation of RNA Biosynthetic Process (GO:1902679) | 1.91489362 | 0.0025095 |
| Response to Progesterone (GO:0032570) | 15.3846154 | 0.0027993 |
| Regulation of Intrinsic Apoptotic Signaling Pathway (GO:2001242) | 6.52173913 | 0.0028413 |
| Positive Regulation of miRNA Transcription (GO:1902895) | 6.38297872 | 0.0030216 |
| Cellular Response to Transforming Growth Factor Beta Stimulus (GO:0071560) | 4.08163265 | 0.0031528 |
| Ribosome Biogenesis (GO:0042254) | 3.10559006 | 0.0032013 |
| Cellular Response to Misfolded Protein (GO:0071218) | 13.3333333 | 0.0037381 |
| Blood Vessel Morphogenesis (GO:0048514) | 5.88235294 | 0.0038121 |
| RNA Splicing, via Transesterification Reactions With Bulged Adenosine as Nucleophile (GO:0000377) | 2.92397661 | 0.0041378 |
| Positive Regulation of miRNA Metabolic Process (GO:2000630) | 5.66037736 | 0.0042502 |
| Regulation of Stem Cell Proliferation (GO:0072091) | 12.5 | 0.004255 |
| Regulation of Autophagosome Maturation (GO:1901096) | 12.5 | 0.004255 |
| Metaphase Chromosome Alignment (GO:0051310) | 5.45454546 | 0.0047177 |
| Positive Regulation by Host of Viral Transcription (GO:0043923) | 11.7647059 | 0.004803 |
| Negative Regulation of Anoikis (GO:2000811) | 11.7647059 | 0.004803 |
| Mitotic Metaphase Chromosome Alignment (GO:0007080) | 5.35714286 | 0.0049626 |
| Positive Regulation of DNA-templated Transcription (GO:0045893) | 1.25588697 | 0.005024 |
| Protein Localization to Chromosome, Centromeric Region (GO:0071459) | 11.1111111 | 0.0053818 |
| Transforming Growth Factor Beta Receptor Superfamily Signaling Pathway (GO:0141091) | 3.50877193 | 0.005403 |
| Positive Regulation of Cell Cycle Phase Transition (GO:1901989) | 10.5263158 | 0.0059908 |
| Regulation of Centriole Replication (GO:0046599) | 10.5263158 | 0.0059908 |
| Post-Transcriptional Regulation of Gene Expression (GO:0010608) | 5 | 0.0060183 |
| Ribonucleoprotein Complex Biogenesis (GO:0022613) | 3.33333333 | 0.0064672 |
| Hippo Signaling (GO:0035329) | 10 | 0.0066299 |
| Spliceosomal Complex Assembly (GO:0000245) | 4.76190476 | 0.0068916 |
| G1/S Transition of Mitotic Cell Cycle (GO:0000082) | 4.6875 | 0.0071985 |
| Positive Regulation of RNA Biosynthetic Process (GO:1902680) | 1.62162162 | 0.0073556 |
| Positive Regulation of Transcription by RNA Polymerase II (GO:0045944) | 1.3224822 | 0.0074934 |

| | | |
|---|---|---|
| Regulation of miRNA Transcription (GO:1902893) | 4.54545455 | 0.0078364 |
| Cell Cycle G1/S Phase Transition (GO:0044843) | 4.54545455 | 0.0078364 |
| Protein Localization to Nucleus (GO:0034504) | 3.1496063 | 0.0078744 |
| Positive Regulation of Cholesterol Efflux (GO:0010875) | 9.09090909 | 0.0079962 |
| Regulation of Anoikis (GO:2000209) | 9.09090909 | 0.0079962 |
| Negative Regulation of Apoptotic Process (GO:0043066) | 1.68421053 | 0.0091353 |
| Negative Regulation of Programmed Cell Death (GO:0043069) | 1.83246073 | 0.0094128 |
| Negative Regulation of Extrinsic Apoptotic Signaling Pathway via Death Domain Receptors (GO:1902042) | 8.33333333 | 0.0094777 |
| Response to Ketone (GO:1901654) | 8.33333333 | 0.0094777 |
| Mitochondrion Organization (GO:0007005) | 2.96296296 | 0.0097131 |
| Negative Regulation of Intracellular Signal Transduction (GO:1902532) | 2.02020202 | 0.0101919 |
| Negative Regulation of Macroautophagy (GO:0016242) | 7.69230769 | 0.0110715 |
| Regulation of Fat Cell Differentiation (GO:0045598) | 4 | 0.0111119 |
| Positive Regulation of Protein Ubiquitination (GO:0031398) | 3.94736842 | 0.0115175 |
| Spindle Assembly Checkpoint Signaling (GO:0071173) | 7.40740741 | 0.0119095 |
| Protein Quality Control for Misfolded or Incompletely Synthesized Proteins (GO:0006515) | 7.40740741 | 0.0119095 |
| Mitotic Spindle Assembly Checkpoint Signaling (GO:0007094) | 7.40740741 | 0.0119095 |
| Mitotic Spindle Checkpoint Signaling (GO:0071174) | 7.40740741 | 0.0119095 |
| Regulation of Cell Cycle Process (GO:0010564) | 2.73972603 | 0.0126666 |
| Positive Regulation of Cholesterol Transport (GO:0032376) | 7.14285714 | 0.0127745 |
| Positive Regulation of Viral Genome Replication (GO:0045070) | 7.14285714 | 0.0127745 |
| Positive Regulation of Protein Modification by Small Protein Conjugation or Removal (GO:1903322) | 3.79746835 | 0.0127852 |
| Positive Regulation of Metabolic Process (GO:0009893) | 2.72108844 | 0.0129604 |
| Cholesterol Homeostasis (GO:0042632) | 3.75 | 0.0132248 |
| Regulation of Cold-Induced Thermogenesis (GO:0120161) | 2.7027027 | 0.0132586 |
| Negative Regulation of Mitotic Metaphase/Anaphase Transition (GO:0045841) | 6.89655172 | 0.0136661 |
| Sterol Homeostasis (GO:0055092) | 3.7037037 | 0.013673 |
| Translation (GO:0006412) | 2.15517241 | 0.0144006 |
| Negative Regulation of Response to Endoplasmic Reticulum Stress (GO:1903573) | 6.66666667 | 0.014584 |
| Positive Regulation of Cellular Process (GO:0048522) | 1.44694534 | 0.0147081 |
| Intracellular Protein Transport (GO:0006886) | 1.8404908 | 0.0155326 |

| | | |
|---|---|---|
| Regulation of Extrinsic Apoptotic Signaling Pathway via Death Domain Receptors (GO:1902041) | 6.25 | 0.016497 |
| miRNA Processing (GO:0035196) | 6.25 | 0.016497 |
| Mitochondrial Membrane Organization (GO:0007006) | 6.25 | 0.016497 |
| Regulation of Cholesterol Efflux (GO:0010874) | 6.25 | 0.016497 |
| Negative Regulation of Cellular Process (GO:0048523) | 1.50659134 | 0.0169186 |
| Vasculogenesis (GO:0001570) | 6.06060606 | 0.0174915 |
| Negative Regulation of Endocytosis (GO:0045806) | 5.88235294 | 0.0185109 |
| Integrated Stress Response Signaling (GO:0140467) | 5.88235294 | 0.0185109 |
| Regulation of Double-Strand Break Repair (GO:2000779) | 3.2967033 | 0.0186291 |
| Positive Regulation of Mitotic Nuclear Division (GO:0045840) | 5.71428571 | 0.0195548 |
| Regulation of MAPK Cascade (GO:0043408) | 1.92307692 | 0.0223684 |
| Nuclear Transport (GO:0051169) | 5.2631579 | 0.0228303 |
| Cellular Response to Chemical Stress (GO:0062197) | 3 | 0.0238342 |
| Positive Regulation of Intrinsic Apoptotic Signaling Pathway (GO:2001244) | 5.12820513 | 0.0239689 |
| Cytoplasmic Translation (GO:0002181) | 2.97029703 | 0.0244563 |
| mRNA Metabolic Process (GO:0016071) | 2.97029703 | 0.0244563 |
| Negative Regulation of Translation (GO:0017148) | 2.94117647 | 0.0250871 |
| Positive Regulation of Protein Localization (GO:1903829) | 2.94117647 | 0.0250871 |
| Acylglycerol Metabolic Process (GO:0006639) | 4.76190476 | 0.0275209 |
| Ribosome Assembly (GO:0042255) | 4.76190476 | 0.0275209 |
| Regulation of Apoptotic Process (GO:0042981) | 1.27840909 | 0.0298256 |
| Water-Soluble Vitamin Metabolic Process (GO:0006767) | 4.54545455 | 0.0299991 |
| Regulation of Autophagosome Assembly (GO:2000785) | 4.54545455 | 0.0299991 |
| Positive Regulation of Cell Population Proliferation (GO:0008284) | 1.44628099 | 0.0300897 |
| Reg of Epithelial to Mesenchymal Transition Inv in Endocardial Cushion Formation (GO:1905005) | 20 | 0.0303769 |
| Regulation of Fat Cell Proliferation (GO:0070344) | 20 | 0.0303769 |
| Negative Regulation of Keratinocyte Differentiation (GO:0045617) | 20 | 0.0303769 |
| Negative Regulation of Lipid Transport (GO:0032369) | 20 | 0.0303769 |
| Regulation of Stress Granule Assembly (GO:0062028) | 20 | 0.0303769 |
| Peptidyl-Proline Modification (GO:0018208) | 20 | 0.0303769 |
| Phosphatidylglycerol Biosynthetic Process (GO:0006655) | 20 | 0.0303769 |
| snRNA Modification (GO:0040031) | 20 | 0.0303769 |

| | | |
|---|---|---|
| Embryonic Brain Development (GO:1990403) | 20 | 0.0303769 |
| Positive Regulation of Gap Junction Assembly (GO:1903598) | 20 | 0.0303769 |
| Tricuspid Valve Development (GO:0003175) | 20 | 0.0303769 |
| Tricuspid Valve Morphogenesis (GO:0003186) | 20 | 0.0303769 |
| mRNA 3'-Splice Site Recognition (GO:0000389) | 20 | 0.0303769 |
| Negative Regulation of T-helper 17 Cell Lineage Commitment (GO:2000329) | 20 | 0.0303769 |
| Negative Regulation of Cell Fate Commitment (GO:0010454) | 20 | 0.0303769 |
| Gene Expression (GO:0010467) | 1.57480315 | 0.0304672 |
| Regulation of Nervous System Development (GO:0051960) | 4.44444444 | 0.0312704 |
| Regulation of Protein Ubiquitination (GO:0031396) | 2.65486726 | 0.0326008 |
| Mitotic Cell Cycle Phase Transition (GO:0044772) | 2.65486726 | 0.0326008 |
| Negative Regulation of Cold-Induced Thermogenesis (GO:0120163) | 4.25531915 | 0.0338755 |
| Ribosomal Large Subunit Biogenesis (GO:0042273) | 4.25531915 | 0.0338755 |
| Positive Regulation of Pattern Recognition Receptor Signaling Pathway (GO:0062208) | 4.25531915 | 0.0338755 |
| Regulation of Translation (GO:0006417) | 1.99004975 | 0.0356824 |
| Positive Regulation of Protein Metabolic Process (GO:0051247) | 1.98019802 | 0.0362362 |
| Regulation of Interleukin-1-Mediated Signaling Pathway (GO:2000659) | 16.6666667 | 0.0363416 |
| RIG-I Signaling Pathway (GO:0039529) | 16.6666667 | 0.0363416 |
| Negative Regulation of Response to Food (GO:0032096) | 16.6666667 | 0.0363416 |
| Regulation of Mitotic Cytokinetic Process (GO:1903436) | 16.6666667 | 0.0363416 |
| Regulation of Vascular Associated Smooth Muscle Cell Apoptotic Process (GO:1905459) | 16.6666667 | 0.0363416 |
| Centriole Assembly (GO:0098534) | 16.6666667 | 0.0363416 |
| Ribosomal Large Subunit Export From Nucleus (GO:0000055) | 16.6666667 | 0.0363416 |
| Ribosomal Small Subunit Export From Nucleus (GO:0000056) | 16.6666667 | 0.0363416 |
| Positive Regulation of Endothelial Cell-Matrix Adhesion via Fibronectin (GO:1904906) | 16.6666667 | 0.0363416 |
| Endothelin Receptor Signaling Pathway (GO:0086100) | 16.6666667 | 0.0363416 |
| Positive Regulation of Lipoprotein Lipase Activity (GO:0051006) | 16.6666667 | 0.0363416 |
| Positive Regulation of Mitotic Cytokinetic Process (GO:1903438) | 16.6666667 | 0.0363416 |
| Positive Regulation of Protein Localization to Centrosome (GO:1904781) | 16.6666667 | 0.0363416 |
| Positive Regulation of Sterol Transport (GO:0032373) | 16.6666667 | 0.0363416 |
| Long-Term Synaptic Depression (GO:0060292) | 16.6666667 | 0.0363416 |
| Regulation of T Cell Tolerance Induction (GO:0002664) | 16.6666667 | 0.0363416 |

| | | |
|---|---|---|
| Regulation of Cell Adhesion Molecule Production (GO:0060353) | 16.6666667 | 0.0363416 |
| Positive Regulation of Epithelial Cell Proliferation (GO:0050679) | 2.54237288 | 0.0363621 |
| Negative Regulation of Small GTPase Mediated Signal Transduction (GO:0051058) | 4.08163265 | 0.0365623 |
| Regulation of Wound Healing (GO:0061041) | 4.08163265 | 0.0365623 |
| Positive Regulation of Canonical NF-kappaB Signal Transduction (GO:0043123) | 1.96078431 | 0.0373587 |
| Mitotic Sister Chromatid Segregation (GO:0000070) | 2.5 | 0.0379265 |
| Nuclear Export (GO:0051168) | 4 | 0.0379356 |
| Nucleocytoplasmic Transport (GO:0006913) | 4 | 0.0379356 |
| DNA Metabolic Process (GO:0006259) | 1.65562914 | 0.0389311 |
| Regulation of Mitotic Metaphase/Anaphase Transition (GO:0030071) | 3.92156863 | 0.0393284 |
| Regulation of Ubiquitin-Dependent Protein Catabolic Process (GO:2000058) | 3.92156863 | 0.0393284 |
| Positive Regulation of Reactive Oxygen Species Metabolic Process (GO:2000379) | 3.84615385 | 0.0407405 |
| Intrinsic Apoptotic Signaling Pathway in Response to DNA Damage (GO:0008630) | 3.84615385 | 0.0407405 |
| Regulation of Cell Development (GO:0060284) | 3.84615385 | 0.0407405 |
| Cellular Response to Ionizing Radiation (GO:0071479) | 3.77358491 | 0.0421714 |
| Cytoplasmic Microtubule Organization (GO:0031122) | 3.77358491 | 0.0421714 |
| Negative Regulation of Epidermal Cell Differentiation (GO:0045605) | 14.2857143 | 0.0422698 |
| DNA Geometric Change (GO:0032392) | 14.2857143 | 0.0422698 |
| Regulation of Endothelial Cell-Matrix Adhesion via Fibronectin (GO:1904904) | 14.2857143 | 0.0422698 |
| Negative Regulation of Response to Type II Interferon (GO:0060331) | 14.2857143 | 0.0422698 |
| Negative Regulation of Type II Interferon-Mediated Signaling Pathway (GO:0060336) | 14.2857143 | 0.0422698 |
| Positive Regulation of Cardiac Epithelial to Mesenchymal Transition (GO:0062043) | 14.2857143 | 0.0422698 |
| tRNA Aminoacylation for Mitochondrial Protein Translation (GO:0070127) | 14.2857143 | 0.0422698 |
| Positive Regulation of Leukocyte Differentiation (GO:1902107) | 14.2857143 | 0.0422698 |
| Positive Regulation of Triglyceride Lipase Activity (GO:0061365) | 14.2857143 | 0.0422698 |
| Reg of Cardiac Muscle Cell Action Potential Inv in Reg of Contraction (GO:0098909) | 14.2857143 | 0.0422698 |
| Regulation of Signal Transduction by P53 Class Mediator (GO:1901796) | 3.7037037 | 0.0436211 |
| DNA Damage Response (GO:0006974) | 1.42517815 | 0.0458472 |
| Regulation of Proteasomal Protein Catabolic Process (GO:0061136) | 3.57142857 | 0.0465753 |
| Regulation of Epithelial Cell Apoptotic Process (GO:1904035) | 12.5 | 0.0481619 |
| Regulation of Gap Junction Assembly (GO:1903596) | 12.5 | 0.0481619 |
| Negative Regulation of Nitric Oxide Biosynthetic Process (GO:0045019) | 12.5 | 0.0481619 |
| Negative Regulation of Nitric Oxide Metabolic Process (GO:1904406) | 12.5 | 0.0481619 |

| | | |
|---|---|---|
| Regulation of Response to Wounding (GO:1903034) | 12.5 | 0.0481619 |
| Cellular Response to Hydroxyurea (GO:0072711) | 12.5 | 0.0481619 |
| Positive Regulation of Megakaryocyte Differentiation (GO:0045654) | 12.5 | 0.0481619 |
| Regulation of CD4-positive, Alpha-Beta T Cell Proliferation (GO:2000561) | 12.5 | 0.0481619 |
| Negative Regulation of DNA-templated DNA Replication (GO:2000104) | 12.5 | 0.0481619 |
| Regulation of Myoblast Differentiation (GO:0045661) | 3.44827586 | 0.0496009 |

| **GO:BP(LSD1i plus HDAC1i)** | **Percentage of Genes** | **P-value** |
|---|---|---|
| RNA Processing (GO:0006396) | 7.560137 | 1.15E-09 |
| mRNA Processing (GO:0006397) | 9.734513 | 1.00E-07 |
| Regulation of Translation (GO:0006417) | 10.94527 | 1.10E-07 |
| mRNA Splicing, via Spliceosome (GO:0000398) | 7.582938 | 2.14E-07 |
| Regulation of DNA-templated Transcription (GO:0006355) | 2.898551 | 9.39E-07 |
| RNA Splicing, via Transesterification Reactions With Bulged Adenosine as Nucleophile (GO:0000377) | 7.602339 | 2.89E-06 |
| Chromatin Organization (GO:0006325) | 5.572755 | 3.41E-06 |
| Regulation of Transcription by RNA Polymerase II (GO:0006357) | 2.755556 | 4.98E-06 |
| protein-RNA Complex Assembly (GO:0022618) | 7.971014 | 1.07E-05 |
| Negative Regulation of DNA-templated Transcription (GO:0045892) | 3.379722 | 1.85E-05 |
| Regulation of mRNA Splicing, via Spliceosome (GO:0048024) | 9.278351 | 2.04E-05 |
| Regulation of Stem Cell Population Maintenance (GO:2000036) | 12.5 | 2.48E-05 |
| mRNA Metabolic Process (GO:0016071) | 8.910891 | 2.83E-05 |
| Positive Regulation of DNA-templated Transcription (GO:0045893) | 3.061224 | 4.06E-05 |
| Positive Regulation of Organelle Assembly (GO:1902117) | 10.76923 | 6.61E-05 |
| RNA Splicing (GO:0008380) | 8.695652 | 9.37E-05 |
| Regulation of Mitotic Metaphase/Anaphase Transition (GO:0030071) | 11.76471 | 0.00013467 |
| Regulation of G0 to G1 Transition (GO:0070316) | 15.15152 | 0.00014616 |
| Positive Regulation of RNA Biosynthetic Process (GO:1902680) | 3.783784 | 0.00017444 |
| Negative Regulation of Transcription by RNA Polymerase II (GO:0000122) | 3.415301 | 0.00021085 |
| Chromatin Remodeling (GO:0006338) | 4.961832 | 0.0002486 |
| N-glycan Processing (GO:0006491) | 19.04762 | 0.00027849 |
| Positive Regulation of Translation (GO:0045727) | 7.407407 | 0.00028618 |
| Negative Regulation of Cell-Substrate Adhesion (GO:0010812) | 12.82051 | 0.00032855 |
| Positive Regulation of Cell Cycle G2/M Phase Transition (GO:1902751) | 17.3913 | 0.00040206 |

| | | |
|---|---|---|
| CRD-mediated mRNA Stabilization (GO:0070934) | 30 | 0.00041208 |
| Positive Regulation of Centriole Replication (GO:0046601) | 30 | 0.00041208 |
| RNA Metabolic Process (GO:0016070) | 9.52381 | 0.00043474 |
| Spliceosomal Complex Assembly (GO:0000245) | 9.52381 | 0.00043474 |
| Positive Regulation of Mitotic Cell Cycle Phase Transition (GO:1901992) | 9.375 | 0.00047354 |
| RNA Transport (GO:0050658) | 11.62791 | 0.00052214 |
| Negative Regulation of Platelet-Derived Growth Factor Receptor Signaling Pathway (GO:0010642) | 27.27273 | 0.0005601 |
| Regulation of Chromatin Organization (GO:1902275) | 27.27273 | 0.0005601 |
| Regulation of Telomere Maintenance via Telomerase (GO:0032210) | 11.36364 | 0.00058166 |
| Regulation of Nucleotide-Excision Repair (GO:2000819) | 15.38462 | 0.00065433 |
| Positive Regulation of Double-Strand Break Repair (GO:2000781) | 7.446809 | 0.00066244 |
| Regulation of mRNA Catabolic Process (GO:0061013) | 10.6383 | 0.00079097 |
| Protein Localization to Nucleus (GO:0034504) | 6.299213 | 0.00084568 |
| Nuclear RNA Surveillance (GO:0071027) | 23.07692 | 0.00094868 |
| Cell-Substrate Junction Assembly (GO:0007044) | 13.7931 | 0.00100213 |
| Negative Regulation of Translation (GO:0017148) | 6.862745 | 0.00107576 |
| Import Into Nucleus (GO:0051170) | 8 | 0.00110296 |
| Regulation of Metaphase/Anaphase Transition of Cell Cycle (GO:1902099) | 12.5 | 0.00146274 |
| Negative Regulation of mRNA Splicing, via Spliceosome (GO:0048025) | 20 | 0.00147488 |
| Protein Import Into Nucleus (GO:0006606) | 7.5 | 0.00154307 |
| Positive Regulation of Transcription by RNA Polymerase II (GO:0045944) | 2.848423 | 0.00155087 |
| Regulation of Alternative mRNA Splicing, via Spliceosome (GO:0000381) | 9.090909 | 0.00162143 |
| Negative Regulation of Gene Expression (GO:0010629) | 3.846154 | 0.00178255 |
| Negative Regulation of mRNA Processing (GO:0050686) | 18.75 | 0.00179445 |
| Intracellular Protein Transport (GO:0006886) | 3.98773 | 0.00187659 |
| Regulation of Protein Ubiquitination (GO:0031396) | 6.19469 | 0.0019473 |
| Regulation of Myoblast Differentiation (GO:0045661) | 8.62069 | 0.00205646 |
| Regulation of Cell Cycle Process (GO:0010564) | 5.479452 | 0.00206468 |
| Regulation of Sodium Ion Transmembrane Transporter Activity (GO:2000649) | 17.64706 | 0.00215406 |
| Post-Transcriptional Regulation of Gene Expression (GO:0010608) | 8.333333 | 0.00239018 |
| Negative Regulation of Cell Differentiation (GO:0045596) | 4.545455 | 0.00242039 |
| Regulation of Mitotic Cell Cycle Phase Transition (GO:1901990) | 6.818182 | 0.00251011 |

| | | |
|---|---|---|
| Regulation of mRNA Processing (GO:0050684) | 10.81081 | 0.00252789 |
| Regulation of G2/M Transition of Mitotic Cell Cycle (GO:0010389) | 10.81081 | 0.00252789 |
| Negative Regulation of DNA-templated Transcription, Elongation (GO:0032785) | 16.66667 | 0.00255532 |
| Establishment of Protein Localization to Organelle (GO:0072594) | 6.666667 | 0.00281053 |
| Negative Regulation of RNA Biosynthetic Process (GO:1902679) | 3.404255 | 0.00296896 |
| Regulation of Double-Strand Break Repair (GO:2000779) | 6.593407 | 0.00297045 |
| Positive Regulation of Cellular Component Biogenesis (GO:0044089) | 6.451613 | 0.00331059 |
| Nucleic Acid Catabolic Process (GO:0141188) | 10 | 0.00337281 |
| Response to UV-C (GO:0010225) | 33.33333 | 0.00346909 |
| Regulation of Nuclear-Transcribed mRNA Catabolic Process, Deadenylation-Dependent Decay (GO:1900151) | 15 | 0.0034889 |
| Iron Ion Transport (GO:0006826) | 15 | 0.0034889 |
| Negative Regulation of RNA Splicing (GO:0033119) | 15 | 0.0034889 |
| Nucleosome Disassembly (GO:0006337) | 15 | 0.0034889 |
| Positive Regulation of G2/M Transition of Mitotic Cell Cycle (GO:0010971) | 15 | 0.0034889 |
| Positive Regulation of Myoblast Differentiation (GO:0045663) | 9.756098 | 0.00369236 |
| Ribosome Biogenesis (GO:0042254) | 4.968944 | 0.00377012 |
| RNA Export From Nucleus (GO:0006405) | 7.462687 | 0.00387065 |
| RNA Biosynthetic Process (GO:0032774) | 5.46875 | 0.00391679 |
| Regulation of Cell Cycle (GO:0051726) | 3.773585 | 0.00431151 |
| Regulation of Cell Differentiation (GO:0045595) | 4.477612 | 0.00433511 |
| Positive Regulation of Protein Metabolic Process (GO:0051247) | 4.455446 | 0.00447711 |
| Focal Adhesion Assembly (GO:0048041) | 13.63636 | 0.00460624 |
| Positive Regulation of Telomere Maintenance via Telomerase (GO:0032212) | 13.63636 | 0.00460624 |
| protein-DNA Complex Disassembly (GO:0032986) | 13.63636 | 0.00460624 |
| U2-type Prespliceosome Assembly (GO:1903241) | 13.04348 | 0.00523687 |
| Positive Regulation of Cell-Substrate Junction Organization (GO:0150117) | 13.04348 | 0.00523687 |
| Regulation of RNA Splicing (GO:0043484) | 5.769231 | 0.00572261 |
| Protein Phosphorylation (GO:0006468) | 3.485255 | 0.00585917 |
| Positive Regulation of Cytoskeleton Organization (GO:0051495) | 6.756757 | 0.00591394 |
| RNA Splicing, via Transesterification Reactions (GO:0000375) | 12.5 | 0.00591692 |
| Negative Regulation of Extrinsic Apoptotic Signaling Pathway via Death Domain Receptors (GO:1902042) | 12.5 | 0.00591692 |
| Transferrin Transport (GO:0033572) | 25 | 0.00634367 |
| Regulation of RNA Biosynthetic Process (GO:2001141) | 3.225806 | 0.00643822 |

| | | |
|---|---|---|
| Regulation of Muscle Cell Differentiation (GO:0051147) | 12 | 0.00664734 |
| Positive Regulation of Telomere Maintenance via Telomere Lengthening (GO:1904358) | 12 | 0.00664734 |
| Nucleosome Organization (GO:0034728) | 6.493506 | 0.00698651 |
| Regulation of Stem Cell Differentiation (GO:2000736) | 8.163265 | 0.00701222 |
| Phosphorylation (GO:0016310) | 3.691275 | 0.00721685 |
| Macromolecule Methylation (GO:0043414) | 11.53846 | 0.00742904 |
| rRNA Processing (GO:0006364) | 5.454545 | 0.00748061 |
| Nuclear Export (GO:0051168) | 8 | 0.00753051 |
| RNA Catabolic Process (GO:0006401) | 6.329114 | 0.00777277 |
| Regulation of Cell Cycle G1/S Phase Transition (GO:1902806) | 6.329114 | 0.00777277 |
| Regulation of G1/S Transition of Mitotic Cell Cycle (GO:2000045) | 5.405405 | 0.00780762 |
| Negative Regulation of Protein Metabolic Process (GO:0051248) | 4.794521 | 0.00794601 |
| Sex-Chromosome Dosage Compensation (GO:0007549) | 22.22222 | 0.00807278 |
| Negative Regulation of PERK-mediated Unfolded Protein Response (GO:1903898) | 22.22222 | 0.00807278 |
| Positive Regulation of Keratinocyte Migration (GO:0051549) | 22.22222 | 0.00807278 |
| rRNA 3'-End Processing (GO:0031125) | 22.22222 | 0.00807278 |
| Transcription Initiation-Coupled Chromatin Remodeling (GO:0045815) | 7.843137 | 0.0080734 |
| Regulation of mRNA Stability (GO:0043488) | 6.25 | 0.00818808 |
| Negative Regulation of Cell Migration (GO:0030336) | 4.347826 | 0.00828337 |
| Intrinsic Apoptotic Signaling Pathway in Response to DNA Damage (GO:0008630) | 7.692308 | 0.00864134 |
| Negative Regulation of Protein Ubiquitination (GO:0031397) | 7.692308 | 0.00864134 |
| Negative Regulation of Telomere Maintenance via Telomere Lengthening (GO:1904357) | 10.71429 | 0.00914942 |
| Regulation of Signal Transduction by P53 Class Mediator (GO:1901796) | 7.407407 | 0.00985415 |
| Regulation of Telomere Capping (GO:1904353) | 20 | 0.00998797 |
| Establishment of Endothelial Intestinal Barrier (GO:0090557) | 20 | 0.00998797 |
| Nuclear mRNA Surveillance (GO:0071028) | 20 | 0.00998797 |
| Positive Regulation of Attachment of Spindle Microtubules to Kinetochore (GO:0051987) | 20 | 0.00998797 |
| Regulation of Androgen Receptor Signaling Pathway (GO:0060765) | 10.34483 | 0.0100895 |
| Negative Regulation of Transforming Growth Factor Beta Receptor Signaling Pathway (GO:0030512) | 5.882353 | 0.01049699 |
| Regulation of Peptide Hormone Secretion (GO:0090276) | 7.272727 | 0.01049985 |
| Negative Regulation of Cell-Matrix Adhesion (GO:0001953) | 10 | 0.01108364 |

| | | |
|---|---|---|
| Regulation of Transforming Growth Factor Beta Receptor Signaling Pathway (GO:0017015) | 5 | 0.01122867 |
| Ribonucleoprotein Complex Biogenesis (GO:0022613) | 5 | 0.01122867 |
| mRNA Stabilization (GO:0048255) | 7.017544 | 0.01187187 |
| N-terminal Protein Amino Acid Acetylation (GO:0006474) | 18.18182 | 0.01208306 |
| Regulation of Keratinocyte Migration (GO:0051547) | 18.18182 | 0.01208306 |
| sno(s)RNA 3'-End Processing (GO:0031126) | 18.18182 | 0.01208306 |
| Dosage Compensation by Inactivation of X Chromosome (GO:0009048) | 18.18182 | 0.01208306 |
| - Reg of Nuclear-Transcribed mRNA Cat Proc, Deadenylation-Dep Decay (GO:1900152) | 18.18182 | 0.01208306 |
| Positive Regulation of Metaphase/Anaphase Transition of Cell Cycle (GO:1902101) | 18.18182 | 0.01208306 |
| Negative Regulation of Intracellular Steroid Hormone Receptor Signaling Pathway (GO:0033144) | 9.677419 | 0.01213236 |
| Regulation of Cellular Component Organization (GO:0051128) | 4.402516 | 0.01234404 |
| Negative Regulation of Protein Modification by Small Protein Conjugation or Removal (GO:1903321) | 6.896552 | 0.01259894 |
| Nuclear-Transcribed mRNA Catabolic Process (GO:0000956) | 5.617978 | 0.01263803 |
| Regulation of Extrinsic Apoptotic Signaling Pathway via Death Domain Receptors (GO:1902041) | 9.375 | 0.01323611 |
| mRNA Transport (GO:0051028) | 6.779661 | 0.01335386 |
| Regulation of Epithelial to Mesenchymal Transition (GO:0010717) | 5.494505 | 0.0138118 |
| Proteasomal Protein Catabolic Process (GO:0010498) | 3.225806 | 0.0140343 |
| Negative Regulation of Multicellular Organismal Process (GO:0051241) | 3.703704 | 0.01412399 |
| Negative Regulation of Substrate Adhesion-Dependent Cell Spreading (GO:1900025) | 16.66667 | 0.01435203 |
| Regulation of PERK-mediated Unfolded Protein Response (GO:1903897) | 16.66667 | 0.01435203 |
| Positive Regulation of Canonical NF-kappaB Signal Transduction (GO:0043123) | 3.921569 | 0.01476687 |
| Regulation of Protein Metabolic Process (GO:0051246) | 5.376344 | 0.01505681 |
| Positive Regulation of DNA Repair (GO:0045739) | 5.376344 | 0.01505681 |
| Protein Acylation (GO:0043543) | 8.823529 | 0.01561023 |
| Regulation of DNA Repair (GO:0006282) | 4.615385 | 0.01614186 |
| Gene Expression (GO:0010467) | 3.149606 | 0.01662675 |
| Regulation of Hematopoietic Stem Cell Differentiation (GO:1902036) | 15.38462 | 0.01678901 |
| Formation of Cytoplasmic Translation Initiation Complex (GO:0001732) | 15.38462 | 0.01678901 |
| Microvillus Organization (GO:0032528) | 15.38462 | 0.01678901 |
| Negative Regulation of Stem Cell Differentiation (GO:2000737) | 15.38462 | 0.01678901 |
| Positive Regulation of Mitotic Metaphase/Anaphase Transition (GO:0045842) | 15.38462 | 0.01678901 |

| | | |
|---|---|---|
| Positive Regulation of Mitotic Nuclear Division (GO:0045840) | 8.571429 | 0.01688118 |
| Negative Regulation of Epithelial to Mesenchymal Transition (GO:0010719) | 8.333333 | 0.01820836 |
| Regulation of Cell Migration (GO:0030334) | 2.922756 | 0.01831638 |
| rRNA Metabolic Process (GO:0016072) | 5.102041 | 0.01849205 |
| Regulation of Protein Secretion (GO:0050708) | 5.050505 | 0.01923601 |
| Extrinsic Apoptotic Signaling Pathway in Absence of Ligand (GO:0097192) | 14.28571 | 0.01938826 |
| Microvillus Assembly (GO:0030033) | 14.28571 | 0.01938826 |
| Negative Regulation of Androgen Receptor Signaling Pathway (GO:0060766) | 14.28571 | 0.01938826 |
| Negative Regulation of Transcription Elongation by RNA Polymerase II (GO:0034244) | 14.28571 | 0.01938826 |
| Negative Regulation of Binding (GO:0051100) | 6.060606 | 0.01944519 |
| Positive Regulation of Gene Expression (GO:0010628) | 2.981651 | 0.01950009 |
| Spliceosomal snRNP Assembly (GO:0000387) | 8.108108 | 0.01959194 |
| Negative Regulation of Endothelial Cell Migration (GO:0010596) | 8.108108 | 0.01959194 |
| Maturation of SSU-rRNA (GO:0030490) | 7.894737 | 0.02103201 |
| Nuclear Transport (GO:0051169) | 7.894737 | 0.02103201 |
| Ras Protein Signal Transduction (GO:0007265) | 5.882353 | 0.02145333 |
| Extrinsic Apoptotic Signaling Pathway (GO:0097191) | 5.882353 | 0.02145333 |
| Membraneless Organelle Assembly (GO:0140694) | 4.316547 | 0.02168495 |
| Regulation of Peptidase Activity (GO:0052547) | 13.33333 | 0.02214417 |
| Clathrin Coat Assembly (GO:0048268) | 13.33333 | 0.02214417 |
| Positive Regulation of Cytoplasmic Translation (GO:2000767) | 13.33333 | 0.02214417 |
| Positive Regulation of Mitotic Sister Chromatid Separation (GO:1901970) | 13.33333 | 0.02214417 |
| Regulation of Translational Initiation (GO:0006446) | 5.797101 | 0.02250322 |
| Positive Regulation of Lymphocyte Differentiation (GO:0045621) | 7.692308 | 0.02252863 |
| Peptidyl-Serine Phosphorylation (GO:0018105) | 5.714286 | 0.02358392 |
| Positive Regulation of Stem Cell Population Maintenance (GO:1902459) | 7.5 | 0.0240818 |
| Regulation of Stem Cell Proliferation (GO:0072091) | 12.5 | 0.02505128 |
| Endoplasmic Reticulum to Cytosol Transport (GO:1903513) | 12.5 | 0.02505128 |
| Regulation of Hippo Signaling (GO:0035330) | 7.317073 | 0.0256915 |
| RNA Stabilization (GO:0043489) | 7.317073 | 0.0256915 |
| Regulation of Mitotic Spindle Organization (GO:0060236) | 7.317073 | 0.0256915 |
| Positive Regulation of DNA Biosynthetic Process (GO:2000573) | 7.317073 | 0.0256915 |
| Regulation of Canonical NF-kappaB Signal Transduction (GO:0043122) | 3.333333 | 0.02591194 |

| | | |
|---|---|---|
| Positive Regulation of Intracellular Signal Transduction (GO:1902533) | 2.607362 | 0.02690279 |
| Heterochromatin Formation (GO:0031507) | 5.479452 | 0.02701268 |
| Regulation of DNA Replication (GO:0006275) | 5.479452 | 0.02701268 |
| Ribosome Assembly (GO:0042255) | 7.142857 | 0.02735763 |
| Positive Regulation of Double-Strand Break Repair via Homologous Recombination (GO:1905168) | 7.142857 | 0.02735763 |
| Regulation of mRNA Metabolic Process (GO:1903311) | 11.76471 | 0.02810425 |
| Release of Cytochrome C From Mitochondria (GO:0001836) | 11.76471 | 0.02810425 |
| Negative Regulation of mRNA Catabolic Process (GO:1902373) | 6.976744 | 0.02908007 |
| Positive Regulation of Gene Expression, Epigenetic (GO:0141137) | 6.976744 | 0.02908007 |
| Regulation of Gene Expression (GO:0010468) | 2.30701 | 0.02921849 |
| Peptidyl-Serine Modification (GO:0018209) | 5.333333 | 0.02945555 |
| Protein-Containing Complex Assembly (GO:0065003) | 3.095975 | 0.03034861 |
| Embryonic Organ Morphogenesis (GO:0048562) | 6.818182 | 0.03085865 |
| NLS-bearing Protein Import Into Nucleus (GO:0006607) | 11.11111 | 0.03129787 |
| Regulation of Microtubule Nucleation (GO:0010968) | 11.11111 | 0.03129787 |
| Retrograde Protein Transport, ER to Cytosol (GO:0030970) | 11.11111 | 0.03129787 |
| Negative Regulation of Endoplasmic Reticulum Unfolded Protein Response (GO:1900102) | 11.11111 | 0.03129787 |
| Negative Regulation of Nucleocytoplasmic Transport (GO:0046823) | 11.11111 | 0.03129787 |
| Negative Regulation of Translational Initiation (GO:0045947) | 11.11111 | 0.03129787 |
| Neuron Maturation (GO:0042551) | 11.11111 | 0.03129787 |
| Positive Regulation of Focal Adhesion Assembly (GO:0051894) | 11.11111 | 0.03129787 |
| Positive Regulation of Muscle Cell Differentiation (GO:0051149) | 11.11111 | 0.03129787 |
| Positive Regulation of Stem Cell Proliferation (GO:2000648) | 11.11111 | 0.03129787 |
| Positive Regulation of Wnt Signaling Pathway (GO:0030177) | 4.385965 | 0.03281224 |
| Response to Endoplasmic Reticulum Stress (GO:0034976) | 3.571429 | 0.03400601 |
| Regulation of Cyclin-Dependent Protein Serine/Threonine Kinase Activity (GO:0000079) | 6.521739 | 0.03458338 |
| Homotypic Cell-Cell Adhesion (GO:0034109) | 6.521739 | 0.03458338 |
| Negative Regulation of Type I Interferon-Mediated Signaling Pathway (GO:0060339) | 10.52632 | 0.03462706 |
| Regulation of Centriole Replication (GO:0046599) | 10.52632 | 0.03462706 |
| Regulation of Transcription Elongation by RNA Polymerase II (GO:0034243) | 5.063291 | 0.03472231 |
| Regulation of Double-Strand Break Repair via Homologous Recombination (GO:0010569) | 5 | 0.03611885 |

| | | |
|---|---|---|
| Protein Localization to Membrane (GO:0072657) | 3.517588 | 0.03645594 |
| Positive Regulation of mRNA Catabolic Process (GO:0061014) | 6.382979 | 0.03652901 |
| Regulation of T Cell Differentiation (GO:0045580) | 6.382979 | 0.03652901 |
| Regulation of Cytokinesis (GO:0032465) | 4.938272 | 0.03754744 |
| Regulation of Catabolic Process (GO:0009894) | 4.938272 | 0.03754744 |
| Alternative mRNA Splicing, via Spliceosome (GO:0000380) | 10 | 0.03808686 |
| Regulation of Sodium Ion Transmembrane Transport (GO:1902305) | 10 | 0.03808686 |
| Regulation of Spindle Organization (GO:0090224) | 10 | 0.03808686 |
| Positive Regulation of Developmental Process (GO:0051094) | 3.265306 | 0.03834121 |
| Positive Regulation of Cell Cycle Process (GO:0090068) | 4.201681 | 0.03839063 |
| Proteasome-Mediated Ubiquitin-Dependent Protein Catabolic Process (GO:0043161) | 2.941176 | 0.04076096 |
| Anaphase-Promoting Complex-Dependent Catabolic Process (GO:0031145) | 9.52381 | 0.04167244 |
| Regulation of Transcription Regulatory Region DNA Binding (GO:2000677) | 9.52381 | 0.04167244 |
| Intrinsic Apoptotic Signaling Pathway in Response to DNA Damage by P53 Class Mediator (GO:0042771) | 9.52381 | 0.04167244 |
| Myoblast Differentiation (GO:0045445) | 9.52381 | 0.04167244 |
| Negative Regulation of ERBB Signaling Pathway (GO:1901185) | 9.52381 | 0.04167244 |
| Negative Regulation of Signal Transduction by P53 Class Mediator (GO:1901797) | 9.52381 | 0.04167244 |
| Negative Regulation of Telomere Maintenance via Telomerase (GO:0032211) | 9.52381 | 0.04167244 |
| Positive Regulation of Extracellular Matrix Organization (GO:1903055) | 9.52381 | 0.04167244 |
| - Reg of Transmembrane Receptor Prot Serine/Threonine Kinase Sgnlng Pway (GO:0090101) | 4.098361 | 0.04199931 |
| + Reg of Transmembrane Receptor Prot Serine/Threonine Kinase Sgnlng Pway (GO:0090100) | 4.761905 | 0.04202576 |
| RNA Methylation (GO:0001510) | 6 | 0.04269512 |
| Negative Regulation of Protein Binding (GO:0032091) | 6 | 0.04269512 |
| Nucleocytoplasmic Transport (GO:0006913) | 6 | 0.04269512 |
| Negative Regulation of Intracellular Signal Transduction (GO:1902532) | 3.030303 | 0.04339706 |
| Positive Regulation of Cell Differentiation (GO:0045597) | 3.030303 | 0.04339706 |
| Ribosomal Small Subunit Biogenesis (GO:0042274) | 4.705882 | 0.04358273 |
| Regulation of Mitotic Cell Cycle (GO:0007346) | 4.032258 | 0.04451513 |
| Regulation of Epidermal Growth Factor Receptor Signaling Pathway (GO:0042058) | 5.882353 | 0.04485898 |
| Epithelial Structure Maintenance (GO:0010669) | 9.090909 | 0.04537907 |
| Peptidyl-Tyrosine Dephosphorylation (GO:0035335) | 9.090909 | 0.04537907 |

| | | |
|---|---|---|
| Positive Regulation of p38MAPK Cascade (GO:1900745) | 9.090909 | 0.04537907 |
| Receptor Metabolic Process (GO:0043112) | 9.090909 | 0.04537907 |
| Protein Modification Process (GO:0036211) | 2.568807 | 0.04665417 |
| Glycoprotein Metabolic Process (GO:0009100) | 5.769231 | 0.04707639 |
| Regulatory ncRNA-mediated Post-Transcriptional Gene Silencing (GO:0035194) | 8.695652 | 0.04920214 |
| Positive Regulation of DNA Replication (GO:0045740) | 8.695652 | 0.04920214 |
| Regulation of Cholesterol Biosynthetic Process (GO:0045540) | 8.695652 | 0.04920214 |
| Regulation of Focal Adhesion Assembly (GO:0051893) | 5.660377 | 0.04934689 |
| Regulation of Microtubule Polymerization (GO:0031113) | 5.660377 | 0.04934689 |
| Nucleotide-Excision Repair (GO:0006289) | 5.660377 | 0.04934689 |

**Supplementary Table 3:** **List of antibodies used for**

**A. NEED-seq**

| | |
|---|---|
| IgG rabbit | CST # 2729S |
| H3 | CST # 4499S |
| H3K9ac | CST # 9649S |
| H3K4me1 | CST # 5326S |
| H3K4me2 | CST # 9725S |
| LSD1 | CST # 2184S |
| HDAC1 | CST # 34589S |
| RCOR1 | Abcam # 183711 |
| Jun B | CST # 3753 |
| RUNX | Abcam # 92336 |

**B. Western blot**

| | |
|---|---|
| H3 | CST # 4499S |
| H3K9ac | CST # 9649S |
| H3K4me1 | CST # 5326S |
| H3K4me2 | CST # 9725S |
| LSD1 | CST # 2184S |
| HDAC1 | Abcam # ab7028 |
| RCOR1 | Abcam # 183711 |
| Jun B | CST # 3753 |
| RUNX | Abcam # 92336 |

**Supplementary Material 1: Complete description of NicEL Stages and Model Architecture**

**1. Introduction**

It's the method we're using for object detection in our microscopy images. Its primary design advantage is its robust handling of densely packed cells, mitigating the risks of over-segmentation (accidental merging) or under-segmentation (ignoring cells) by utilizing a **star-convex polygon** representation rather than conventional bounding boxes

**2. Model Architecture**

At its heart, the StarDist model is a U-Net. This is a type of Convolutional Neural Network (CNN) that's incredibly good at image-to-image tasks, which is exactly what segmentation is.

- **Encoder-Decoder Structure:** Think of a U-Net as having two parts:
    1. **Encoder (Contracting Path):** This is the "down" part of the "U". It's a standard CNN setup that uses convolutional and max-pooling layers to shrink the image down. As it shrinks, it captures the "what" (i.e., "that's a cell") but starts to lose the "where."
    2. **Decoder (Expanding Path):** This is the "up" part of the "U". It takes the compressed info from the encoder and progressively blows it back up to the original image size.
- **Skip Connections:** This is the magic of the U-Net. It adds "shortcuts" that connect layers from the encoder (the "down" part) directly across to the corresponding layers in the decoder (the "up" part). This lets the model combine the deep, abstract "what" information with the high-resolution "where" information. It's why U-Nets are so good at drawing precise outlines.

This U-Net backbone is set up with two different output heads, which are trained together to predict the two things StarDist cares about.

**3. Model Prediction Outputs**

For *every single pixel* in the input image, the model spits out two separate predictions:

1. **Object Probability (*d*):**
    - This is a one-channel map showing the probability that a pixel is part of *any* object.
    - Here's the key: this is **not** just a simple "cell vs. background" map. The "ground truth" it's trained on is the *normalized distance to the nearest background pixel*.
    - This is a clever design. It means pixels right in the *center* of a cell get a high probability (close to 1.0), while pixels near the edge get a low probability. This is super useful later because it gives us a clean way to find the best "center" pixel for each cell.
2. **Radial Distances ($\{r^k\}$):**
    - This is a multi-channel map. If the model is set to use n=32 radial directions (like 32 spokes on a wheel), this map will have 32 channels.

- For any given pixel, each channel k predicts the *Euclidean distance* from that pixel straight out to the cell's boundary along that $k_{th}$ direction.
- If a pixel is in the background, its distances are ignored during training.

So, for a 32-direction model, the U-Net is outputting 1 + 32 = 33 channels for every pixel.

**4. Initial Training Methodology**

**Ground Truth Generation**

The model learns from "instance segmentation masks" (where every cell has a unique ID). The training code takes these masks and, *on-the-fly*, generates the "answers" for its two predictions:

1. It calculates the distance-to-background probability map for every cell.
2. For every pixel inside a cell, it calculates the 32 radial distances to that cell's boundary.

**Loss Function**

The model's "loss" (what it's trying to minimize) is a combination of two parts:

1. **Object Probability Loss:** A standard **Binary Cross-Entropy** loss. This just pushes the model to get the probability map right (i.e., learn to find the cell centers).
2. **Radial Distance Loss:** A **Mean Absolute Error (MAE)** loss, which compares the predicted 32 distances to the real ones. But—and this is another clever part—this loss is *weighted* by the ground-truth probability.
   - **Background pixels** have a probability of 0, so their (garbage) distance predictions contribute *zero* to the loss.
   - **Pixels near the cell center** have a high probability, so their distance predictions are *heavily weighted*.
   - **Pixels near the boundary** have a low probability, so their predictions matter less.

This whole setup forces the model to get good at predicting distances *from the center of the cell*, which is exactly where we'll be making predictions from later.

**5. Inference and Non-Maximum Suppression (NMS)**

When we run the model on a new image (this is "inference"), it's a multi-step process to get from the raw predictions to the final masks:

1. **Dense Prediction:** The model runs, spitting out its 1-channel probability map and n-channel distance map.
2. **Candidate Generation:** We take the probability map and set a threshold (e.g., show me all pixels with a probability > 0.5). Every pixel that passes is now a "candidate" that will try to draw a polygon.
3. **Polygon Creation:** Each candidate pixel uses its n predicted distances to instantly "draw" a star-convex polygon. This creates a *massive* mess of thousands of overlapping polygons.

4. **Non-Maximum Suppression (NMS):** This is the cleanup step. It's an algorithm that filters out all the duplicates.
   - It finds the candidate polygon with the *highest object probability* (remember, this is our "center" pixel). This one is kept.
   - It then finds all other polygons that overlap with this one (using an **Intersection over Union, or IoU,** threshold).
   - All these high-overlap "duplicates" are *suppressed* (thrown away).
   - The algorithm repeats, finding the next-highest-probability polygon that hasn't been suppressed yet, and so on.

The final output is just the list of polygons that "won" the NMS process.

#### 6. Evaluation Metrics

How do we know if it's working? We use the standard **Intersection over Union (IoU)**, which is just a score of how much a predicted polygon ($I_{pred}$) and a ground-truth mask ($I_{gt}$) overlap.

- **IoU:** $\frac{I_{pred} \cap I_{gt}}{I_{pred} \cup I_{gt}}$ (A score of 1.0 is a perfect match).
- A **True Positive (TP)** is a prediction that overlaps a ground-truth mask with an IoU *above* a threshold (e.g., $\tau = 0.5$).
- A **False Positive (FP)** is a predicted polygon that doesn't match any ground-truth mask.
- A **False Negative (FN)** is a ground-truth mask that didn't get matched by any prediction.
- **Average Precision (AP):** This is the main score we look at. It's a single number calculated from the TPs, FPs, and FNs that summarizes how good the model is. We usually check this score at a few different IoU thresholds to get the full story.

#### 7. Fine-Tuning Workflow for New Data

Okay, this is the main part for us. Instead of training from scratch (which would take *ages* and need thousands of images), we're doing **fine-tuning**. This just means we take a pre-trained model and "nudge" it to be better at our specific images.

##### A. Data Annotation for Fine-Tuning

This is the most important and time-consuming part. The model *cannot* learn from raw images or simple (binary) cell vs. background masks.

##### Required Annotation: Instance Segmentation Masks

- We have to create a label image (e.g., a 16-bit TIFF) for each of our raw images.
- In this label image, all background pixels **must** be **0**.
- Every pixel for the *first* cell gets a value of **1**.
- Every pixel for the *second* cell gets a value of **2**.
- ...and so on. Every single cell needs its own unique ID.

This is the only way the training process can figure out which pixels belong to which object to calculate all those ground-truth distances and probabilities.

**B. The Fine-Tuning Process**

1. **Load Pre-trained Model:** We start by loading an official StarDist model that was trained on a big, general dataset (like 2D_versatile_fluo for our fluorescent images). This model already has a great "idea" of what a nucleus looks like
2. **Prepare Custom Data:** We load our set of ~100 custom images and their matching instance masks
3. **Split Data:** We'll split our data into a *training set* and a *validation set*
4. **Data Augmentation:** Since 100 images are a *tiny* dataset, we have to use heavy data augmentation. This means the training code randomly messes with the images *as it's training* to create an almost infinite supply of "new" examples. This prevents the model from just memorizing our 90 training images. Our pipeline includes:
   - Separating multi-channel images (if any) into individual black and white images.
   - Random rotations
   - Flipping (horizontal/vertical)
   - Random scaling (zooming)
   - Random brightness and contrast adjustments
5. **Re-Train the Model:** This is the fine-tuning step.
   - To answer your question: we are re-training the *entire network*, not just the last few layers
   - However, we do it with a very **low learning rate**. This is the "nudging" part you mentioned. The model's weights are already 95% of the way there; we're just gently adjusting *all* of them to be a little bit better at recognizing the specific shapes, sizes, and brightness of *our* cells
   - This is way faster than training from scratch and gives a much better result for small datasets.

**8. A Typical Deployment Workflow**

Once we have our fine-tuned model, here's how we'll use it in our pipeline:

1. **Load Model:** The final fine-tuned model is saved as a folder (containing a config.json file and a weight.h5 file). We load this model in our script.
2. **Read Image:** We read one of our raw fluorescent image files (e.g., a .czi file).
3. **Pre-process:** The image file is converted into a standard NumPy array. We also have to "normalize" it (i.e., scale its pixel values) in the same *way* the model was trained.
4. **Predict:** We pass the normalized image array to the model.predict_instances() function. This function runs all the steps from Section 5 (dense prediction, NMS, etc.) internally.
5. **Get Masks:** The function returns our final set of clean, non-overlapping masks. We can then use these masks for our downstream analysis.
6. Detect Nuclei
7. Calculate ACI

To quantify the degree of chromatin accessibility within individual nuclei, we defined the **Accessible Chromatin Index (ACI)**, a metric representing the ratio of peak open chromatin signal to the baseline nuclear intensity. For each segmented nuclear volume, raw multi-channel intensity data $I \in \mathbb{R}^{H \times W \times C}$ was masked to isolate voxel-specific signals. Let $V_{ch1}$ and $V_{ch2}$ represent the sets of non-zero intensity values for Channel 1 (target open chromatin marker) and Channel 2 (reference nuclear marker), respectively, extracted from the i-th nuclear mask. The ACI for each nucleus was calculated as the quotient of the maximum intensity in the target channel and the minimum intensity in the reference channel:

$$ACI_i = \frac{max(V_{ch1,i})}{min(V_{ch2,i})}$$

To ensure robust signal-to-noise ratios and prevent computational artifacts, the index was only computed for nuclei satisfying the following criteria: $V_{ch1}, V_{ch2}$=∅ and $max(V_{ch1})>0$. Nuclei failing to meet these thresholds, often due to edge-case segmentation or signal voids, were excluded from the analysis and assigned a null value (NaN). This normalization approach allows for the characterization of chromatin condensation states across heterogeneous cell populations by anchoring the open chromatin signal against the nuclear interior's minimum intensity baseline.
